# Optimizing selection of restriction enzymes for complexity-reduced genome sequencing in plants

**DOI:** 10.1101/2025.09.21.677652

**Authors:** Kenta Shirasawa, Sachiko Shirasawa

**Author notes:** **Correspondence to:** Kenta Shirasawa, 2-6-7 Kazusa-Kamatari, Kisarazu, Chiba 292-0818, Japan.

## Abstract

**Background:** Reduced-representation sequencing, e.g., restriction-site-associated DNA sequencing and genotyping by sequencing, is a powerful and cost-effective method to detect polymorphism and to genotype pools of individuals. However, restriction enzymes employed to the analyses were often chosen in basis on intuition and precedents.

**Results:** We propose to apply an *in silico* analysis to predict gene concentrations in restriction fragments, which are templates of complexity-reduced genome libraries. It also predicts fragment lengths and physical positions in plant genomes. The *in silico* analysis was verified using actual amplified fragment length polymorphism patterns. The genome-scale differences in the distributions of restriction fragments from methylation-sensitive and resistant enzymes accounted for marker clusters on genetic maps, commonly reported in linkage maps of plants. The *in silico* analysis, using three combinations of enzymes across four model plants, indicated that the combination of MspI and PstI is remarkably informative for reduced-representation sequencing in terms of fragment length, distribution in euchromatic regions, and gene enrichment.

**Conclusions:** Application of our *in silico* restriction analysis provides useful information for optimizing the selection of restriction enzymes in empirical reduced-representation sequencing of genomes of several organisms including not only plants but also animals.

## Background

DNA marker technology has improved in the last few decades [1]. Amplified fragment length polymorphism (AFLP) markers have been especially popular in plant genetics due to their high reproducibility and because they do not require reference genome sequences [2]. In AFLP analysis, genomic DNA is digested with two restriction enzymes, ligated with oligonucleotide adaptors, and used as templates for adaptor-ligation PCR using oligonucleotides that are specific to the adaptor sequences [3]. The complete genome sequence for *Arabidopsis thaliana* has been released [4]. This whole-genome sequence data enabled the development of large numbers of single nucleotide polymorphism (SNP) and simple sequence repeat (SSR) markers. These have replaced AFLP markers because they require less time and fewer resources to develop.

In this decade, massive parallel DNA sequencing technologies, known as next-generation sequencing (NGS) technologies, have been available in plant genetics and genomics. NGS produces mega-, giga-, or tera-base scales of sequence data in a single experiment, which is often enough to generate reference genome sequences even in “non-model” plants, including important crops [5–7]. Moreover, it replaces SNP markers as a molecular genetics tool because whole-genome shotgun sequencing or reduced-representation sequencing for multiplex samples, e.g., mapping or breeding populations, can be archived using NGSs [8]. Reduced-representation sequencing methods include restriction site-associated DNA sequencing (RAD-seq) [9] and genotyping by sequencing (GBS) [10]. For both methods, procedures similar to AFLP analysis have been employed to construct sequencing libraries: genomic DNA is digested with restriction enzymes, ligated with adaptors, and amplified by adaptor-ligation PCR for sequencing. Alternative methods of RAD-seq and GBS, using a couple of enzymes, have also been proposed to reduce the time, labor, and cost of library preparation [11], and to reduce complexity [12].

In linkage analysis, however, AFLP marker loci tend to be clustered in each linkage group when methylation-resistant enzymes, e.g., MseI and EcoRI, are employed [13]. The clusters correspond to repetitive sequence regions of heterochromatin suppressing chromosome recombination [14–16]. Because hypermethylation in the heterochromatin inhibits the action of methylation-sensitive enzymes, e.g., MspI and PstI, they are often used for library construction to avoid marker clustering [15]. The use of methylation-sensitive enzymes has improved GBS [10]. In addition, a computational approach to identify distributions of restriction sites from published genome sequences, called *in silico* AFLP or *in silico* restriction analysis, has been developed [17, 18]. The *in silico* AFLP analysis suggests that biases in GC content, and in the density of polymorphisms and repeated sequences, are responsible for marker clustering. Therefore, the *in silico* restriction analysis optimizes restriction enzymes prior to RAD-seq and GBS analysis [11, 19].

Genome data provide biologically meaningful information, e.g., the number, structure, and density of genes, as well as the distribution of restriction sites. Caballero *et al*. [17] report the proportion of AFLP markers physically linked to genes. However, few studies report restriction fragment lengths and gene content in the reduced-represented fragment libraries. The former is technically important for determining lengths of sequence reads from NGS data, while the latter provides biologically useful information for trait mapping by quantitative trail loci (QTL) analysis and genome-wide association study (GWAS).

In this study, we applied an *in silico* restriction analysis to optimize selection of restriction enzymes for construction of complexity-reduced genome libraries for RAD-seq and GBS analyses. We established the *in silico* restriction analysis method and verified it using actual banding patterns in a previous AFLP study [20]. Next, the distributions of restriction fragments from methylation-sensitive and resistant enzymes were investigated using tomato genome sequence data. Finally, we compared three combinations of enzymes across four model plant species and demonstrated that the *in silico* restriction analysis provides useful information for empirical reduced-representation sequencing analysis.

## Results

### *In silico* restriction analysis and comparison with AFLP gel images in rice

Since the initial release by the International Rice Genome Sequencing Project [21], the rice “Nipponbare” genome sequence has been improved to cover approximately 97% of the genome with an error rate of <0.15 per 10,000 nucleotides [22]. We used the high quality reference genome sequence data to validate the accuracy of the *in silico* restriction analysis developed in this study (see the Methods section) by comparison with actual AFLP gel images of rice “Nipponbare” from our previous study [20].

The rice pseudomolecule corresponding to the 12 chromosomes was *in silico* digested with MseI and EcoRI, which are employed in our previous study [20], into 2,326,086 restriction fragments. The fragments were comprised of 160,831 sequences with EcoRI and MseI sites at both ends (MseI-EcoRI fragments), 7,791 with EcoRI sites on both ends, 2,157,440 with MseI sites on both ends, and 24 chromosome-end fragments. The sizes of the 160,831 MseI-EcoRI fragments, which are potential targets for AFLP analysis, were distributed from 4 to 5,649 bp, with an average of 132 bp and a median of 239 bp.

Subsequently, 102,841 MseI-EcoRI fragments with lengths of 72 to 974 bp, or 100 to 1,000 bp after adaptor-ligation PCR [20], were selected for further analysis (Additional file 1). The 102,841 fragments (mentioned hereafter as amplified fragments) were evenly distributed throughout the rice genome (373,245,519 bp) with 276 fragments/1 Mb (=1 fragment/3,623 bp) on average. Of them, 34,401 (33%) amplified fragments were expected to include 14,884 (39%) of 37,869 gene loci predicted from approximately 30% (113,537,097 bp) of the rice genome [23].

Of the 102,841 amplified sequences, overrepresented fragments with identical lengths and 3 bp flanking sequences of restriction sites on each end were identified. The most frequent fragments comprised 197 bp with MseI-ATT and EcoRI-CTG ends, represented 430 times. Fragments with MseI-AGT and EcoRI-CCA ends were 135 bp long and were represented 423 times. In total, 17, 44, and 324 overrepresented fragments were predicted >100, >50, and >10 times, respectively. On the other hand, when the 102,841 amplified fragments were classified into 4,096 groups based on the selective nucleotides for the EcoRI and MseI sites, substantial bias was observed, i.e., from 4.6 fragments with six CGs in the six bases to 90.4 fragments without CGs (Additional files 1 and 2).

The banding pattern of the amplified fragments from the *in silico* analysis were compared to the pattern in actual AFLP data from our previous study [20]. One hundred and forty-four primer pairs (combinations of 12 primers with three selective nucleotides for each enzyme: ACG, AGC, ATG, CAA, CAC, CAG, CAT, CGT, GCA, GTA, TAC, and TGC) were investigated. In the *in silico* restriction analysis, 3,454 restriction fragments were generated from the 144 primer pairs, but 3,116 amplified fragments ranging from 100 to 1000 bp were predicted (Additional files 1 and 3). Comparing *in silico* and experimental data, 2,571 of the 3,116 predicted fragments from the *in silico* AFLP analysis were detected in the gel images, and DNA bands with high molecular weight (>500 bp) were rarely represented (171 of 484) (Additional file 3). All nucleotide sequences from the 13 bands cloned in the previous study [20] matched *in silico* predictions. These results suggest that the *in silico* restriction analysis can accurately predict sizes, numbers, and distributions of restriction fragments in genomes.

### *In silico* restriction analysis with methylation-resistant and sensitive restriction enzymes in tomato

We compared the effects of combinations of restriction enzymes of MseI and either different methylation sensitivities, EcoRI or PstI, on the complexity-reduced genome. These combinations of enzymes have been used in AFLP analysis in tomato [15]. Tomato has a complex genome compared with rice. It is approximately 900 Mb, and >70% (569 Mb) was identified as pericentromeric heterochromatin, with a high concentration of repetitive sequences such as retrotransposons [24]. On the other hand, 62% of tomato genes are located in the euchromatin (191 Mb), where the chromosome recombination rate is higher [25].

The pseudomolecules of the tomato genome, excluding SL2.40ch00, which are not assigned to the chromosomes, were totally digested with the two combinations of restriction enzymes, i.e., MseI and EcoRI and MseI and PstI, into 7.4 million (M) and 7.2 M fragments, respectively. Of them, 444,837 MseI-EcoRI and 150,965 MseI-PstI fragments were selected. The sizes of the MseI-EcoRI and MseI-PstI fragments ranged from 4 bp to 27,509 bp (mean: 131 bp; median: 76 bp) and from 2 bp to 14,845 bp (mean: 206 bp; median: 118 bp), respectively (Figure 1).

**Figure 1.**
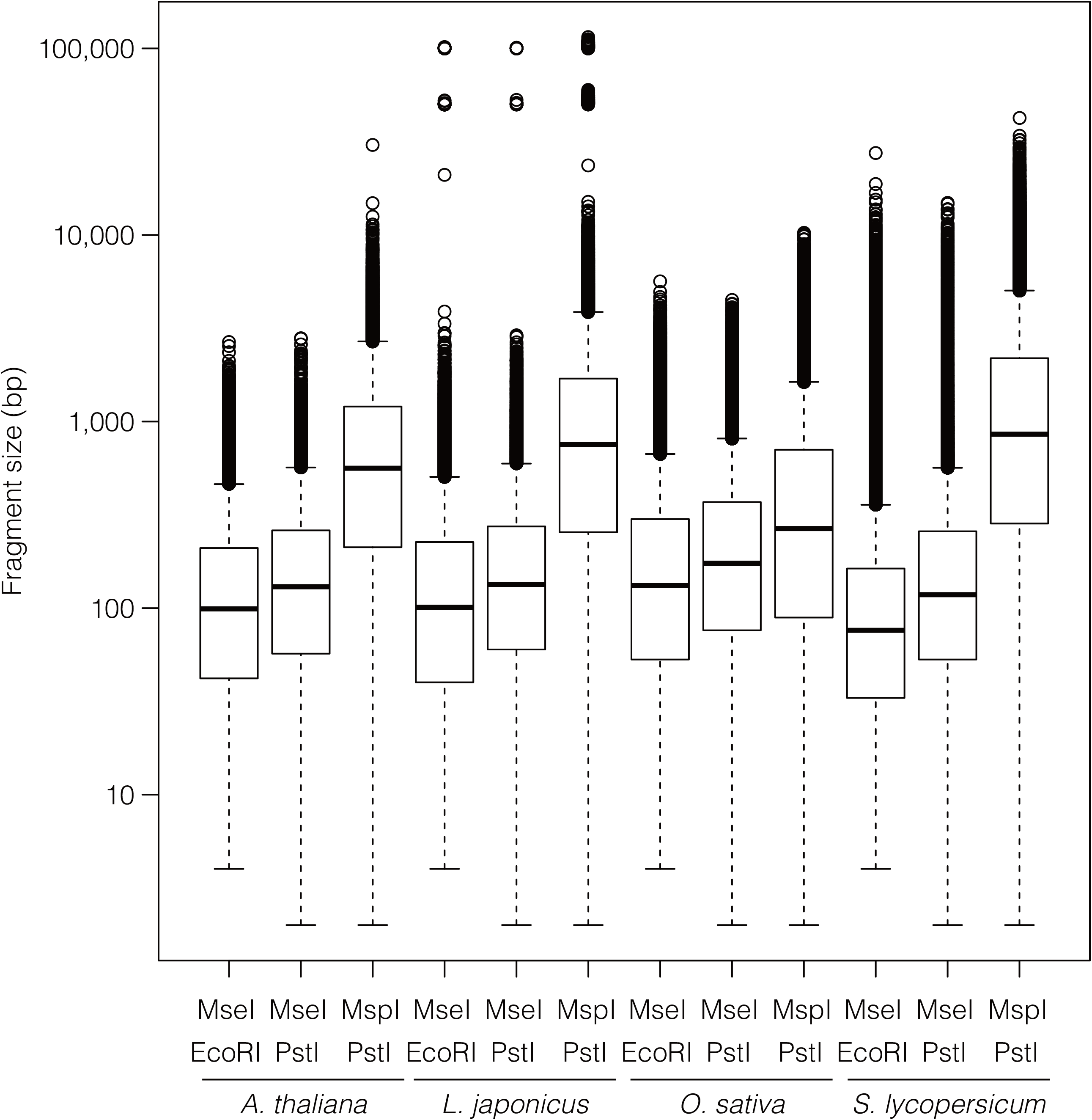
Quantile boxplots for the distributions of restriction fragment sizes from three enzyme combinations over the four plant species. The Y-axis is shown on a logarithmic scale. The bottom and top edges of the boxes are the 25th and 75th quantiles, and horizontal lines in the boxes indicate the median. The whiskers delimit ± 1.5 × interquartile range.

Restriction fragments with lengths from 100 bp to 1,000 bp, which are potential targets for the complexity-reduced genome libraries, were selected for further analysis. For MseI-EcoRI and MseI-PstI, this reduced the sample sizes to 179,814, and 83,276, respectively (Additional files 4 and 5). In accordance with the hetero- and euchromatin portions of the tomato genome [25], 44,018 (24%) MseI-EcoRI and 27,190 (33%) MseI-PstI fragments were found in euchromatic regions (191 Mb), while the other 135,796 (76%) and 56,086 (67%) fragments were derived from heterochromatin regions (569 Mb) (Additional files 6 and 7). There is a higher density of restriction fragments in euchromatin than in heterochromatin for MseI-PstI (99 in heterochromatin vs. 142 in euchromatin per 1 kb window), but not for MseI-EcoRI (239 in heterochromatin vs. 231 in euchromatin per 1 kb window).

### Distributions of restriction fragments in the genomes of model plants

The relationship between enzymes, sizes of restriction fragments, and genome positions was investigated. While more than 50 plant genomes has been released [5–7], in this study, *Arabidopsis thaliana* and *Lotus japonicus*, representing Brassicaceae and Fabaceae, respectively, were chosen in addition to rice (Poaceae) and tomato (Solanaceae) as models of the *in silico* restriction analysis. The genome sequences of the four species were *in silico* digested with each of the three pairs of enzymes, i.e., MseI-EcoRI, MseI-PstI, or MspI-PstI, which represent differences not only in methylation sensitivity, but also in the GC content in the enzyme recognition sites (see the Methods section).

### Solanum lycopersicum

The *in silico* digestion of the tomato genome yielded 180 thousand (k) MseI-EcoRI and 83 k MseI-PstI 100–1000 bp fragments, as described above. The *in silico* analysis generated 500 k MspI-PstI fragments (mean: 1,733 bp; median: 856 bp), and 51 k with lengths of 100–1000 bp (Figure 1, Additional file 8). The MspI-PstI fragments showed higher coverage in the distal ends of the chromosomes compared with the centromeric regions (Additional file 9).

Thirty-six thousand (20%), 29 k (35%), and 17 k (32%) fragments, respectively, were associated with predicted genes from the tomato genome (Table 1). Out of 33,840 predicted genes in the tomato genome, 21,087 and 12,753 were identified in the eu- and heterochromatin regions, respectively (Additional file 10). In non-overlapping windows of 1 Mb, the correlation between the numbers of the restriction fragments and genes was investigated. Positive correlations were observed for MseI-PstI (R^2^=0.540, *P*<2.20E-16) and MspI-PstI (R^2^=0.261, *P*<2.20E-16), but a weak negative correlation was observed for MseI-EcoRI (R^2^=0.010, *P*=4.21E-03).

**Table 1.**
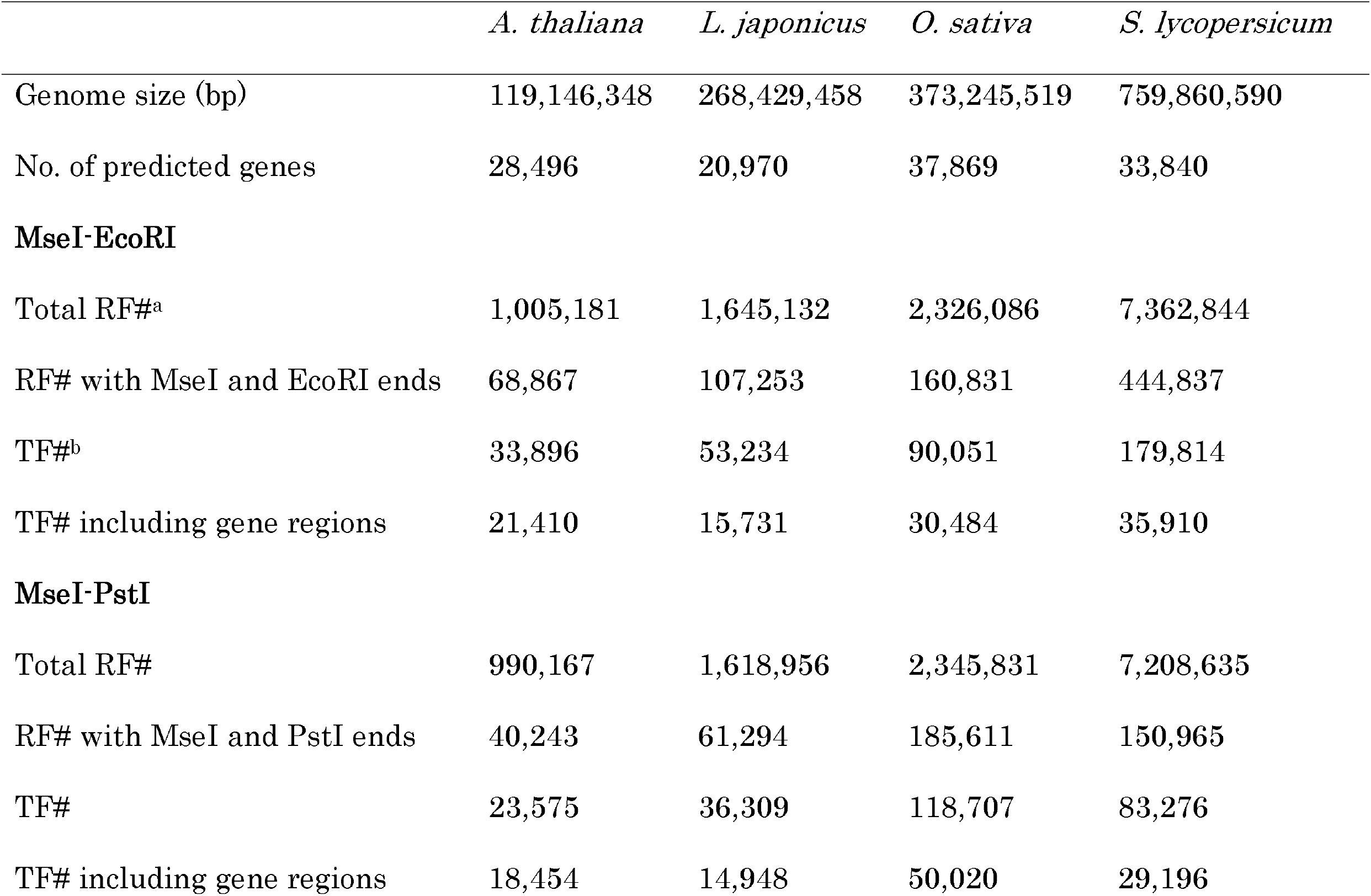
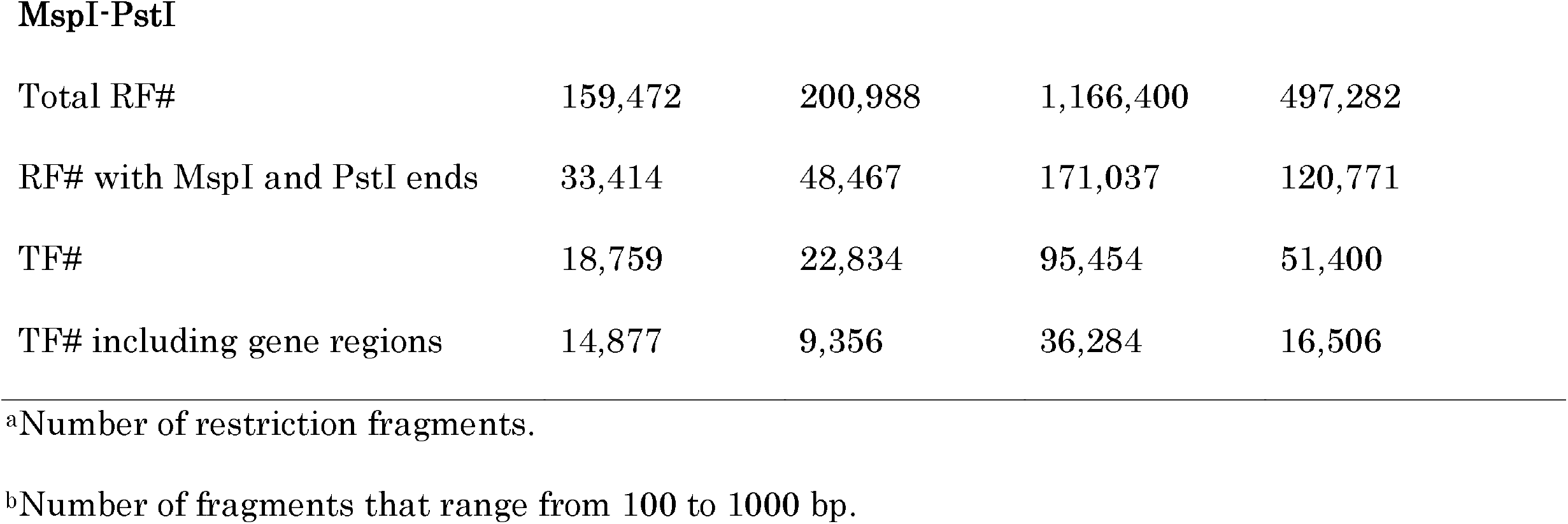
Numbers of restriction fragments from the genomes of four plant species.

|  | *A. thaliana* | *L. japonicus* | *O. sativa* | *S. lycopersicum* |
| --- | --- | --- | --- | --- |
| Genome size (bp) | 119,146,348 | 268,429,458 | 373,245,519 | 759,860,590 |
| No. of predicted genes | 28,496 | 20,970 | 37,869 | 33,840 |
| **MseI-EcoRI** |  |  |  |  |
| Total RF#<sup>a</sup> | 1,005,181 | 1,645,132 | 2,326,086 | 7,362,844 |
| RF# with MseI and EcoRI ends | 68,867 | 107,253 | 160,831 | 444,837 |
| TF#<sup>b</sup> | 33,896 | 53,234 | 90,051 | 179,814 |
| TF# including gene regions | 21,410 | 15,731 | 30,484 | 35,910 |
| **MseI-PstI** |  |  |  |  |
| Total RF# | 990,167 | 1,618,956 | 2,345,831 | 7,208,635 |
| RF# with MseI and PstI ends | 40,243 | 61,294 | 185,611 | 150,965 |
| TF# | 23,575 | 36,309 | 118,707 | 83,276 |
| TF# including gene regions | 18,454 | 14,948 | 50,020 | 29,196 |
| **MspI-PstI** |  |  |  |  |
| Total RF# | 159,472 | 200,988 | 1,166,400 | 497,282 |
| RF# with MspI and PstI ends | 33,414 | 48,467 | 171,037 | 120,771 |
| TF# | 18,759 | 22,834 | 95,454 | 51,400 |
| TF# including gene regions | 14,877 | 9,356 | 36,284 | 16,506 |

<sup>a</sup>Number of restriction fragments.
<sup>b</sup>Number of fragments that range from 100 to 1000 bp.

### Oryza sativa

In the *in silico* analysis of the rice genome, 90 k MseI-EcoRI fragments (100–1000 bp in length), evenly distributed throughout the rice genome (Additional file 11), were obtained (Table 1, Additional file 12). On the other hand, the analysis generated 186 k MseI-PstI (mean: 281 bp; median: 174 bp) and 171 k MspI-PstI fragments (mean: 540 bp; median: 267 bp), and 119 k and 95 k with lengths of 100–1000 bp (Figure 1, Additional files 13 and 14). The MseI-PstI and MspI-PstI fragments showed slightly higher coverage in the distal ends of the chromosomes compared with the centromeric regions (Additional files 15 and 16). Thirty thousand (34%) MseI-EcoRI, 50 k (42%) MseI-PstI, and 36 k (38%) MspI-PstI fragments overlapped the predicted gene regions (Table 1). The 37,869 genes from the rice genome were distributed unevenly over the genome, with low and high density in centromeric regions and distal ends, respectively (Additional file 17). Significant correlations between the numbers of genes and fragments within non-overlapping windows of 1 Mb were observed for MseI-PstI (R^2^=0.620, *P*<2.20E-16) and MspI-PstI (R^2^=0.378, *P*<2.20E-16), but not for MseI-EcoRI (R^2^=0.003, *P*=1.50E-01) (Figure 2).

**Figure 2.**
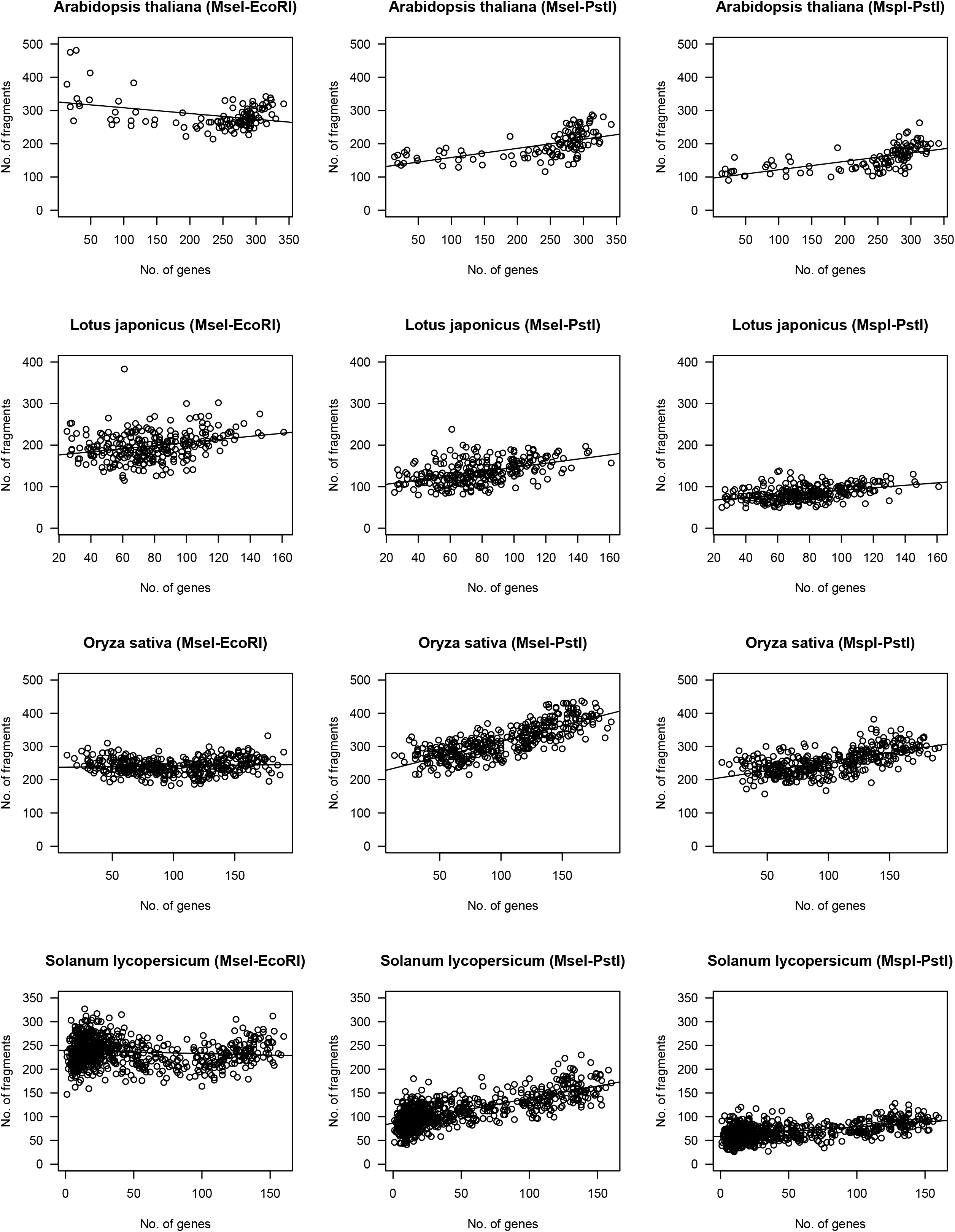
Relationship between numbers of genes and those of the restriction fragments within non-overlapping windows of 1 kb.

### Arabidopsis thaliana

The *Arabidopsis* genome was *in silico* digested with each combination of enzymes. The genome was cleaved to 69 k MseI-EcoRI (mean: 160 bp; median: 99 bp), 24 k MseI-PstI (mean: 193 bp; median: 130 bp), and 19 k MspI-PstI fragments (mean: 897 bp; median: 562 bp) (Figure 1, Table 1). This included 34 k, 24 k, and 19 k target fragments for MseI-EcoRI, MseI-PstI, and MspI-PstI, respectively (Additional files 18, 19, and 20). Similar to tomato and rice, the MseI-PstI and MspI-PstI fragments were located on the distal regions of chromosomes with higher coverage than the centromeric regions, while MseI-EcoRI fragments were evenly distributed over the genome (Additional files 21, 22, and 23). Gene sequences were found in 21 k (63%) MseI-EcoRI fragments, 18 k (78%) MseI-PstI fragments, and 15 k (79%) MspI-PstI fragments (Table 1). The predicted genes were located mainly on the distal ends of chromosomes and were underrepresented on the centromeres (Additional file 24). Positive correlations between the densities of the genes and fragments were observed for MseI-PstI (R^2^=0.350, *P*=1.57E-12) and MspI-PstI (R^2^=0.387, *P*=5.48E-14), but the correlation was negative for MseI-EcoRI (R^2^=0.121, *P*=7.90E-05) (Figure 2).

### Lotus japonicus

The *in silico* restriction analysis was applied to the *L. japonicus* genome, excluding Chr0. The numbers of digested fragments were 1.6 M (mean: 270 bp; median: 101 bp), 1.6 M (mean: 311 bp; median: 134 bp), and 201 k (mean: 1750 bp; median: 754 bp) using MseI-EcoRI, MseI-PstI, and MspI-PstI, respectively. The fragment numbers were reduced to 53 k, 36 k, and 23 k after filtering those that were not 100–1000 bp in length (Additional files 25, 26, and 27). The fragments from all three combinations of restriction enzymes evenly covered the genomes, and 16 k (30%) MseI-EcoRI, 15 k (41%) MseI-PstI, and 9 k (41%)

MspI-PstI fragments contained gene regions (Additional files 28, 29, and 30). On the other hand, the predicted genes in the *Lotus* genome were distributed with high and low coverage in distal and centromeric chromosome regions, respectively (Additional file 31). Even though the correlations between the densities of genes and fragments were weaker than those observed in other species, they were all significantly positive: R^2^=0.073 (*P*=4.58E-06) for MseI-EcoRI, R^2^=0.211 (*P*=1.58E-15) for MseI-PstI, and R^2^=0.188 (*P*=8.32E-14) for MspI-PstI (Figure 2).

For all four species, there were fewer MspI-PstI fragments than MseI-EcoRI and MseI-PstI fragments, and MspI-PstI fragments were the longest (Figure 1, Table 1). The mean densities of the fragments in the genomes ranged from 3 kb of MseI-PstI in rice to 21 kb of MseI-EcoRI in tomato (Table 1). In addition, the MseI-PstI and MspI-PstI fragments were preferentially generated from gene rich-regions, in contrast to MseI-EcoRI (Table 1).

## Discussion

We applied an *in silico* restriction analysis to the genomes of the four model plants. The combination of enzymes influenced fragment size, genomic position, and concentration of genes in the fragments. Marker development and linkage map construction using whole-genome or reduced-representation sequencing are cost-effective. The enzymes used should be optimized by *in silico* analysis prior to trial experiments to further reduce cost.

The *in silico* restriction method was evaluated with rice AFLP data [20]. As expected, predictions of the *in silico* analysis matched the actual experimental data (Additional files 1 and 3), even though fragments of ≥500 bp are sometimes lost in the gel images (Additional file 3). This could be explained by preferential amplifications of short DNAs especially in highly multiplexed PCR. On the other hand, even the predicted bands of <500 bp were occasionally missed in the gel images, and unexpected bands were also observed (Additional file 3). We considered three possibilities that can account for these observations. First, cytoplasmic genomes, 135 and 491 kb for the rice plastid and mitochondrion, respectively, were not included in the analysis. Although both the plastid and mitochondrion genomes of rice have been sequenced [26, 27], they are reported to have many forms due to their intramolecule recombination [28], preventing accurate predictions using the *in silico* analysis. Next, it may be due to sequence variation within Nipponbare strains [22]. The Nipponbare strain sequenced by the International Rice Genome Sequencing Project [21] and analyzed by Shirasawa *et al*. [20] might be slightly different. Finally, it is possible that the rice reference genome is still not complete. Even in the latest version of the rice genome, Os-Nipponbare-Reference-IRGSP-1.0, which we used in this study, coverage is approximately 97% [22] and included 117 kb of uncharacterized bases, Ns. Despite these issues, however, our method was accurate. It can be applied to physical mapping of bands in AFLP analysis and restriction landmark genomic scanning (RLGS), as suggested by Peters *et al*. [18] and Rouillard *et al*. [29].

Clusters of AFLP markers in linkage maps have been observed in tomato [15], *Arabidopsis thaliana* [14], *Lotus japonicus* [16], and so on (referred in Caballero *et al*., [17]). Severe suppression of recombination in the heterochromatin accounts for this clustering [18]. The accumulation of marker loci is observed for the mapping of SSR markers, which are redundantly generated from the genome sequence data of tomato [20]. Peters *et al*. [18], moreover, points out that marker clustering can occur if polymorphism is higher in centromeric regions than that in euchromatic regions comprised of coding sequences. In addition, Caballero *et al*. [17] suggested that GC bias accounts for the marker clustering. In tomato, on the other hand, AFLP markers using a methylation-sensitive enzyme, PstI, are better than those using a methylation-insensitive enzyme, EcoRI, because PstI only recognizes non-methylated sites enriched in the euchromatin [15]. This might account for the results from complexity-reduced genome libraries, which use a methylation-sensitive enzyme, ApeKI [10]. Our observations also indicated than the MseI-PstI fragments were rich in genic regions in comparison with the MseI-EcoRI fragments, even though the current *in silico* analysis did not detect methylation differences. This tendency would be therefore remarkably robust in actual empirical analysis detectable methylated sites. We conclude that the marker clusters were due to the following: 1) the suppression of recombination in heterochromatin; 2) the bias of GC content in some regions of the genome; 3) the different methylation pattern in chromatin; 4) and the distributions of restriction sites over the genome.

In all four species, the lengths of the MspI-PstI fragments were longer than those digested with MseI-EcoRI and MseI-PstI (Figure 1). In standard RAD-seq and GBS [9, 10], the genome library is constructed with a single enzyme followed by random sharing. The sequencing of the library generates ultra-deep sequence reads from the restriction ends, but not from the other ends generated by random sharing. On the other hand, in double digest RAD-seq, ultra-deep sequence reads can be obtained from the both ends, because most restriction sites are conserved among individuals [11]. Therefore, paired-end sequencing technologies can be used for double digested fragments. The ultra-deep paired-end sequencing eliminates errors derived from sequencing and alignment due to the complexity of plant genomes, including repetitive sequences and polyploid genomes. Moreover, pair-end sequencing enables us to distinguish individuals from multiplexed samples by dual-indexed barcodes, which can decrease experimental costs. To make full use of the technology, long insert sizes of the RAD library would be desirable. Moreover, long and paired-end reads are useful for determining haplotype phases, which are necessary for linkage-disequilibrium based mapping strategies, e.g., GWAS.

The MspI-PstI and MseI-PstI fragments were enriched for genes (especially MspI-PstI), but MseI-EcoRI did not show a similar pattern of enrichment (Table 1), even though the numbers of fragments overlapping gene regions in the MspI-PstI and MseI-PstI fragments were less than that of MseI-EcoRI. The number of the fragments that include gene regions from either combination of enzymes might be, however, enough to tag any genes from the tested species genomes because densities ranged from 0.8 (=37,869/50,020) genes/1 fragment of the MseI-PstI in rice to 2.2 (=20,970/9,356) of the MspI-PstI in *L. japonicus* (Table 1). Caballero et al. [17] state that searching for AFLP markers closely linked to selected loci for specific traits is a difficult task, because only a small percentage of markers are expected to be close to any particular gene. For RAD-seq and GBS, on the other hand, huge amounts of sequence data covering the whole genome enhance the opportunity to find polymorphism within the targeted markers, or the genes themselves causing phenotypic variations. From the perspective of technical issues, restriction fragments from the gene-rich distal ends of chromosomes, unique sequences in the genome, would be beneficial to discover accurate SNPs in comparison with the gene-poor regions in the centromeres and pericentromeres, repetitive genome sequences.

## Conclusions

Genome sequence data for several plants, including important crops, have been published and are still accumulating, and it is true for animals as well [30]. Furthermore, many different restriction enzymes are available right now [31]. Using these genome data, restriction enzymes, and resequencing strategies such as RAD-seq and GBS, studies of genetic diversity within and between species can be improved. Applying the *in silico* analysis presented in this study to the organisms of interest together with available enzymes helps predict fragment size, genome location, and the proportion of fragments associated with genes prior to empirical reduced-representation sequencing analysis. This might increase the opportunity for RAD-seq and GBS to detect polymorphisms and mutations directly conferring phenotypic variations.

## Methods

### Data sources

The genome sequence data for *O. sativa* (Os-Nipponbare-Reference-IRGSP-1.0) [22], *S. lycopersicum* (SL2.40) [24], *L. japonicus* (Lj2.5) [32], and *A. thaliana* (TAIR10) [4] were obtained from the Rice Annotation Project Database [33], the Sol Genomics Network [34], the miyakogusa.jp [35], and TAIR [36], respectively. Gene annotation files in the Generic Feature Format were also downloaded from the same databases for the genome sequences, e.g., locus.gff of the IRGSP-1.0_representative_2013-04-24 for rice, ITAG2.3_gene_models.gff3 for tomato, Lj2.5_gene_models.gff3 for *L. japonicus*, and TAIR10_GFF3_genes.gff for *A. thaliana*, respectively. From each genome annotation file, gene sets were extracted when the description in the “type” of column 3 was “gene”.

### *In silico* restriction analysis

Three combinations of the restriction enzymes, e.g., MseI (recognition at site T↓TAA) and EcoRI (G↓AATTC), MseI and PstI (CTGCA↓G), and MspI (C↓CGG) and PstI, were used. The genome sequences were *in silico* treated with three combinations of the enzymes using shell scripts for Linux as: 1) The genome sequences were digested into restriction fragments at the points of the recognitions sites of the enzymes, 2) information on physical positions in the genome sequences of each fragment, sizes of each fragment, and 3-bp selective nucleotide sequences for each end were retained, and 3) the predicted genes in the genomes were assigned to the restriction fragments based on the physical positions. The banding patterns of the fragments were drawn with MapChart program (Voorrips 2002), in which the mobilities of the fragments were represented by converting with a formula of [1-(1/2)^size/100-1^]×1000, where *size* is the size of each fragment (100–1000 bp), to approximate to the actual gel images. The resultant banding patterns were manually compared with the gel images of the AFLP analysis from our previous study [20]. The correlation between the numbers of genes and restriction fragments was analyzed with the linear model in R program [37].

## List of abbreviations used

AFLP: Amplified fragment length polymorphism;
GBS: Genotyping by sequencing;
GWAS: Genome-wide association study;
NGS: Next-generation sequencing;
QTL: Quantitative trail loci;
RAD-seq: Restriction site-associated DNA sequencing;
RLGS: Restriction landmark genomic scanning;
SNP: Single nucleotide polymorphism;
SSR: Simple sequence repeat.

## Competing interests

The authors declare that they have no competing interests.

## Authors’ contributions

KS and SS designed this work, and analyzed the data. KS wrote the manuscript. The all authors read and approved the final manuscript.

## Acknowledgments

This work was supported by the Kazusa DNA Research Institute Foundation.

## Additional files

**Additional file 1 (XLSX) Supplementary Table S1** Genome positions of AFLP markers in the rice genome.

**Additional file 2 (XLSX) Supplementary Table S2** Numbers of fragments amplified with each primer combination for selective PCR in AFLP.

**Additional file 3 (PDF) Supplementary Figure S1** *In silico* AFLP and experimental data.

The left and right pictures are the *in silico* and actual experimental AFLP data, respectively. Primer combinations for MseI (M) and EcoRI (E) are shown on the tops of the pictures. Fragment sizes predicted in the *in silico* AFLP analysis are indicated by numbers, each of which corresponds to a single AFLP fragment. The left and right lanes in the actual images are for “Nipponbare” and “Koshihikari”, respectively. Dotted lines in red from top to bottom indicate molecular weight sizes of 1,000, 500, 400, 300, 200, and 100 bp. Bands for which the sequence has been determined in our previous study [20] are indicated with arrows and marker names in blue.

**Additional file 4 (XLSX) Supplementary Table S3** Restriction fragments with EcoRI and MseI ends from the tomato genome.

**Additional file 5 (XLSX) Supplementary Table S4** Restriction fragments with PstI and MseI ends from the tomato genome.

**Additional file 6 (PDF) Supplementary Figure S2** Distributions of the numbers of restriction fragments digested with EcoRI and MseI over the tomato genome.

The non-overlapping window size is 1 Mb. Black and white bars indicate hetero- and euchromatic regions.

**Additional file 7 (PDF) Supplementary Figure S3** Distributions of the numbers of restriction fragments digested with PstI and MseI over the tomato genome. The non-overlapping window size is 1 Mb. Black and white bars indicate hetero- and euchromatic regions.

**Additional file 8 (XLSX) Supplementary Table S5** Restriction fragments with PstI and MspI ends from the tomato genome.

**Additional file 9 (PDF) Supplementary Figure S4** Distributions of the numbers of restriction fragments digested with PstI and MspI over the tomato genome. The non-overlapping window size is 1 Mb. Black and white bars indicate hetero- and euchromatic regions.

**Additional file 10 (PDF) Supplementary Figure S5** Distributions of the numbers of genes predicted from the tomato genome.

The non-overlapping window size is 1 Mb. Black and white bars indicate hetero- and euchromatic regions.

**Additional file 11 (PDF) Supplementary Figure S6** Distributions of the numbers of restriction fragments digested with EcoRI and MseI over the rice genome. The non-overlapping window size is 1 Mb.

**Additional file 12 (XLSX) Supplementary Table S6** Restriction fragments with EcoRI and MseI ends from the rice genome.

**Additional file 13 (XLSX) Supplementary Table S7** Restriction fragments with PstI and MseI ends from the rice genome.

**Additional file 14 (XLSX) Supplementary Table S8** Restriction fragments with PstI and MspI ends from the rice genome.

**Additional file 15 (PDF) Supplementary Figure S7** Distributions of the numbers of restriction fragments digested with PstI and MseI over the rice genome.

The non-overlapping window size is 1 Mb.

**Additional file 16 (PDF) Supplementary Figure S8** Distributions of the numbers of restriction fragments digested with PstI and MspI over the rice genome.

The non-overlapping window size is 1 Mb.

**Additional file 17 (PDF) Supplementary Figure S9** Distributions of the numbers of genes predicted from the rice genome.

The non-overlapping window size is 1 Mb.

**Additional file 18 (XLSX) Supplementary Table S9** Restriction fragments with EcoRI and MseI ends from the *Arabidopsis* genome.

**Additional file 19 (XLSX) Supplementary Table S10** Restriction fragments with PstI and MseI ends from the *Arabidopsis* genome.

**Additional file 20 (XLSX) Supplementary Table S11** Restriction fragments with PstI and MspI ends from the *Arabidopsis* genome.

**Additional file 21 (PDF) Supplementary Figure S10** Distributions of the numbers of restriction fragments digested with EcoRI and MseI over the *Arabidopsis* genome.

The non-overlapping window size is 1 Mb.

**Additional file 22 (PDF) Supplementary Figure S11** Distributions of the numbers of restriction fragments digested with PstI and MseI over the *Arabidopsis* genome.

The non-overlapping window size is 1 Mb.

**Additional file 23 (PDF) Supplementary Figure S12** Distributions of the numbers of restriction fragments digested with PstI and MspI over the *Arabidopsis* genome.

The non-overlapping window size is 1 Mb.

**Additional file 24 (PDF) Supplementary Figure S13** Distributions of the numbers of genes predicted from the *Arabidopsis* genome.

The non-overlapping window size is 1 Mb.

**Additional file 25 (XLSX) Supplementary Table S12** Restriction fragments with EcoRI and MseI ends from the *Lotus* genome.

**Additional file 26 (XLSX) Supplementary Table S13** Restriction fragments with PstI and MseI ends from the *Lotus* genome.

**Additional file 27 (XLSX) Supplementary Table S14** Restriction fragments with PstI and MspI ends from the *Lotus* genome.

**Additional file 28 (PDF) Supplementary Figure S14** Distributions of the numbers of restriction fragments digested with EcoRI and MseI over the *Lotus* genome.

The non-overlapping window size is 1 Mb.

**Additional file 29 (PDF) Supplementary Figure S15** Distributions of the numbers of restriction fragments digested with PstI and MseI over the *Lotus* genome.

The non-overlapping window size is 1 Mb.

**Additional file 30 (PDF) Supplementary Figure S16** Distributions of the numbers of restriction fragments digested with PstI and MspI over the *Lotus* genome.

The non-overlapping window size is 1 Mb.

**Additional file 31 (PDF) Supplementary Figure S17** Distributions of the numbers of genes predicted from the *Lotus* genome.

The non-overlapping window size is 1 Mb.

## Notes

### Competing Interest Statement

The authors have declared no competing interest.

