## Supplementary Figure S1 for "Optimizing selection of restriction enzymes for complexity-reduced genome sequencing in plants"

### Contents of additional file 3

**Table.** Page numbers of AFLP images for each selective primer combination

| MseI | EcoRI | | | | | | | | | | | |
|---|---|---|---|---|---|---|---|---|---|---|---|---|
| | ACG | AGC | ATG | CAA | CAC | CAG | CAT | CGT | GCA | GTA | TAC | TGC |
| ACG | 1 | 1 | 1 | 2 | 2 | 2 | 3 | 3 | 3 | 4 | 4 | 4 |
| AGC | 5 | 5 | 5 | 6 | 6 | 6 | 7 | 7 | 7 | 8 | 8 | 8 |
| ATG | 9 | 9 | 9 | 10 | 10 | 10 | 11 | 11 | 11 | 12 | 12 | 12 |
| CAA | 13 | 13 | 13 | 14 | 14 | 14 | 15 | 15 | 15 | 16 | 16 | 16 |
| CAC | 17 | 17 | 17 | 18 | 18 | 18 | 19 | 19 | 19 | 20 | 20 | 20 |
| CAG | 21 | 21 | 21 | 22 | 22 | 22 | 23 | 23 | 23 | 24 | 24 | 24 |
| CAT | 25 | 25 | 25 | 26 | 26 | 26 | 27 | 27 | 27 | 28 | 28 | 28 |
| CGT | 29 | 29 | 29 | 30 | 30 | 30 | 31 | 31 | 31 | 32 | 32 | 32 |
| GCA | 33 | 33 | 33 | 34 | 34 | 34 | 35 | 35 | 35 | 36 | 36 | 36 |
| GTA | 37 | 37 | 37 | 38 | 38 | 38 | 39 | 39 | 39 | 40 | 40 | 40 |
| TAC | 41 | 41 | 41 | 42 | 42 | 42 | 43 | 43 | 43 | 44 | 44 | 44 |
| TGC | 45 | 45 | 45 | 46 | 46 | 46 | 47 | 47 | 47 | 48 | 48 | 48 |

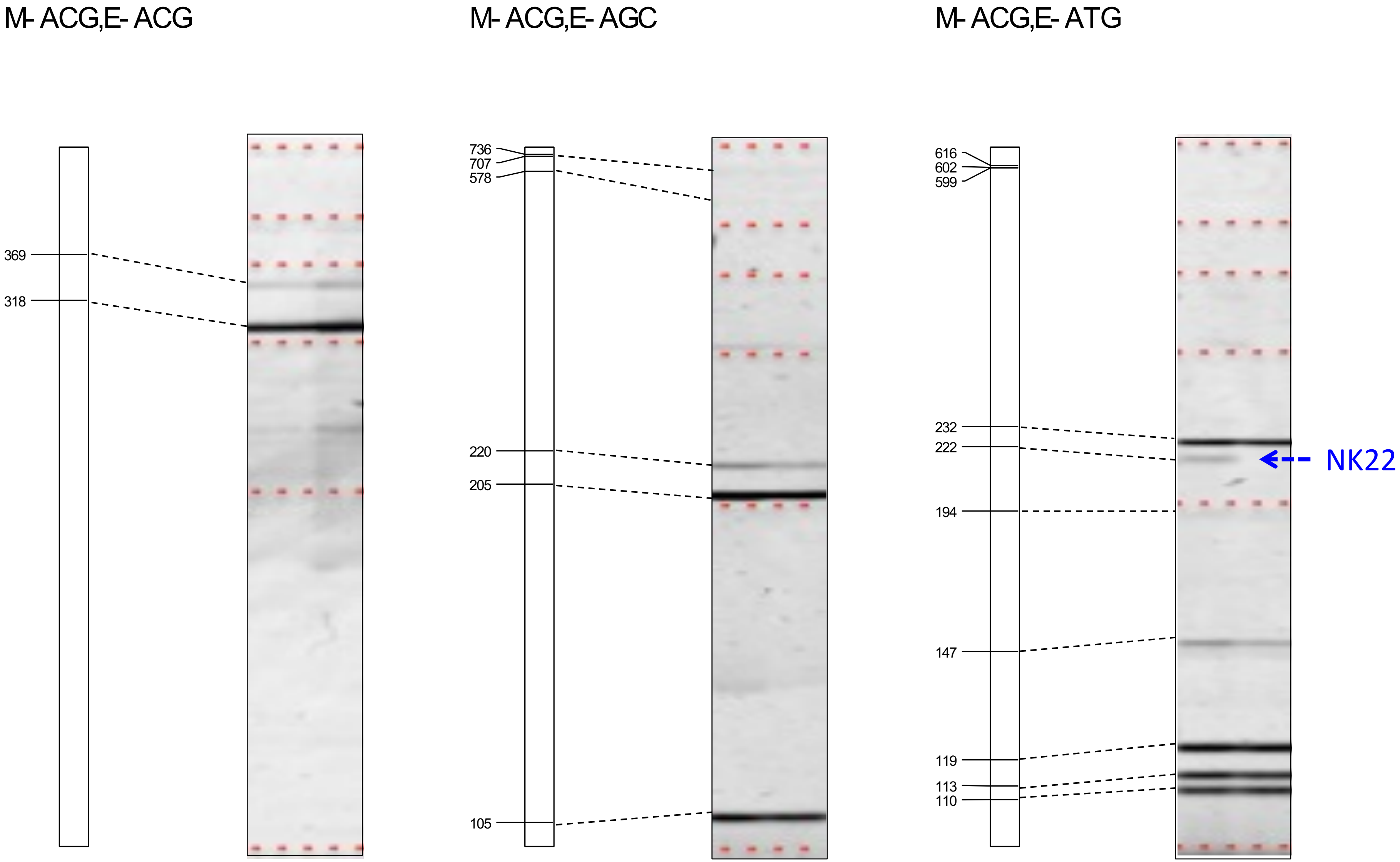


**Supplementary Figure S1** *In silico* AFLP and experimental data.

The left and right pictures are the *in silico* and actual experimental AFLP data, respectively. Primer combinations for MseI (M) and EcoRI (E) are shown on the tops of the pictures. Fragment sizes predicted in the *in silico* AFLP analysis are indicated by numbers, each of which corresponds to a single AFLP fragment. The left and right lanes in the actual images are for “Nipponbare” and “Koshihikari”, respectively. Dotted lines in red from top to bottom indicate molecular weight sizes of 1,000, 500, 400, 300, 200, and 100 bp. Bands for which the sequence has been determined in our previous study (Shirasawa *et al.*, 2004) are indicated with arrows and marker names in blue.

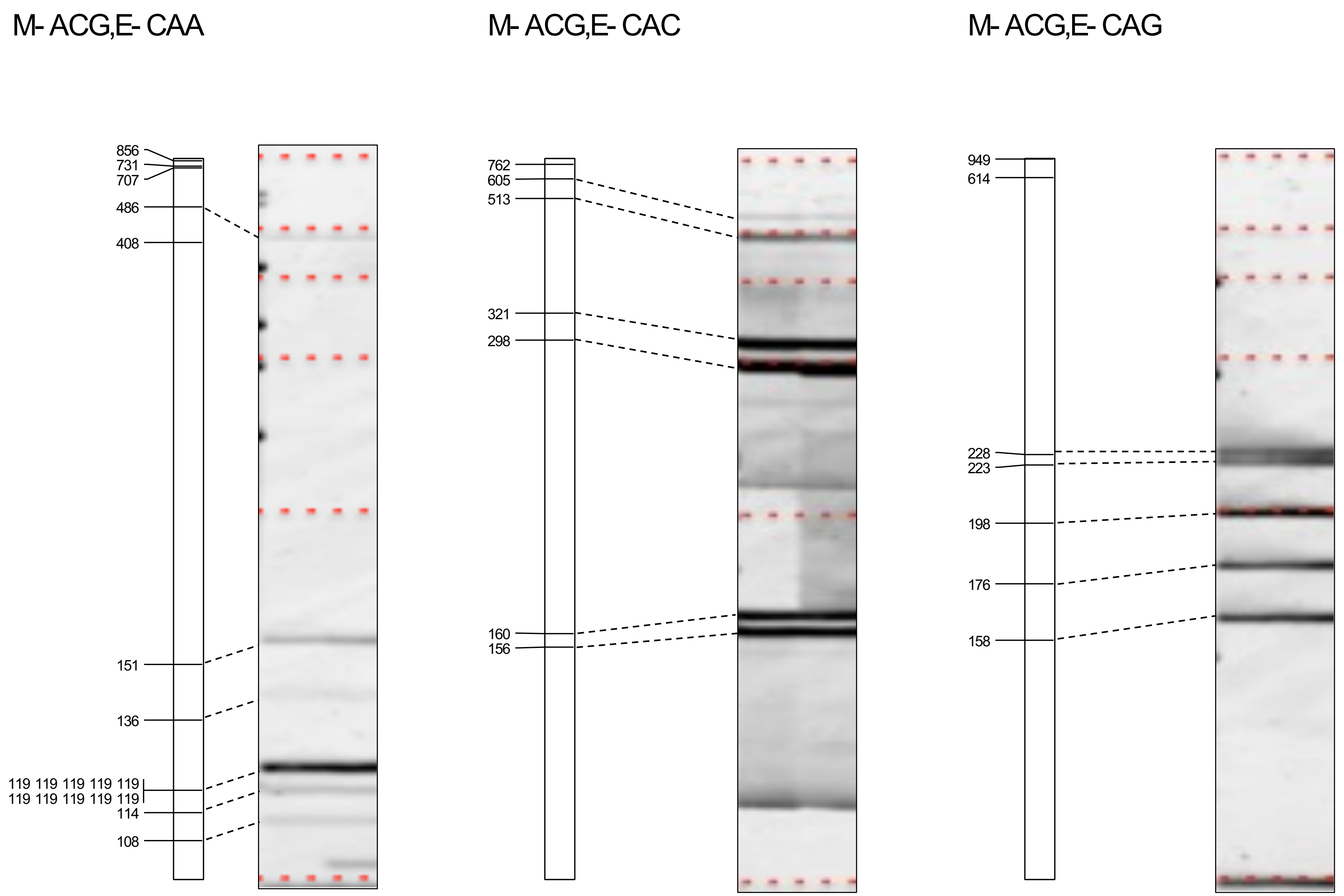


**Supplementary Figure S1 (continued)** *In silico* AFLP and experimental data.

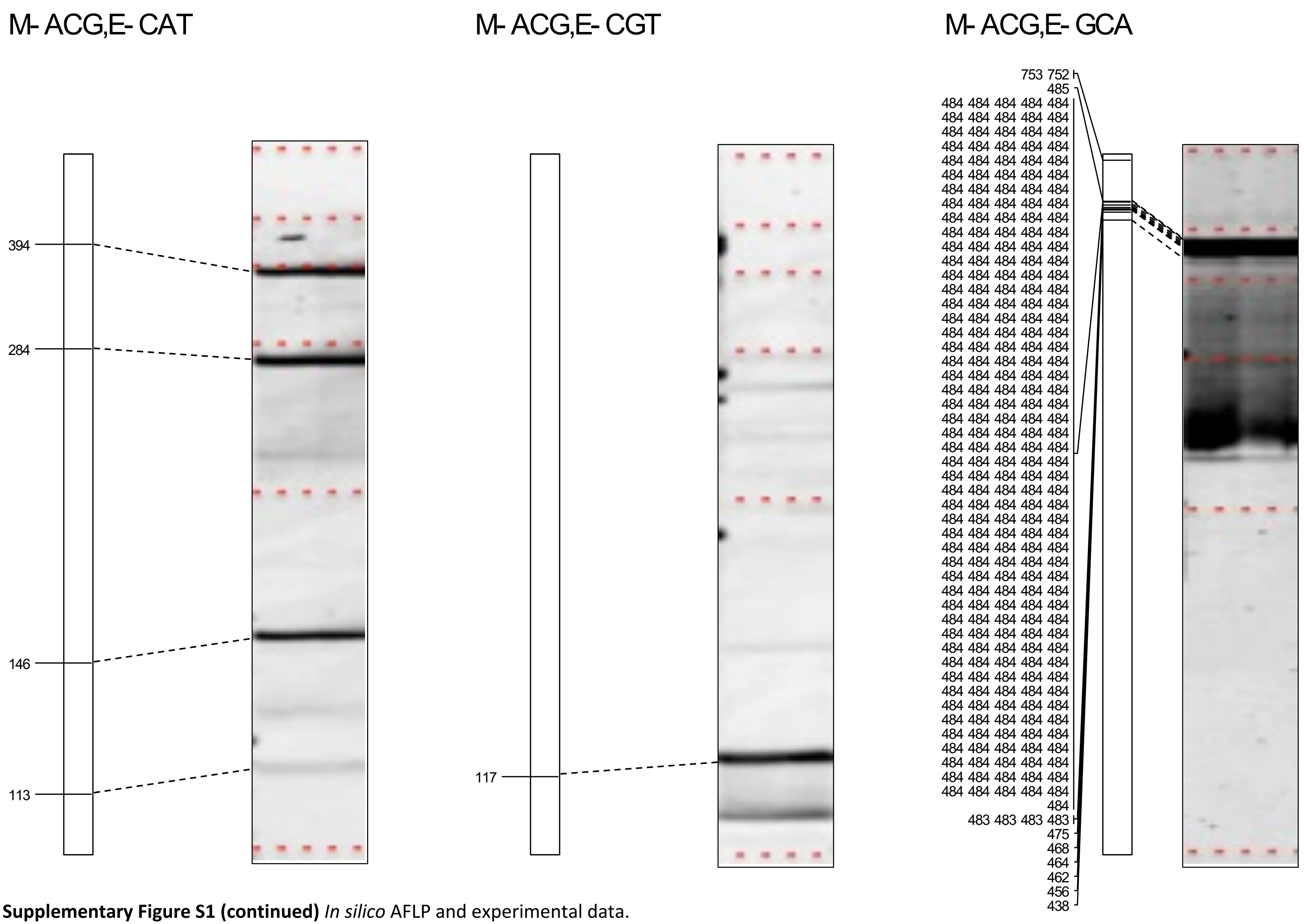


**Supplementary Figure S1 (continued)** *In silico* AFLP and experimental data.

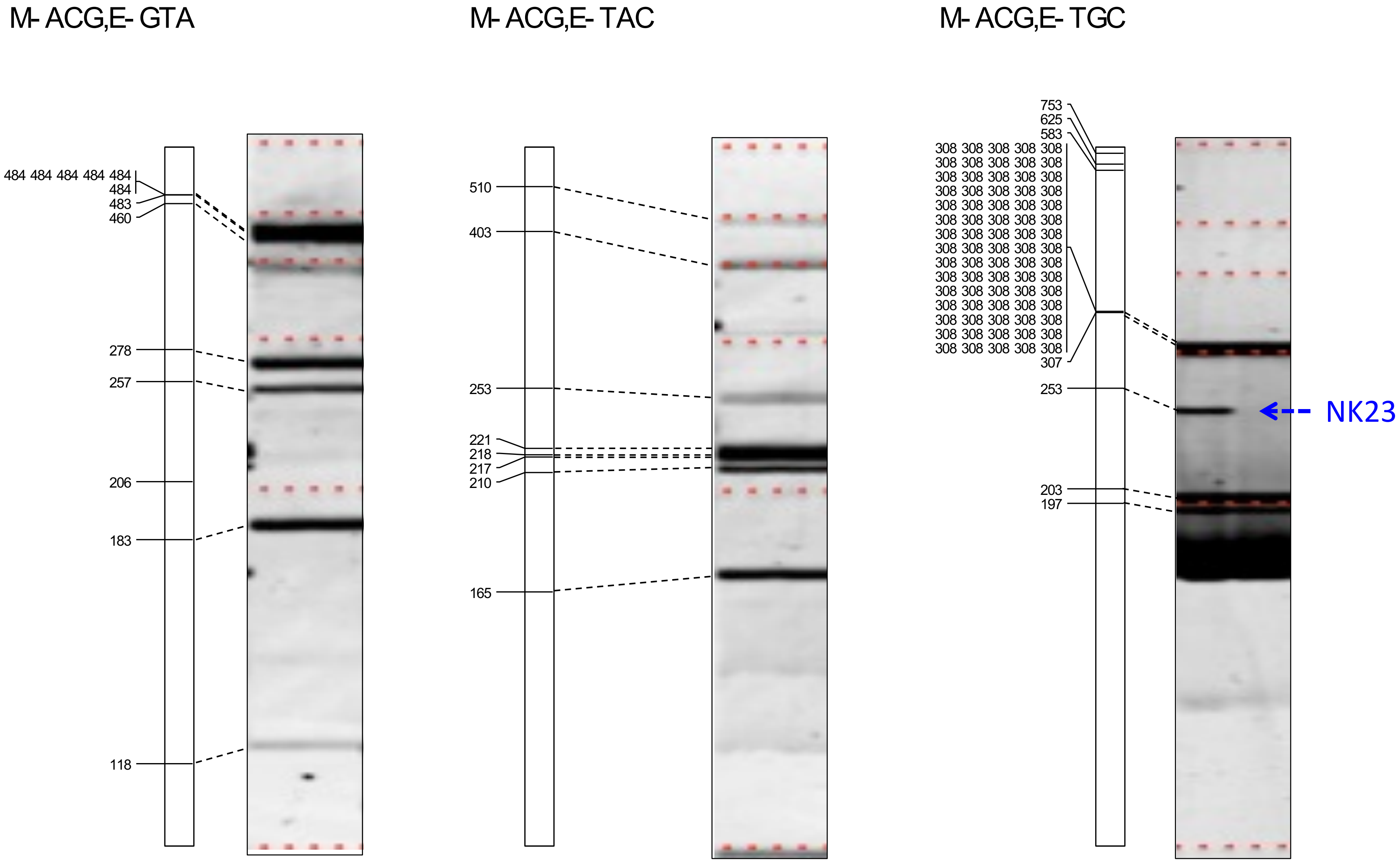


**Supplementary Figure S1 (continued)** *In silico* AFLP and experimental data.

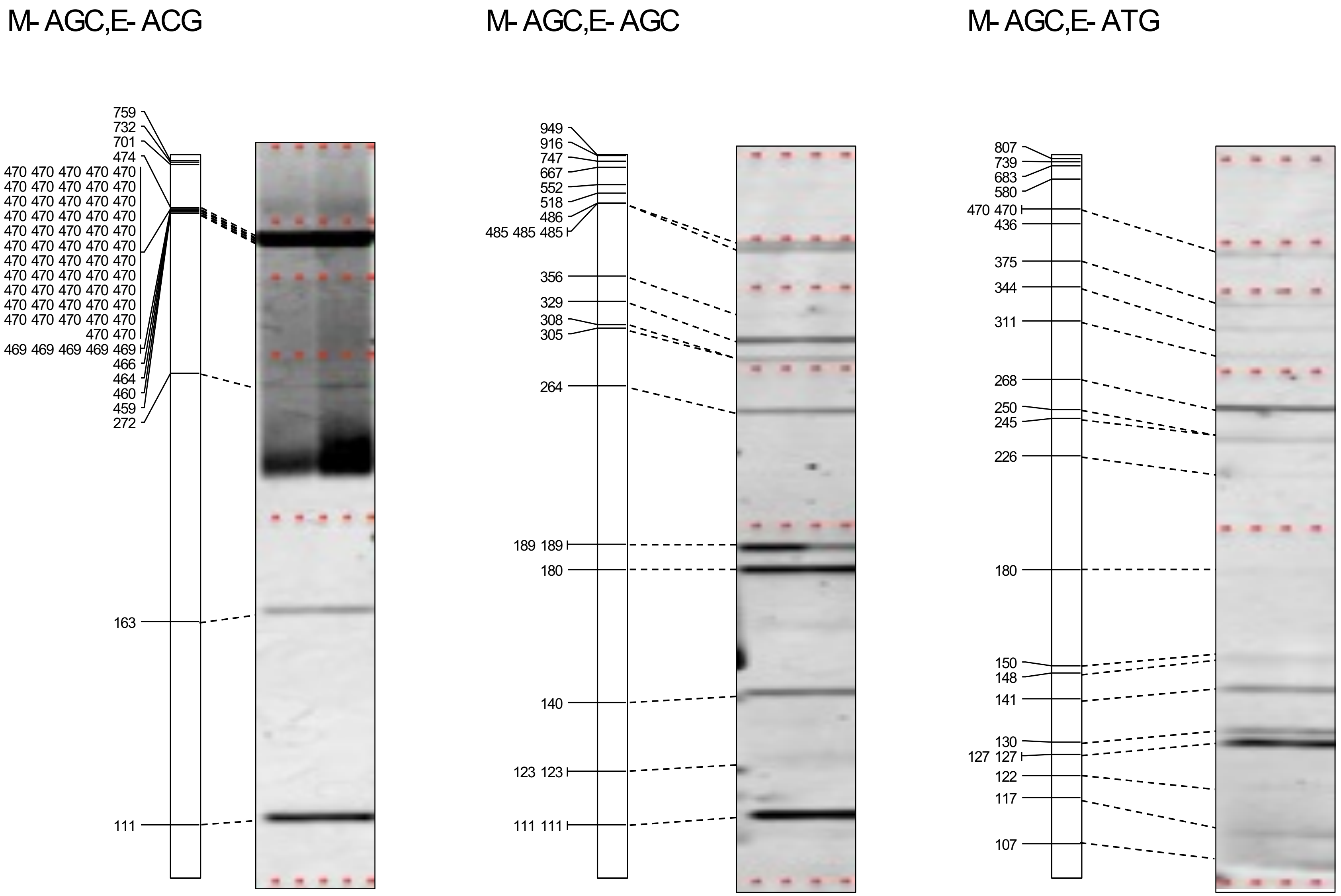


**Supplementary Figure S1 (continued)** *In silico* AFLP and experimental data.

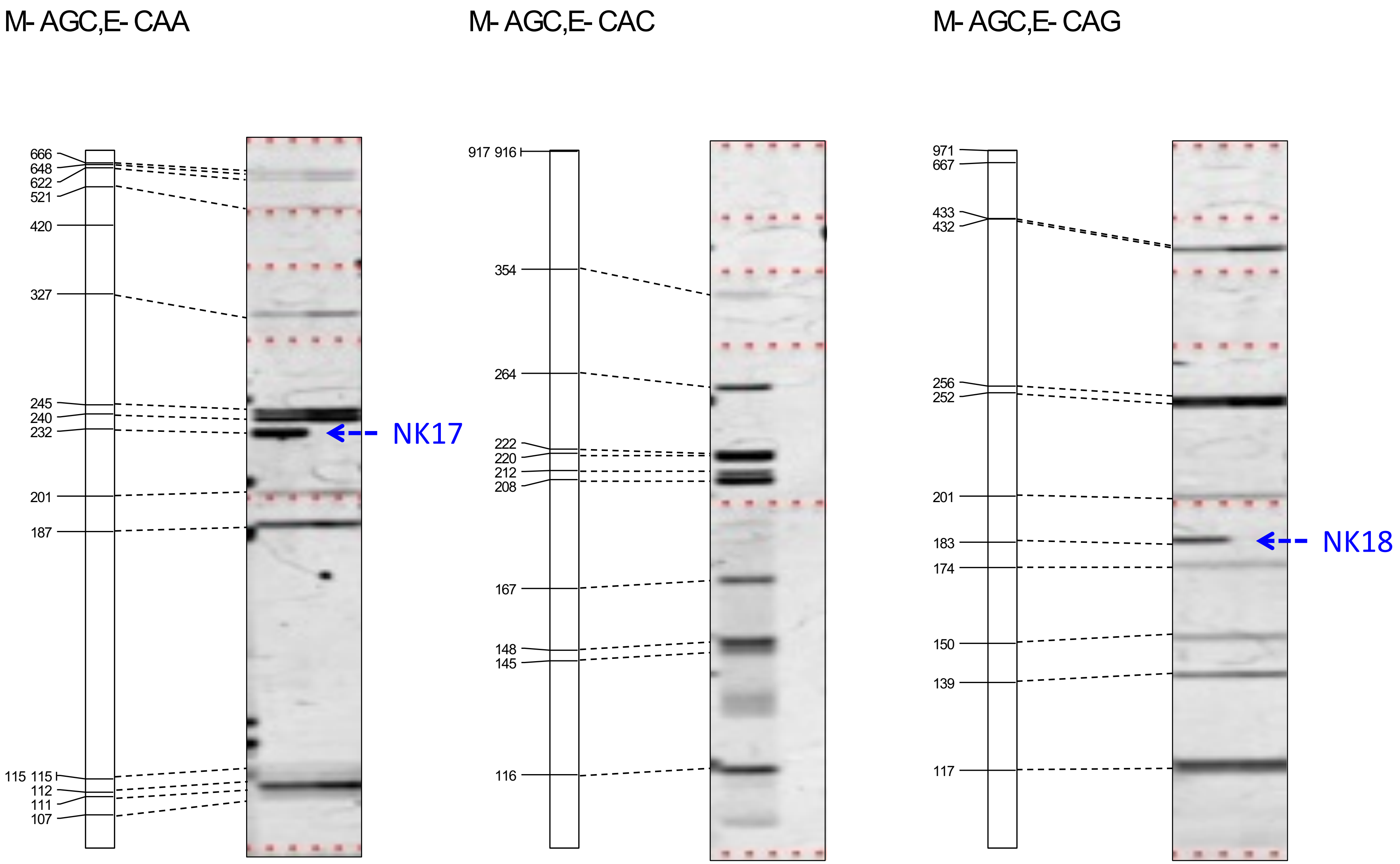


**Supplementary Figure S1 (continued)** *In silico* AFLP and experimental data.

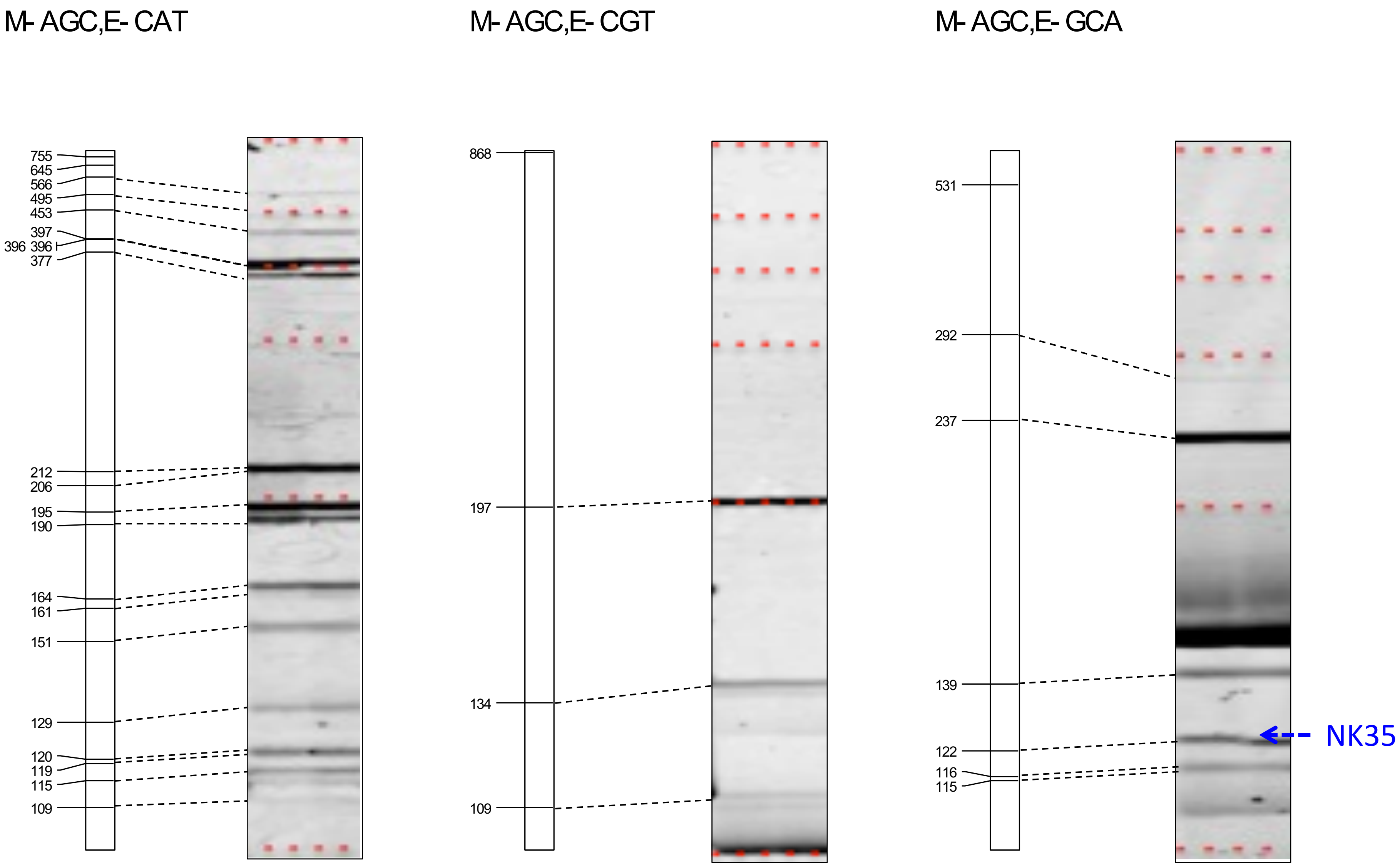


**Supplementary Figure S1 (continued)** *In silico* AFLP and experimental data.

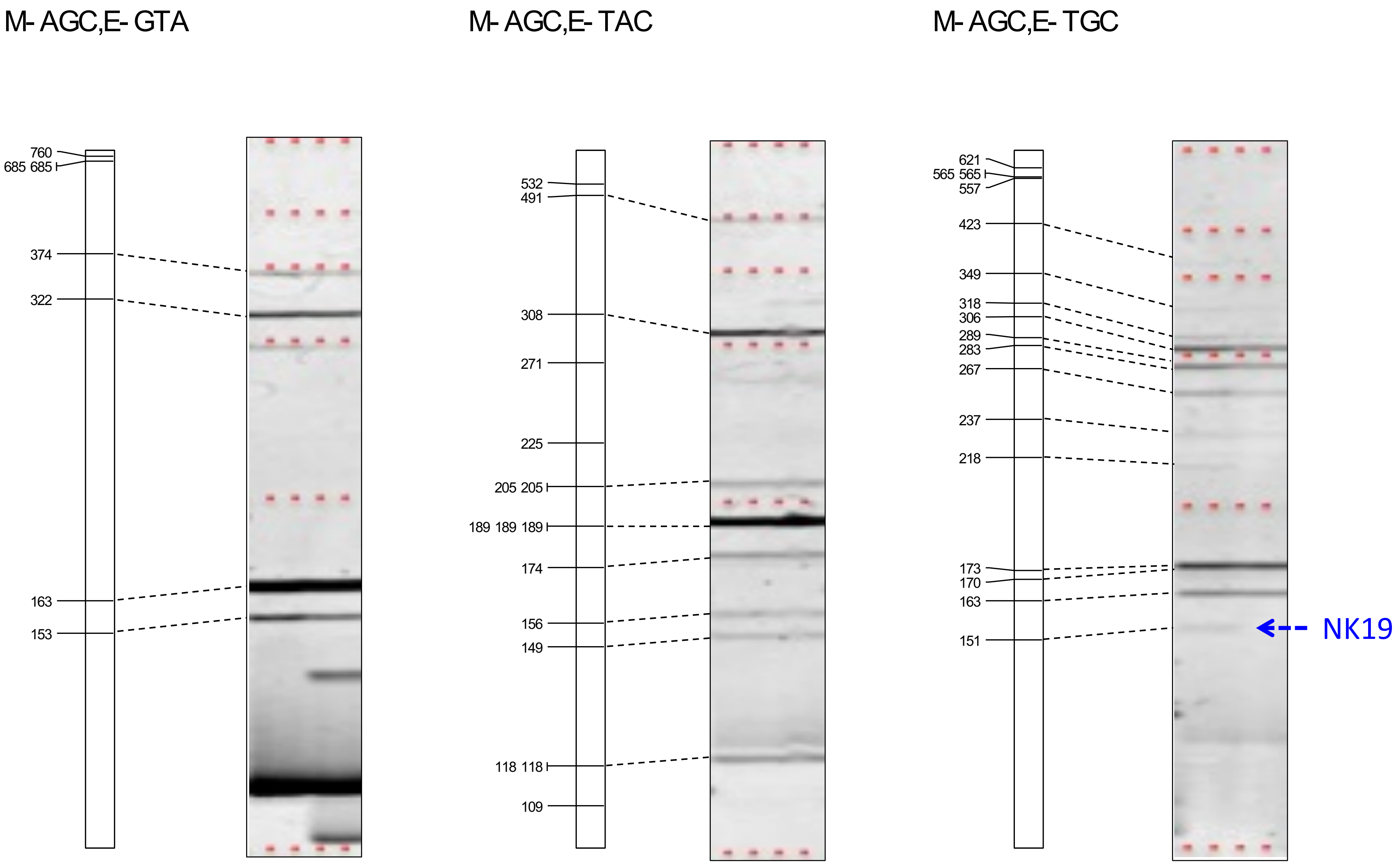


**Supplementary Figure S1 (continued)** *In silico* AFLP and experimental data.

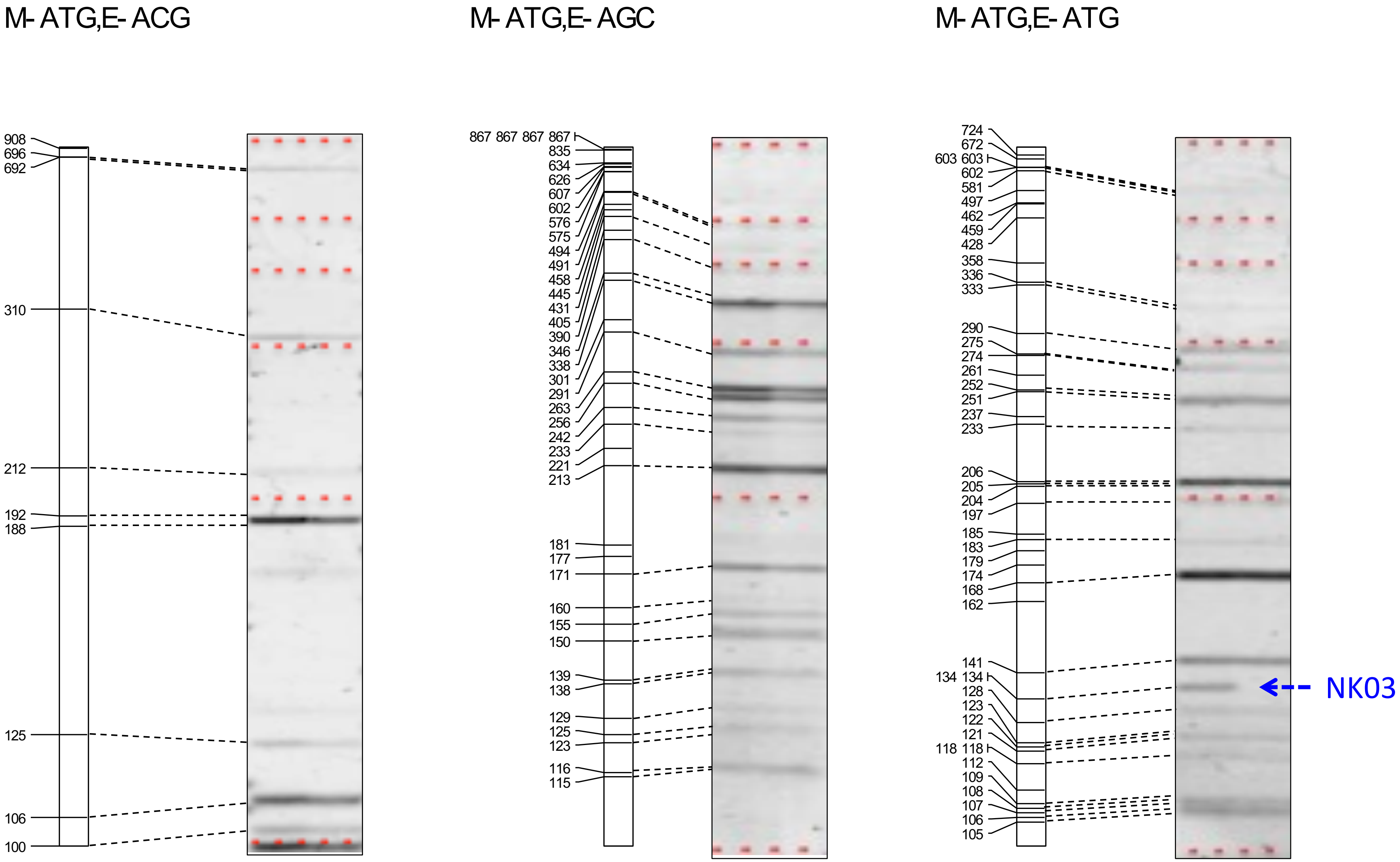


**Supplementary Figure S1 (continued)** *In silico* AFLP and experimental data.

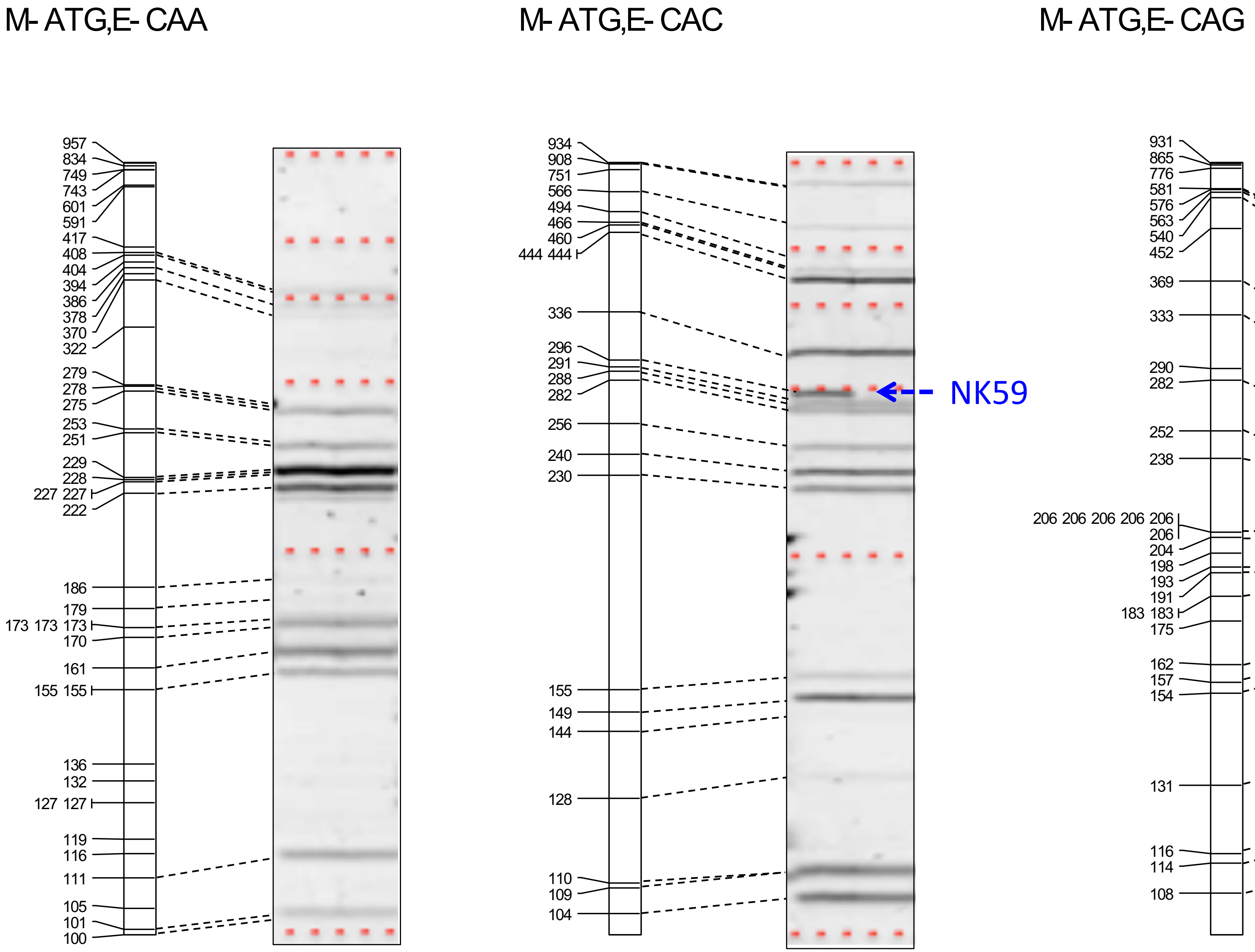


**Supplementary Figure S1 (continued)** *In silico* AFLP and experimental data.

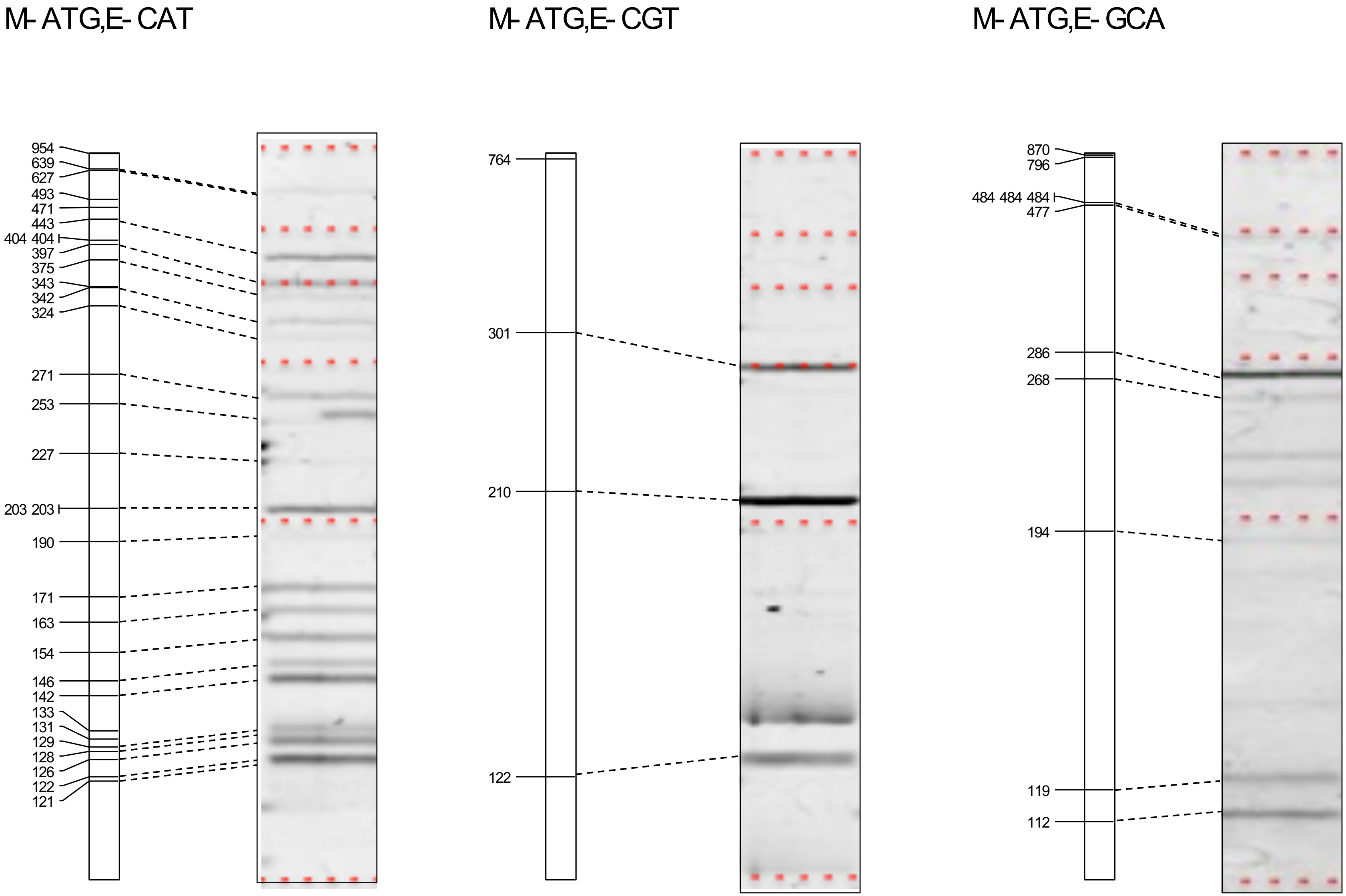


**Supplementary Figure S1 (continued)** *In silico* AFLP and experimental data.

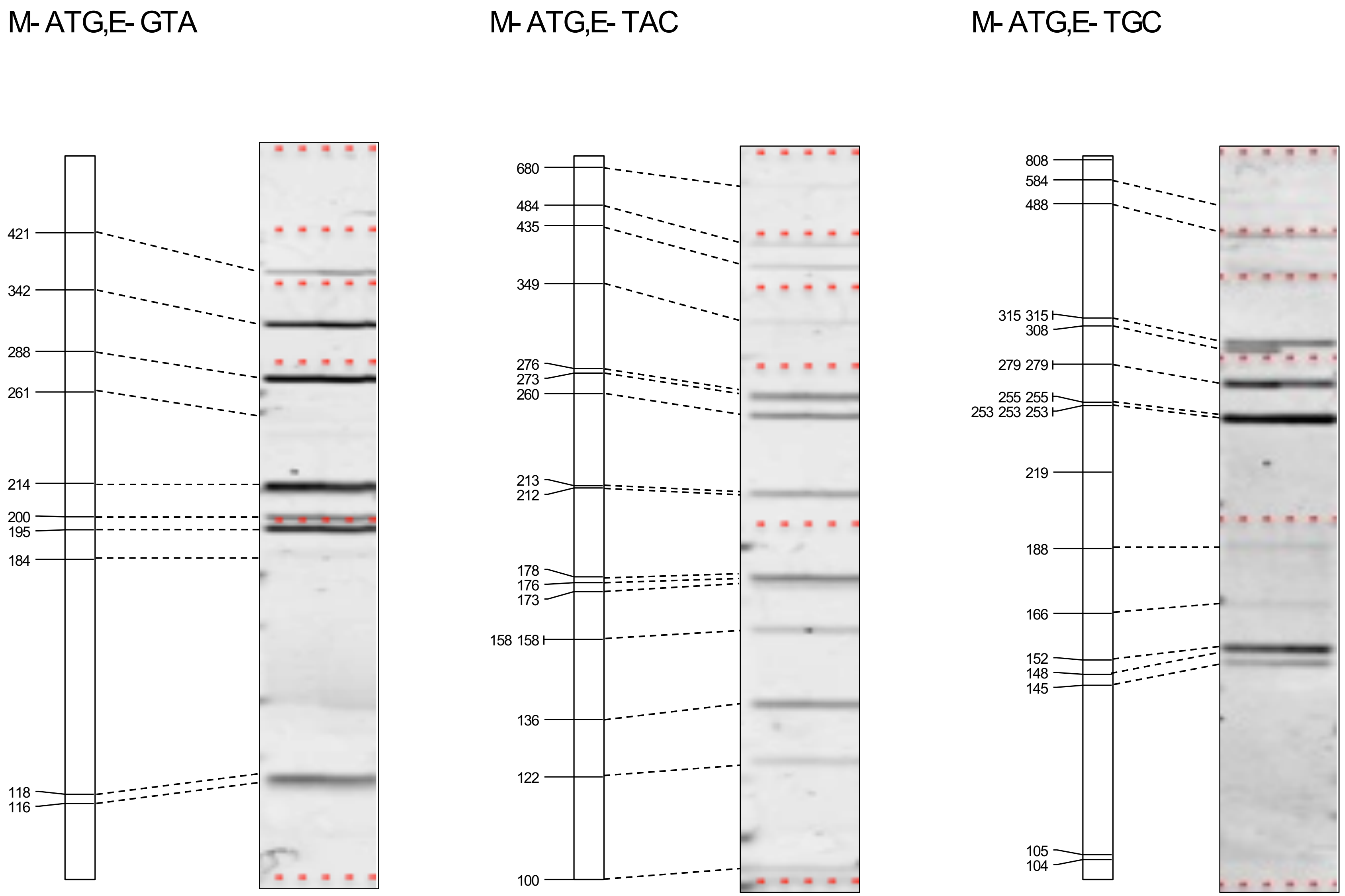


**Supplementary Figure S1 (continued)** *In silico* AFLP and experimental data.

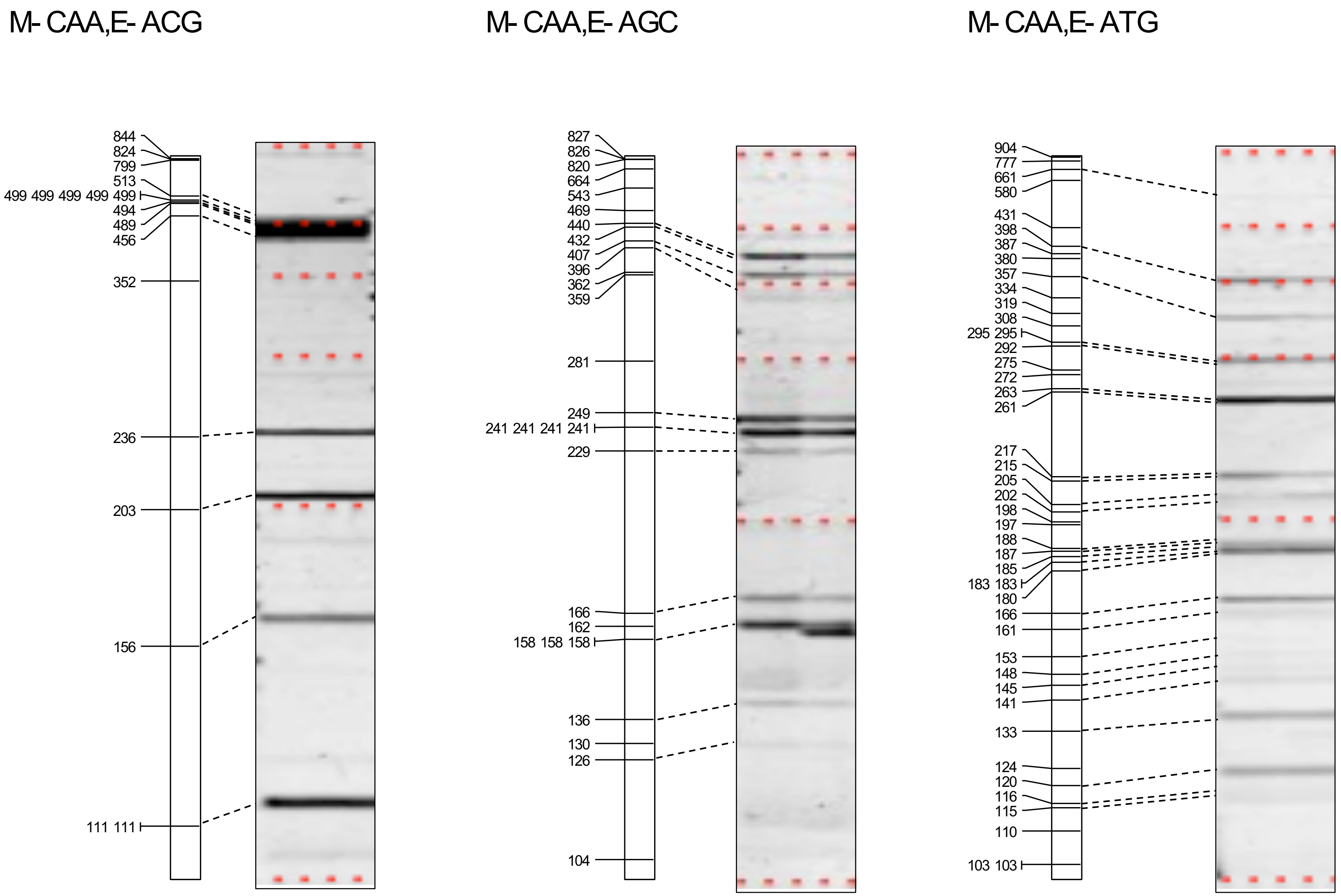


**Supplementary Figure S1 (continued)** *In silico* AFLP and experimental data.

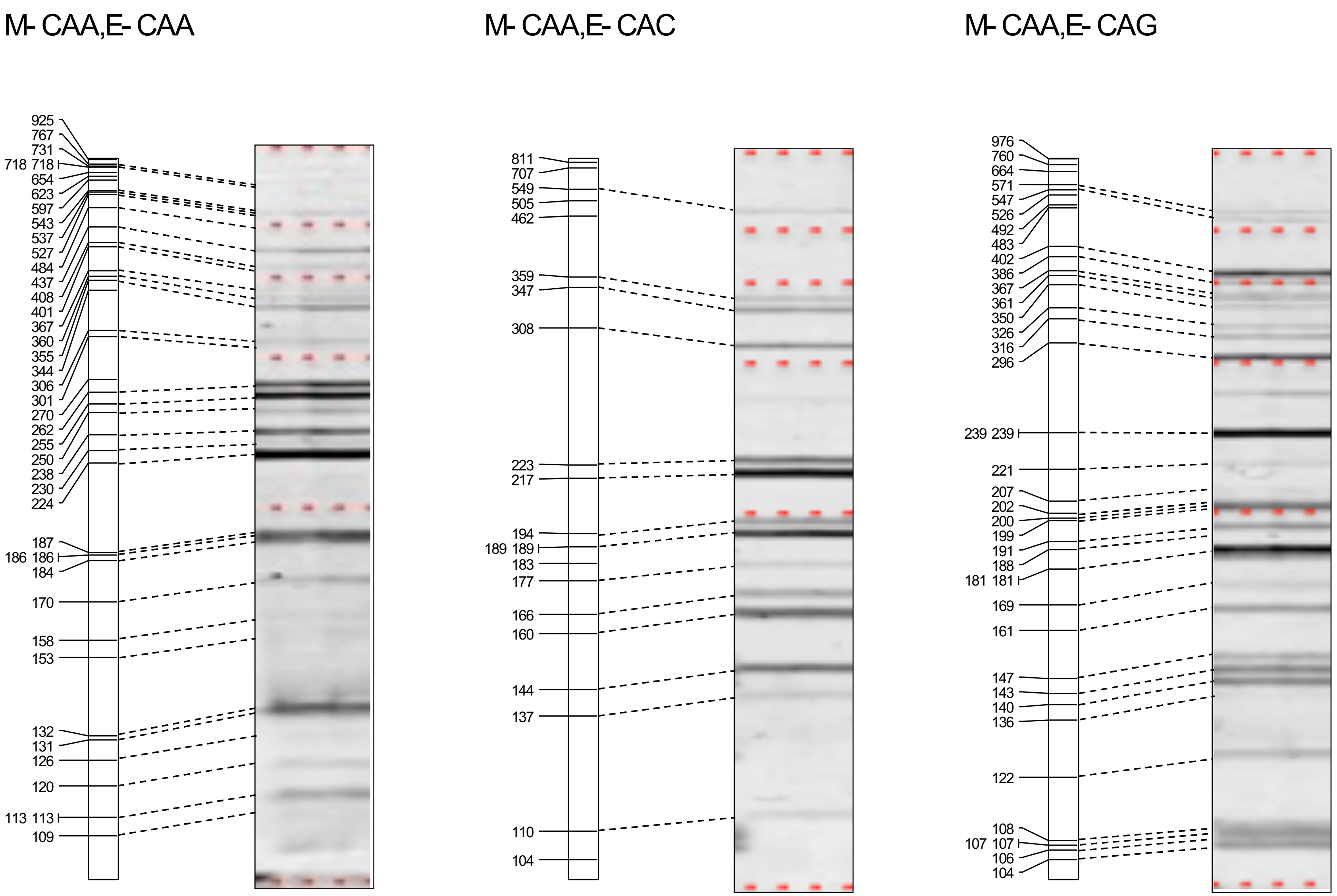


**Supplementary Figure S1 (continued)** *In silico* AFLP and experimental data.

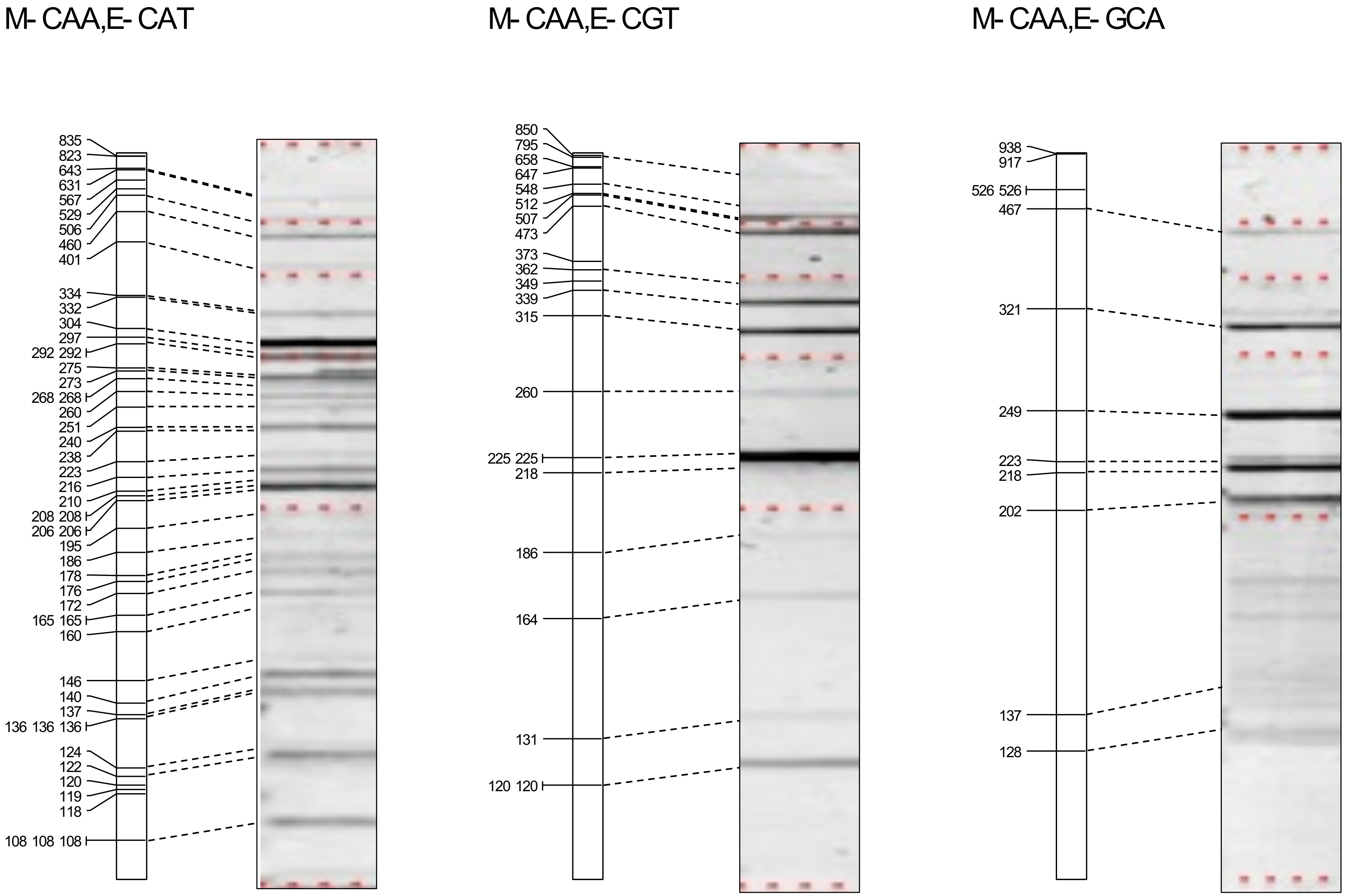


**Supplementary Figure S1 (continued)** *In silico* AFLP and experimental data.

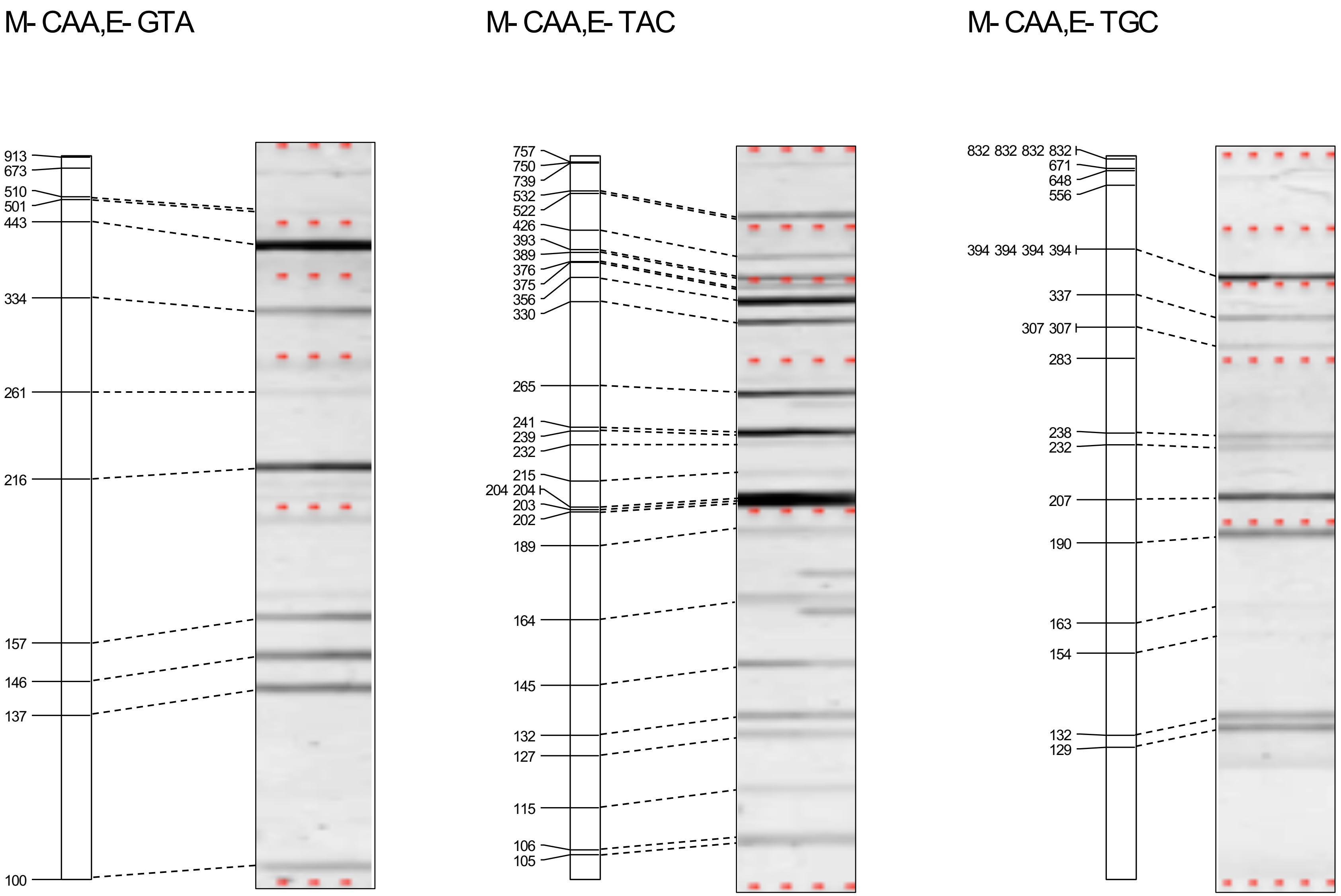


**Supplementary Figure S1 (continued)** *In silico* AFLP and experimental data.

M- CAC,E- ACG

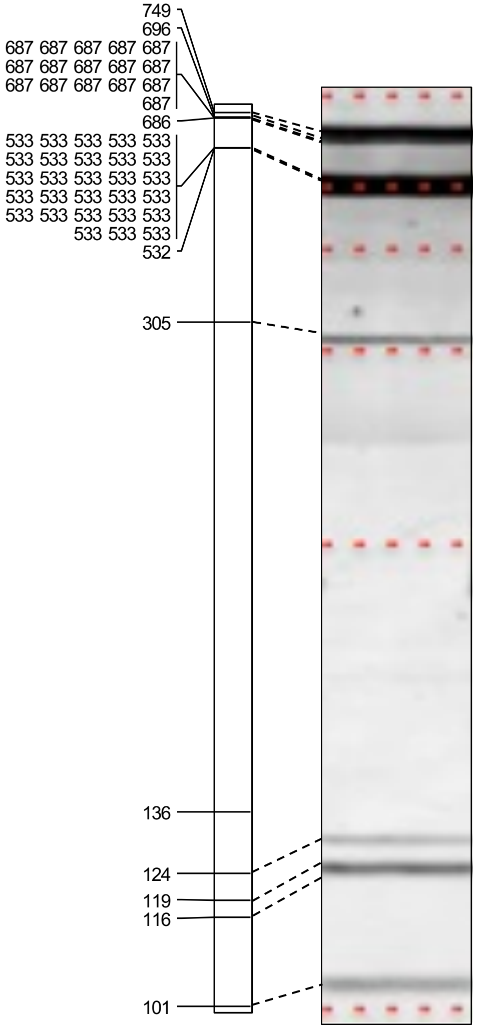


M- CAC,E- AGC

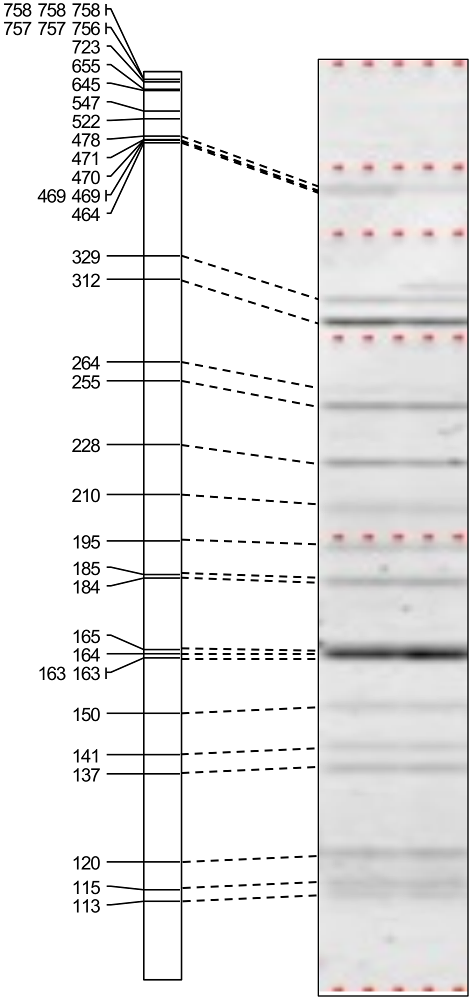


M- CAC,E- ATG

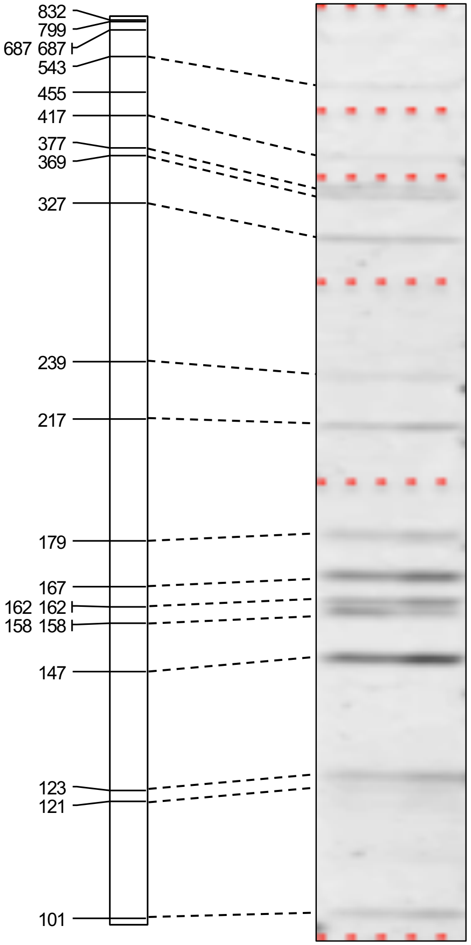


**Supplementary Figure S1 (continued)** *In silico* AFLP and experimental data.

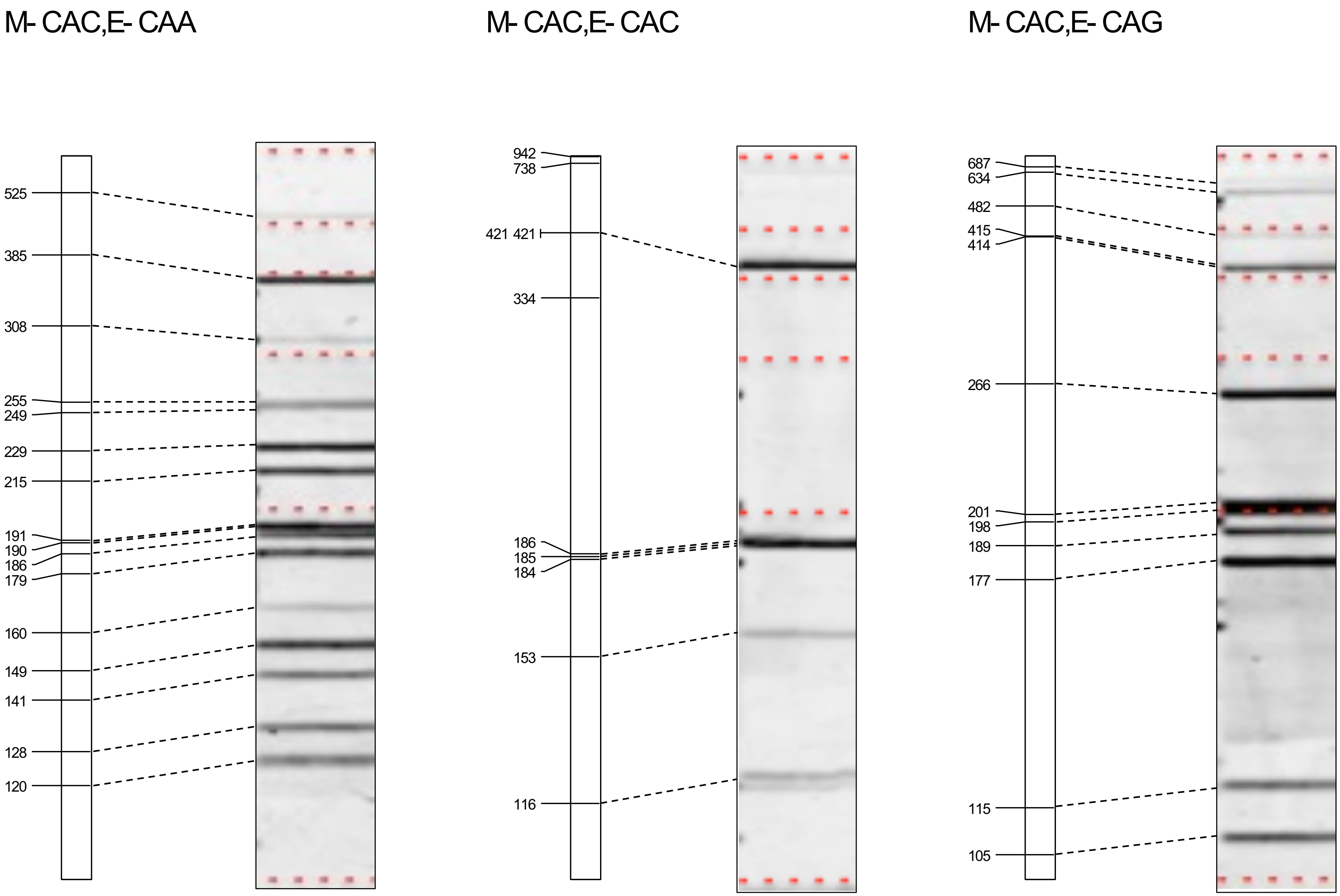


**Supplementary Figure S1 (continued)** *In silico* AFLP and experimental data.

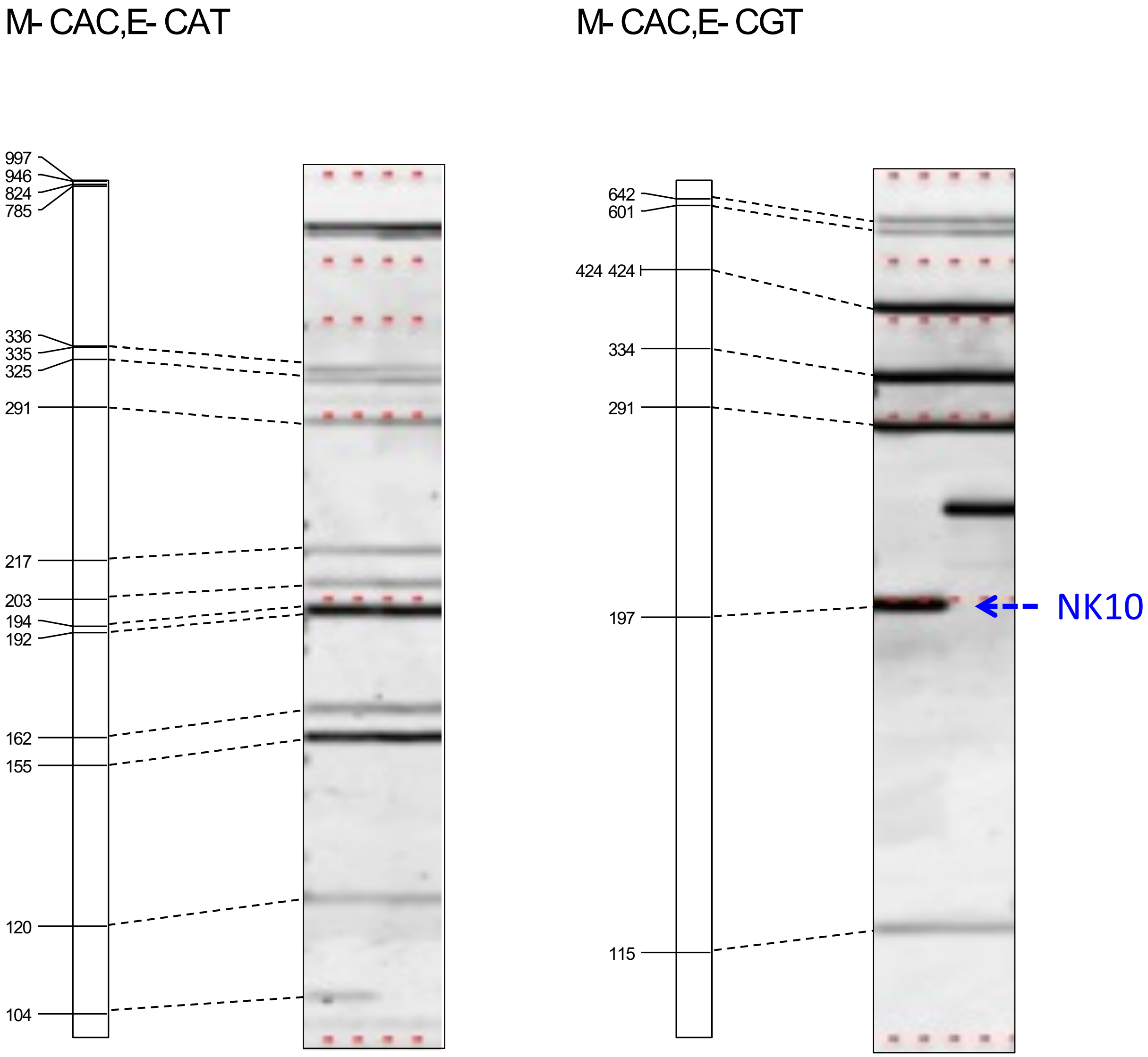


**Supplementary Figure S1 (continued)** *In silico* AFLP and experimental data.

M- CAC,E- GTA

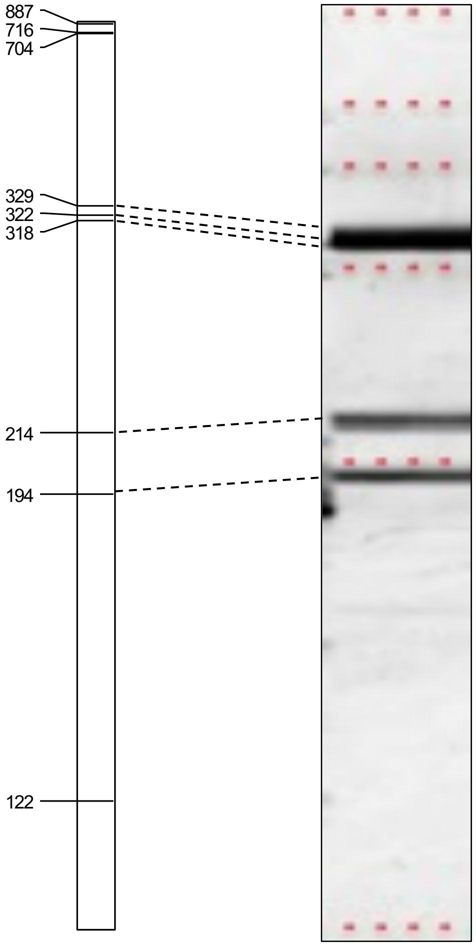


M- CAC,E- TAC

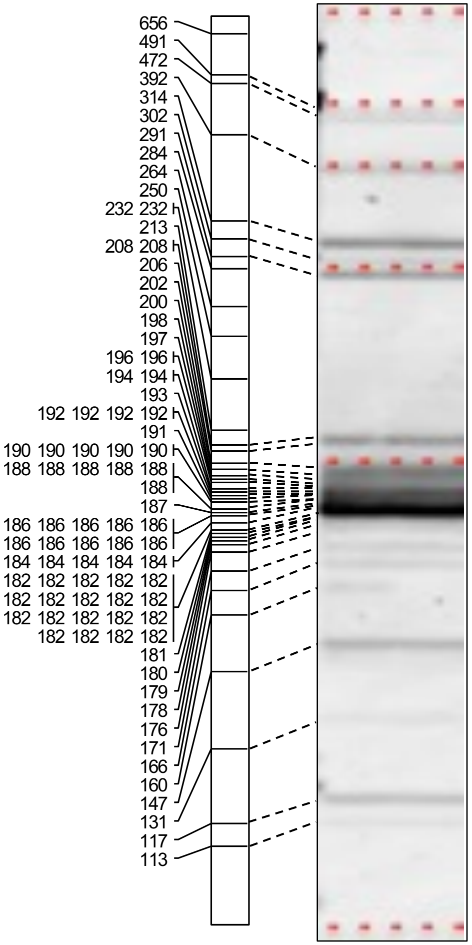


M- CAC,E- TGC

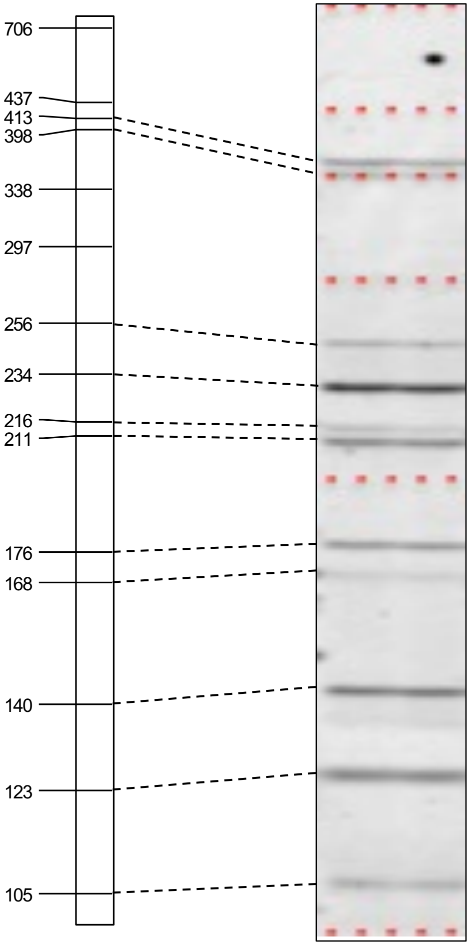


**Supplementary Figure S1 (continued)** *In silico* AFLP and experimental data.

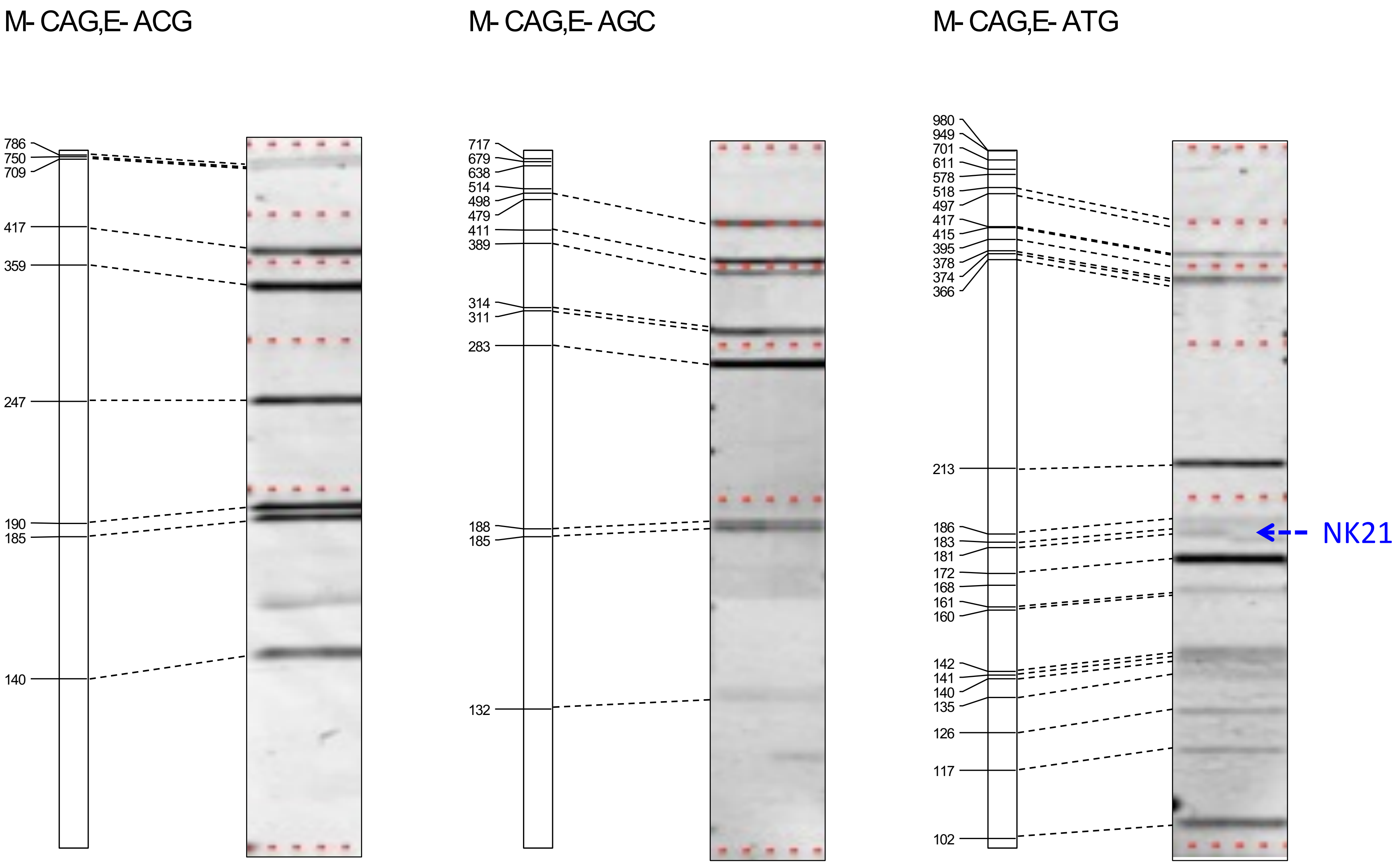


**Supplementary Figure S1 (continued)** *In silico* AFLP and experimental data.

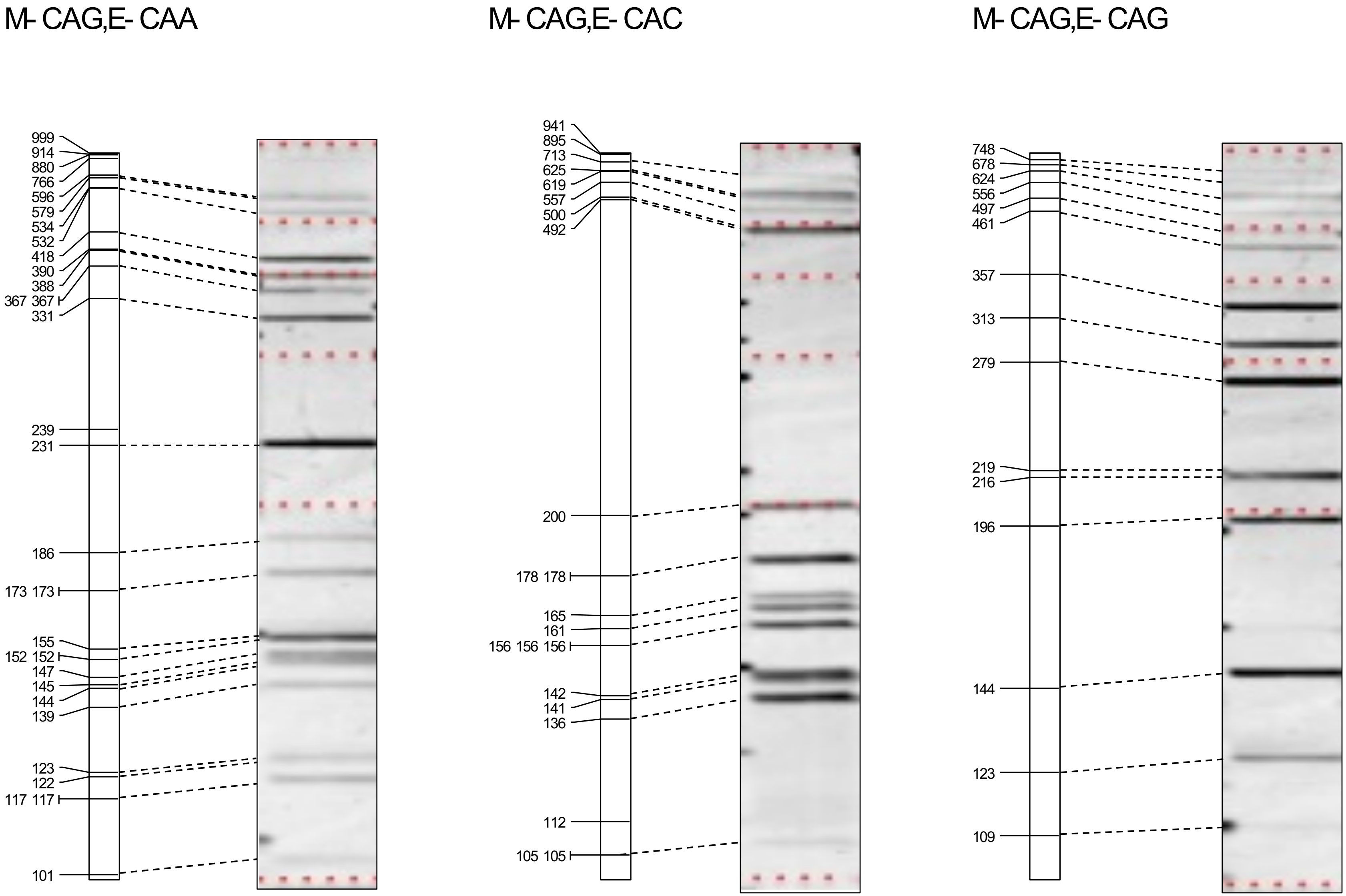


**Supplementary Figure S1 (continued)** *In silico* AFLP and experimental data.

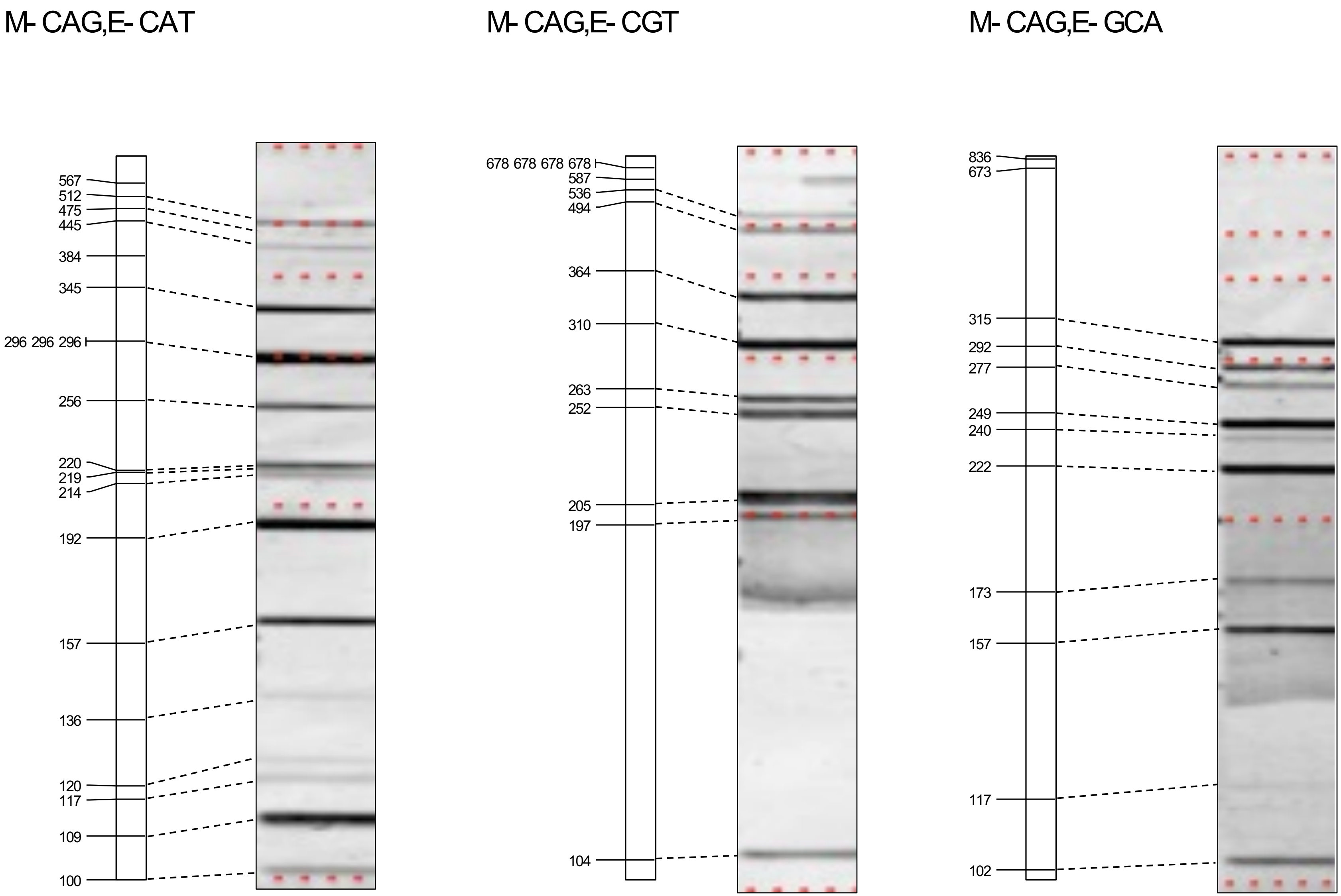


**Supplementary Figure S1 (continued)** *In silico* AFLP and experimental data.

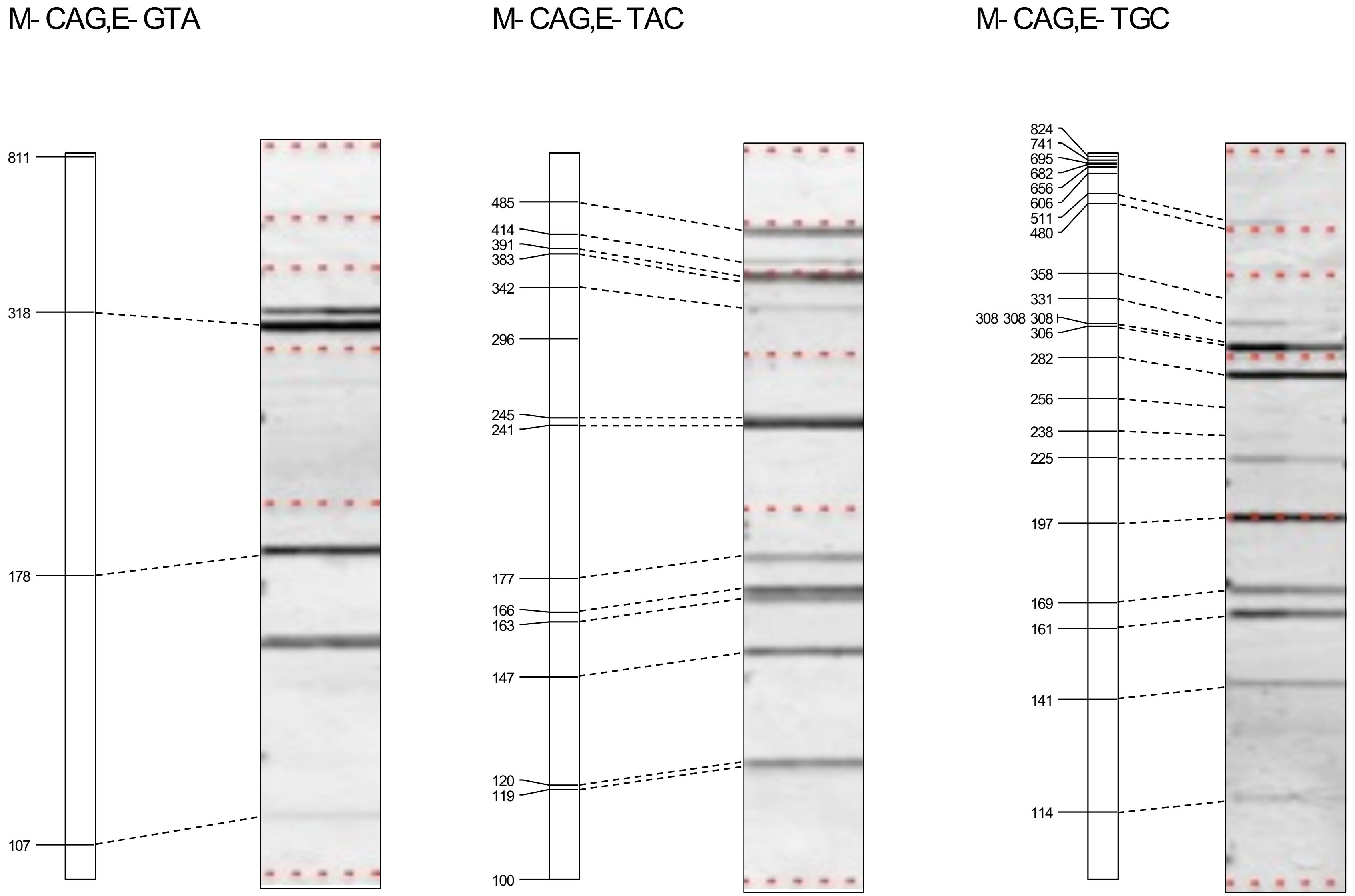


**Supplementary Figure S1 (continued)** *In silico* AFLP and experimental data.

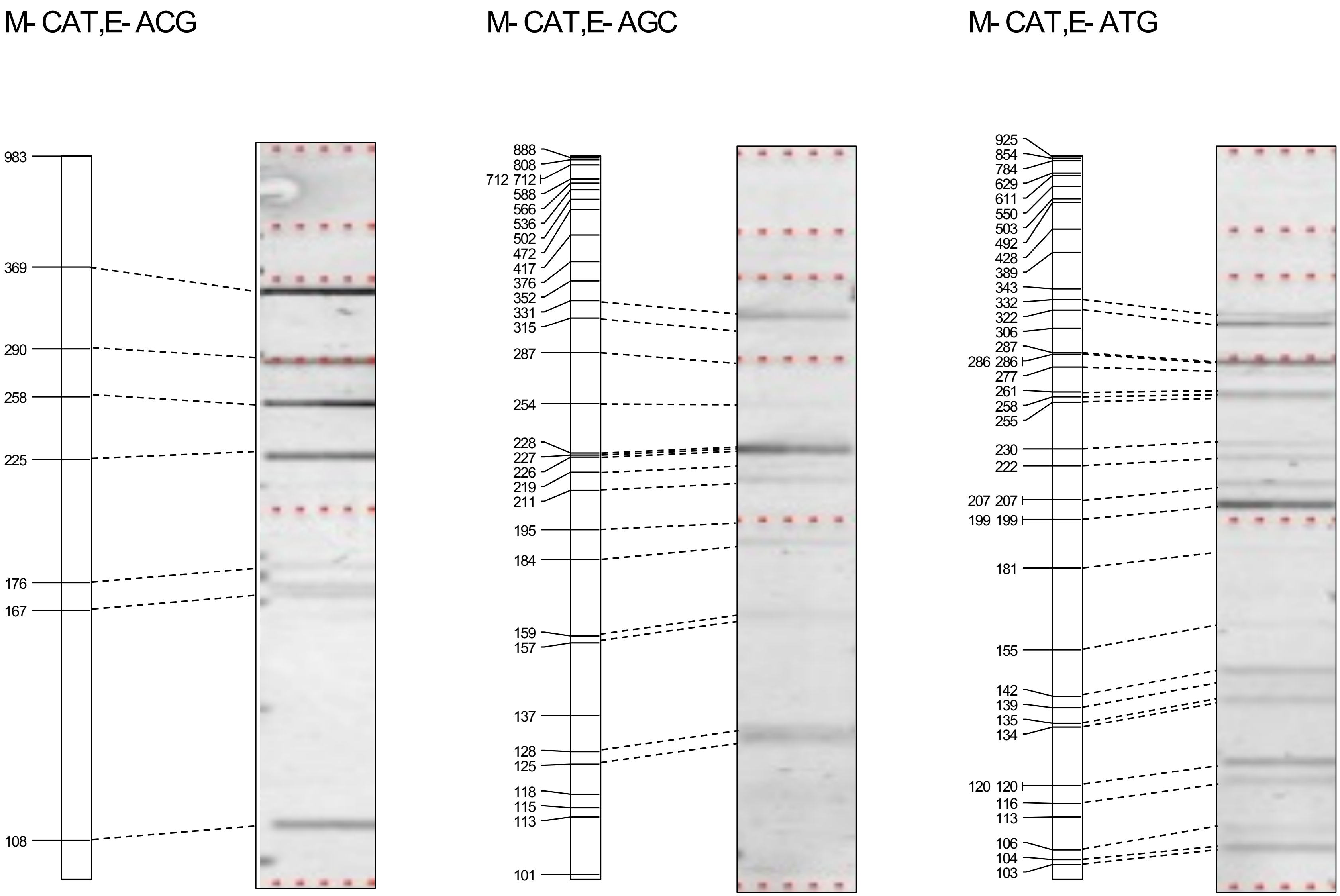


**Supplementary Figure S1 (continued)** *In silico* AFLP and experimental data.

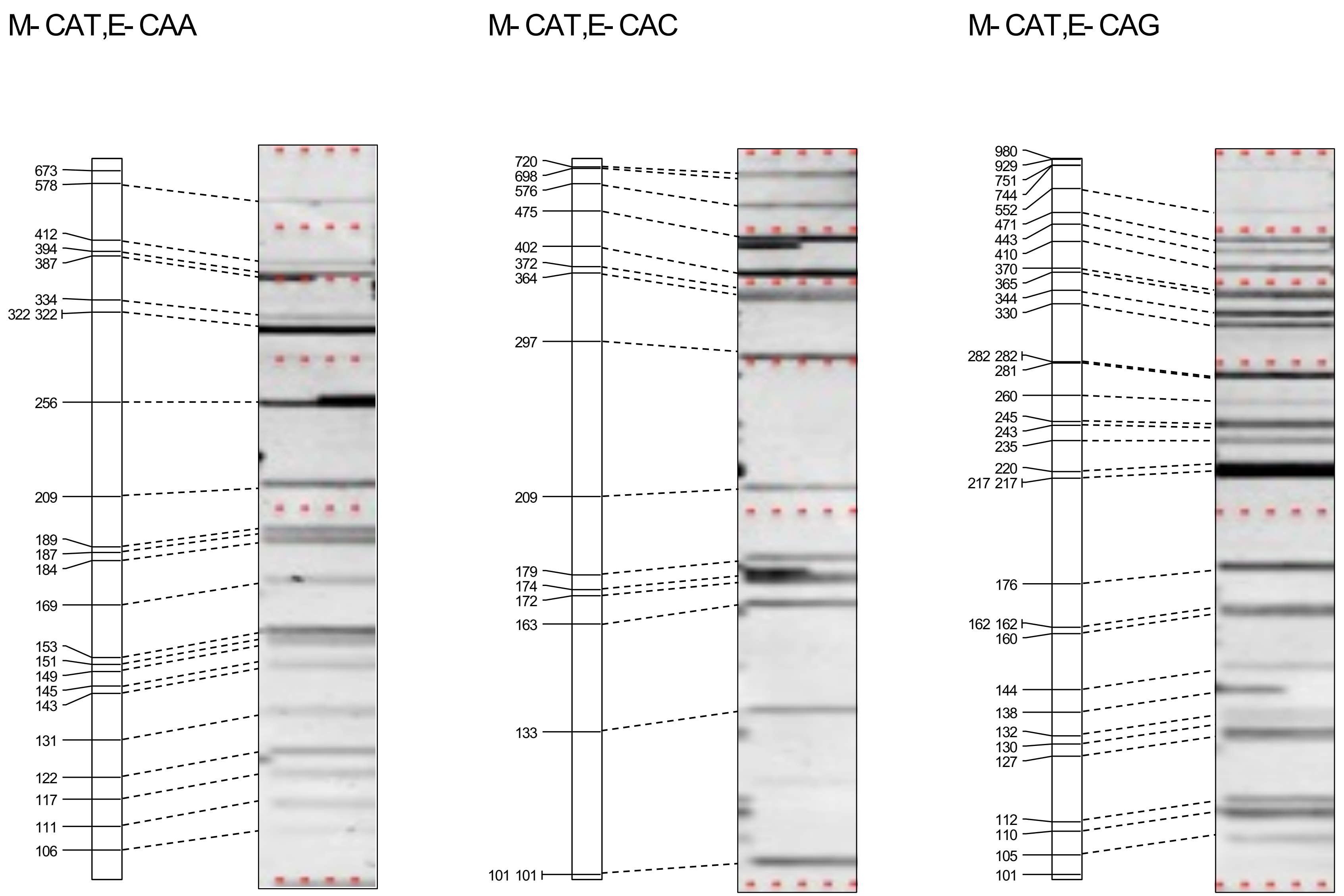


**Supplementary Figure S1 (continued)** *In silico* AFLP and experimental data.

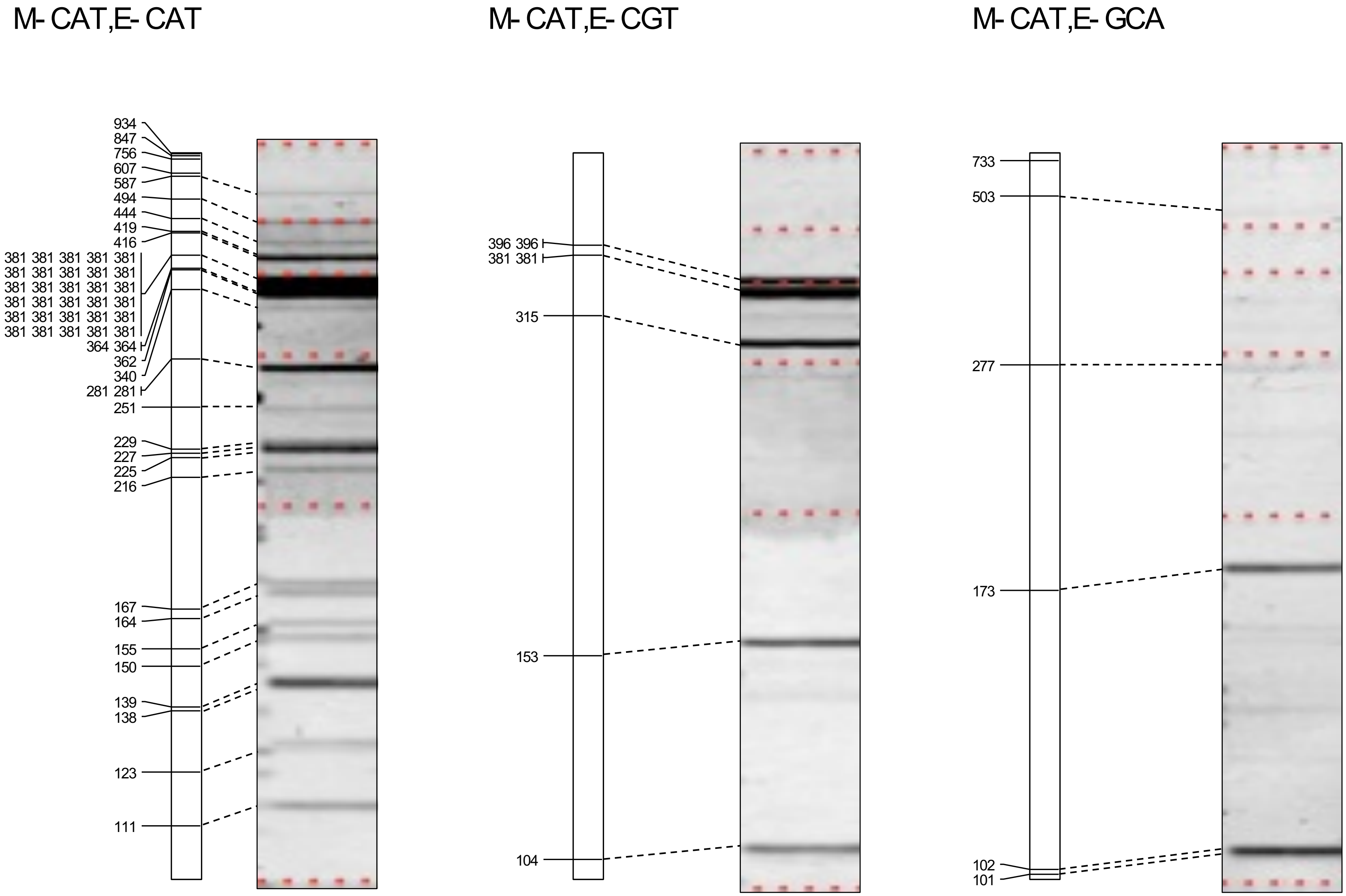


**Supplementary Figure S1 (continued)** *In silico* AFLP and experimental data.

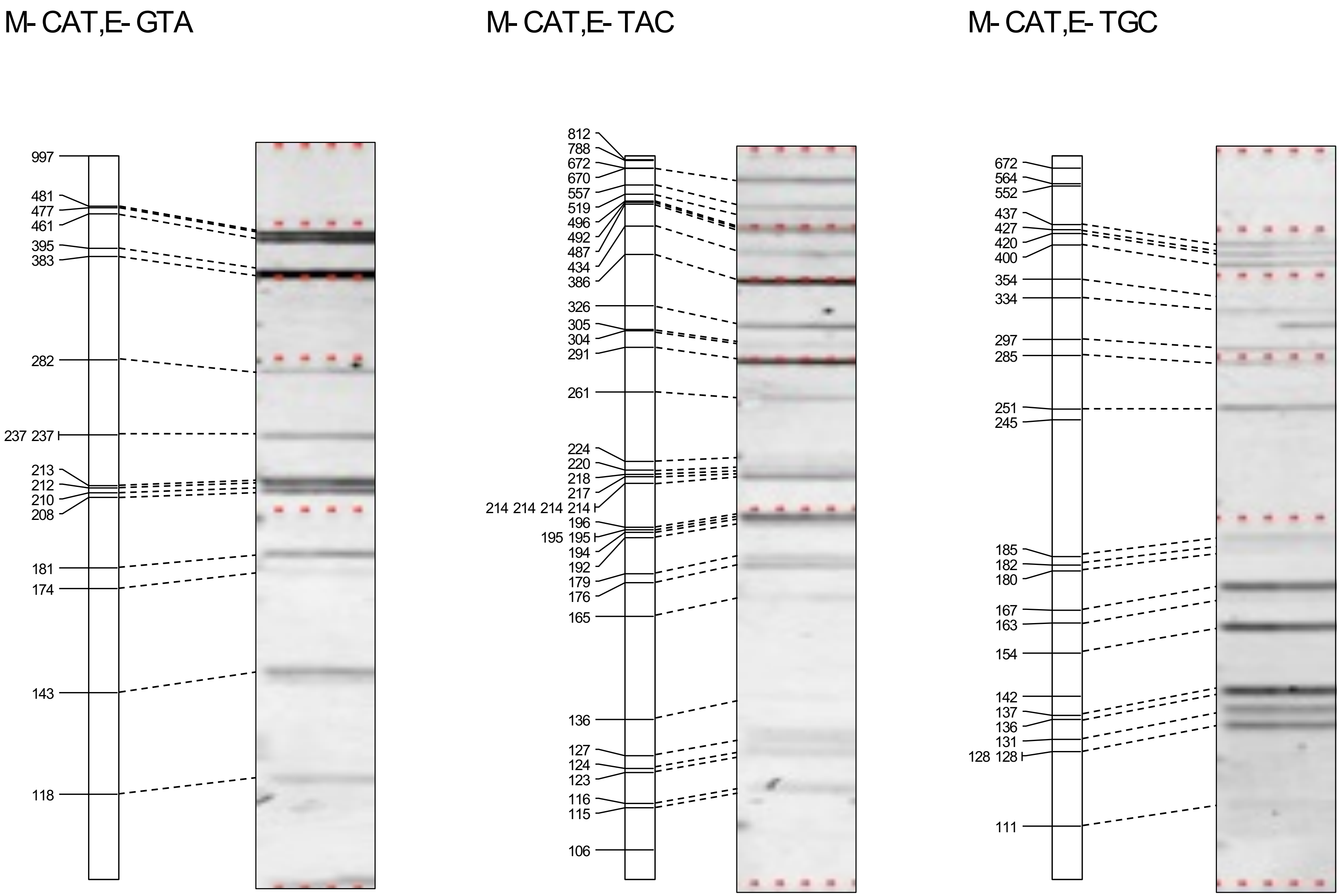


**Supplementary Figure S1 (continued)** *In silico* AFLP and experimental data.

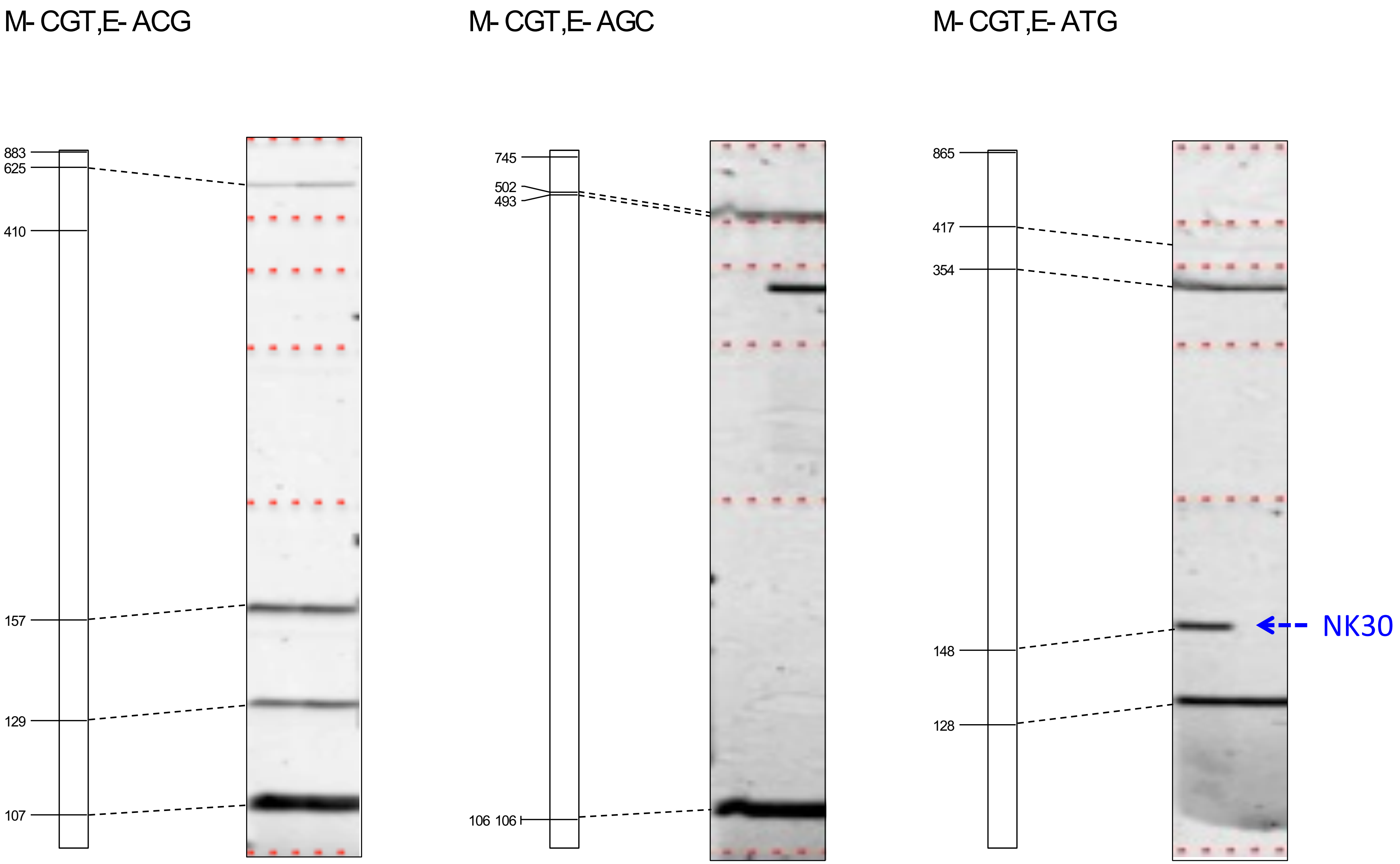


**Supplementary Figure S1 (continued)** *In silico* AFLP and experimental data.

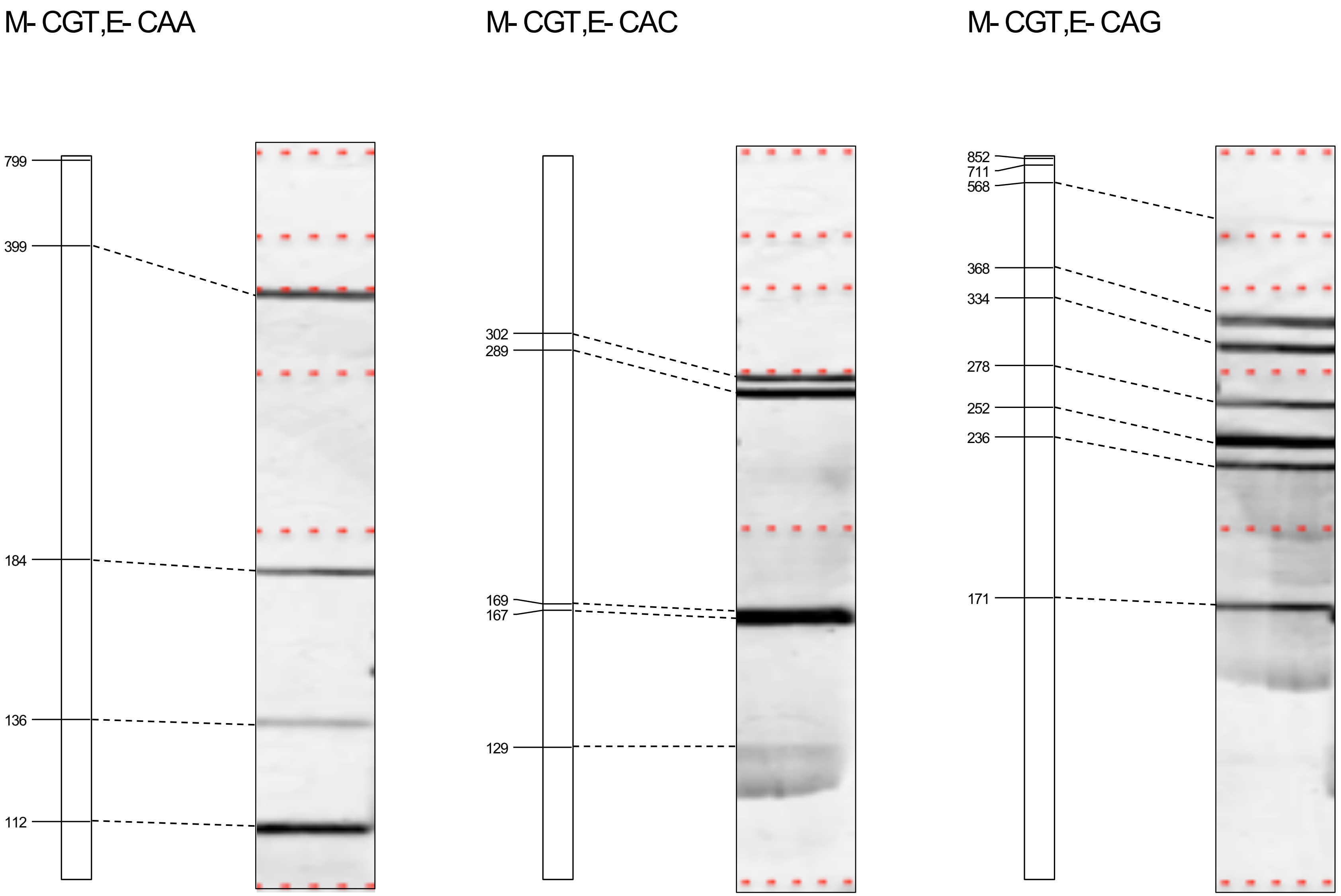


**Supplementary Figure S1 (continued)** *In silico* AFLP and experimental data.

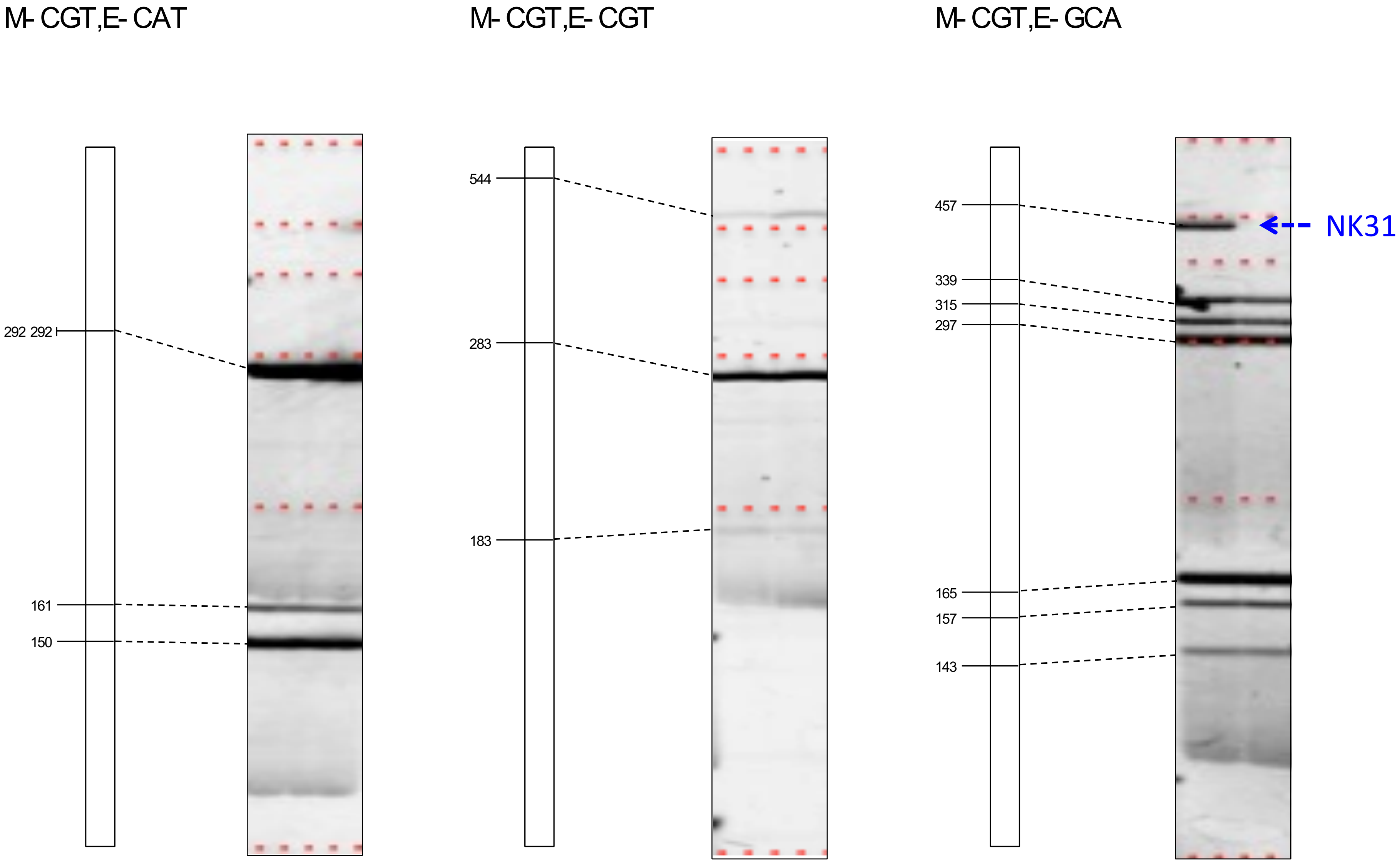


**Supplementary Figure S1 (continued)** *In silico* AFLP and experimental data.

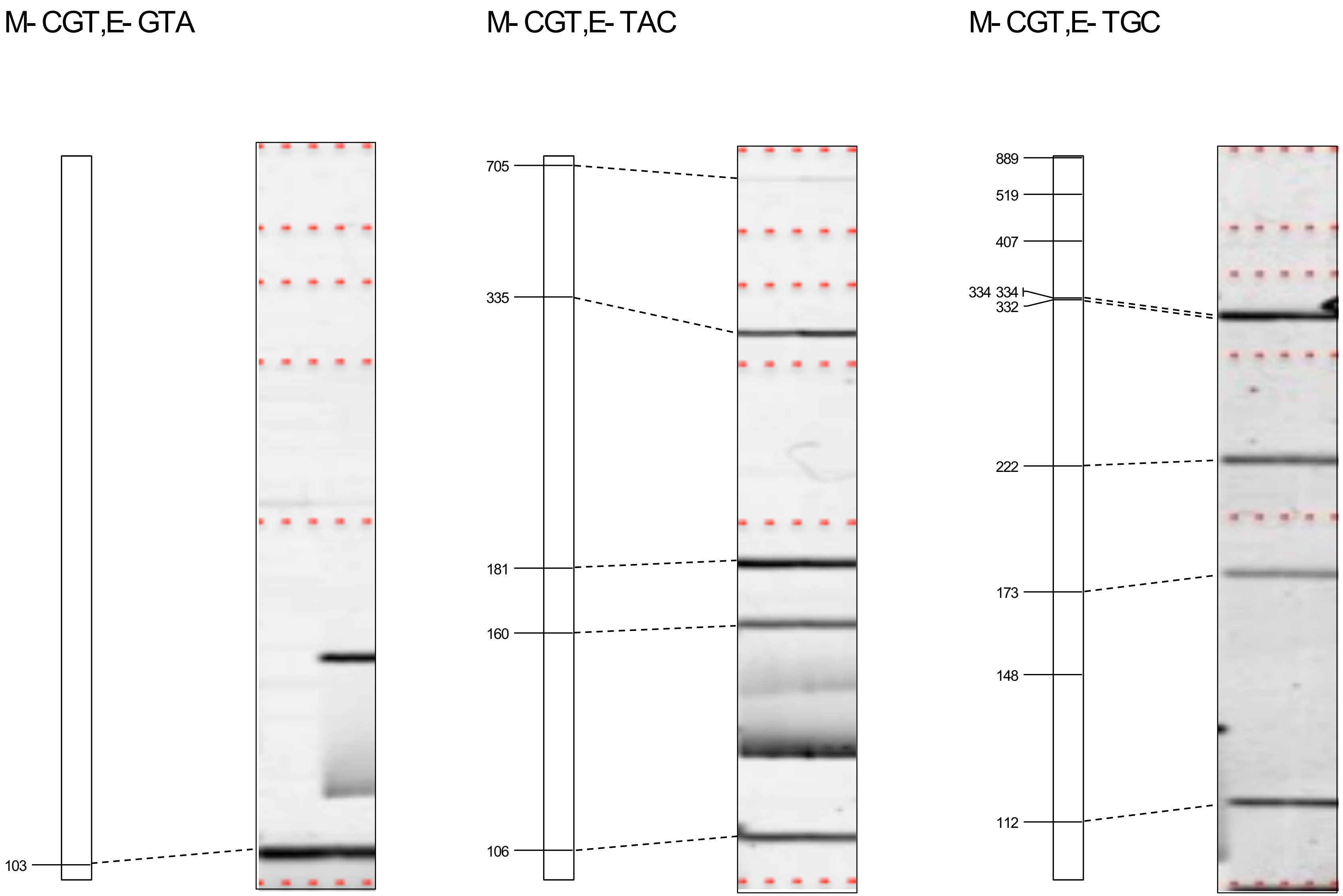


**Supplementary Figure S1 (continued)** *In silico* AFLP and experimental data.

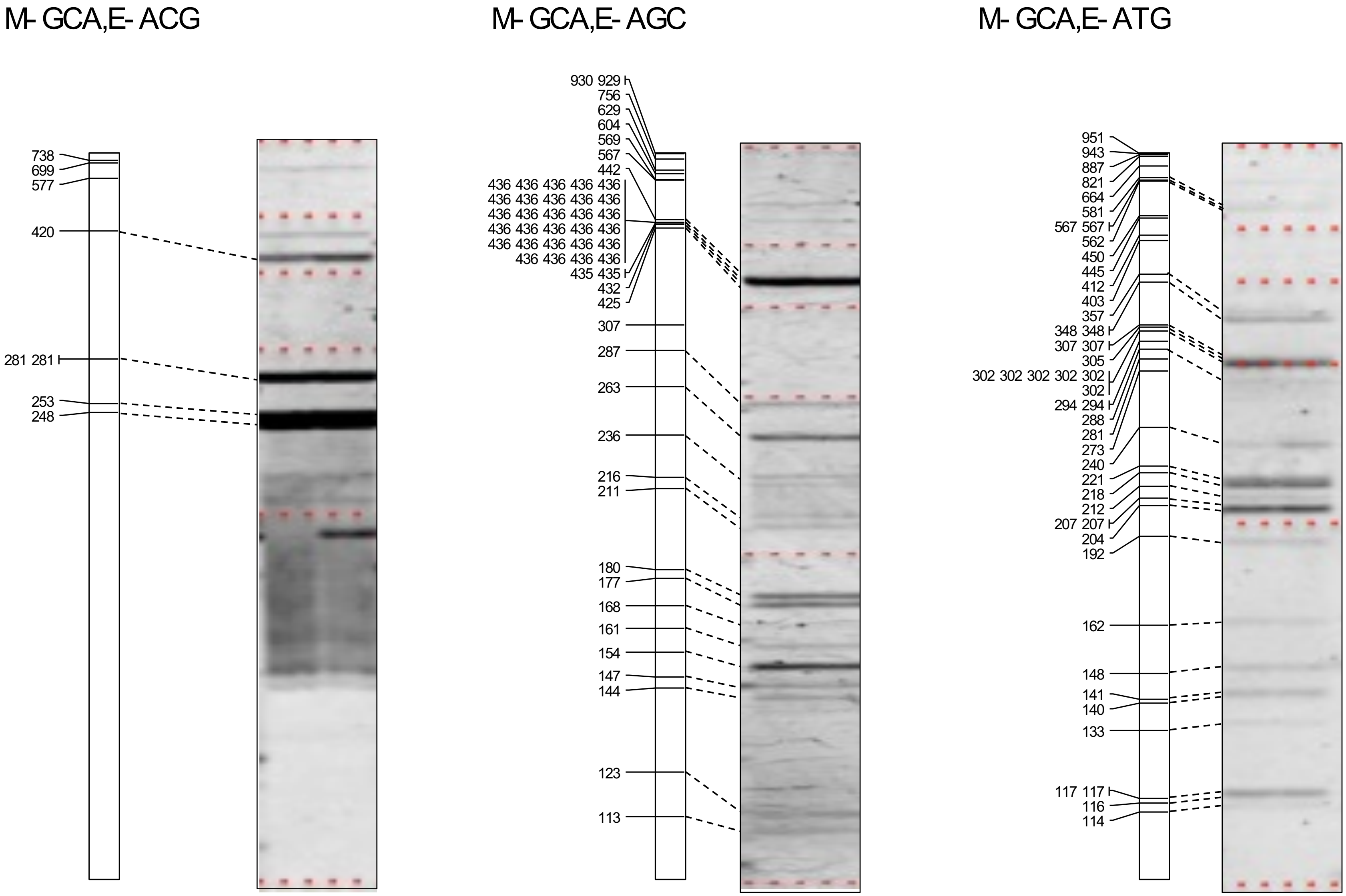


**Supplementary Figure S1 (continued)** *In silico* AFLP and experimental data.

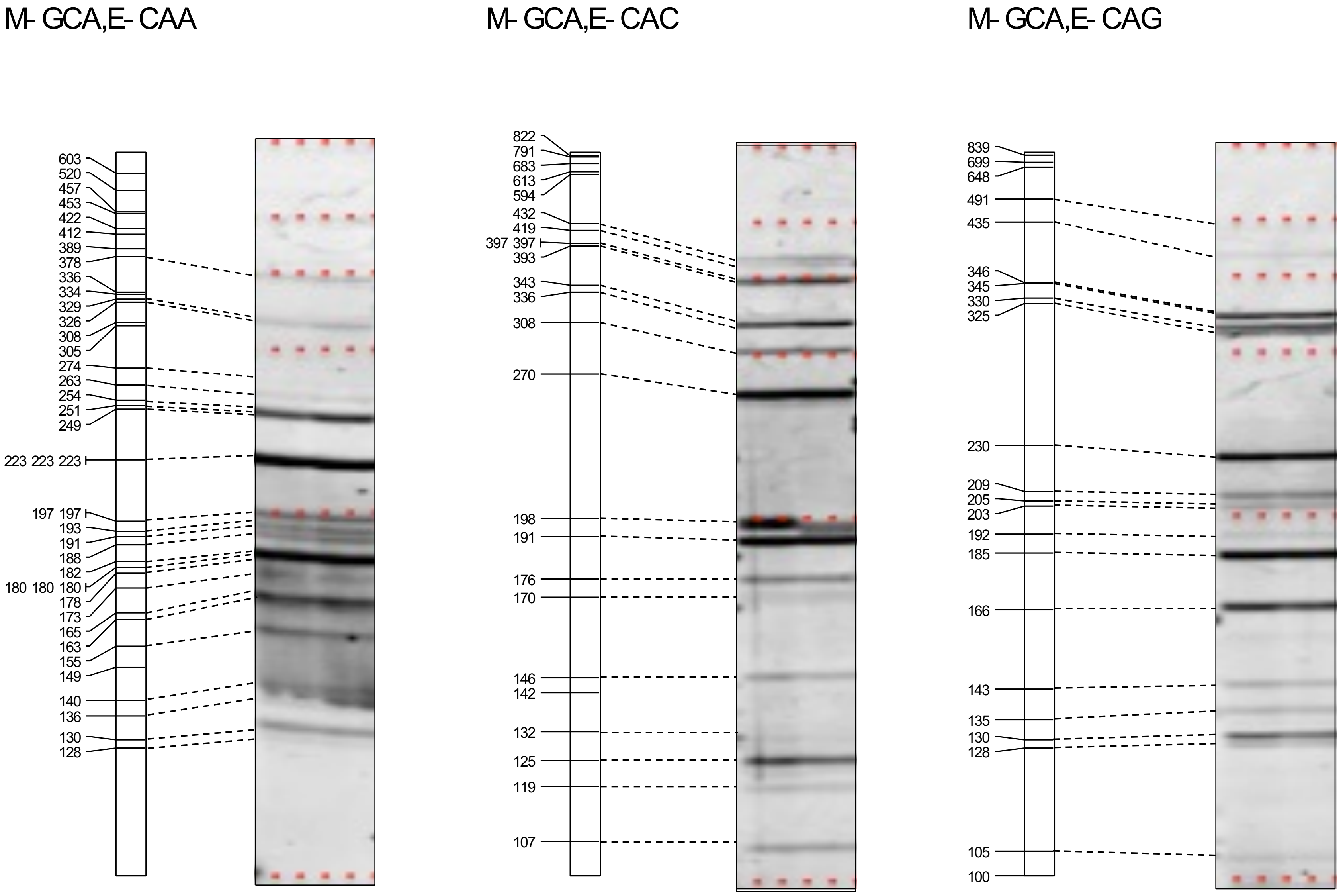


**Supplementary Figure S1 (continued)** *In silico* AFLP and experimental data.

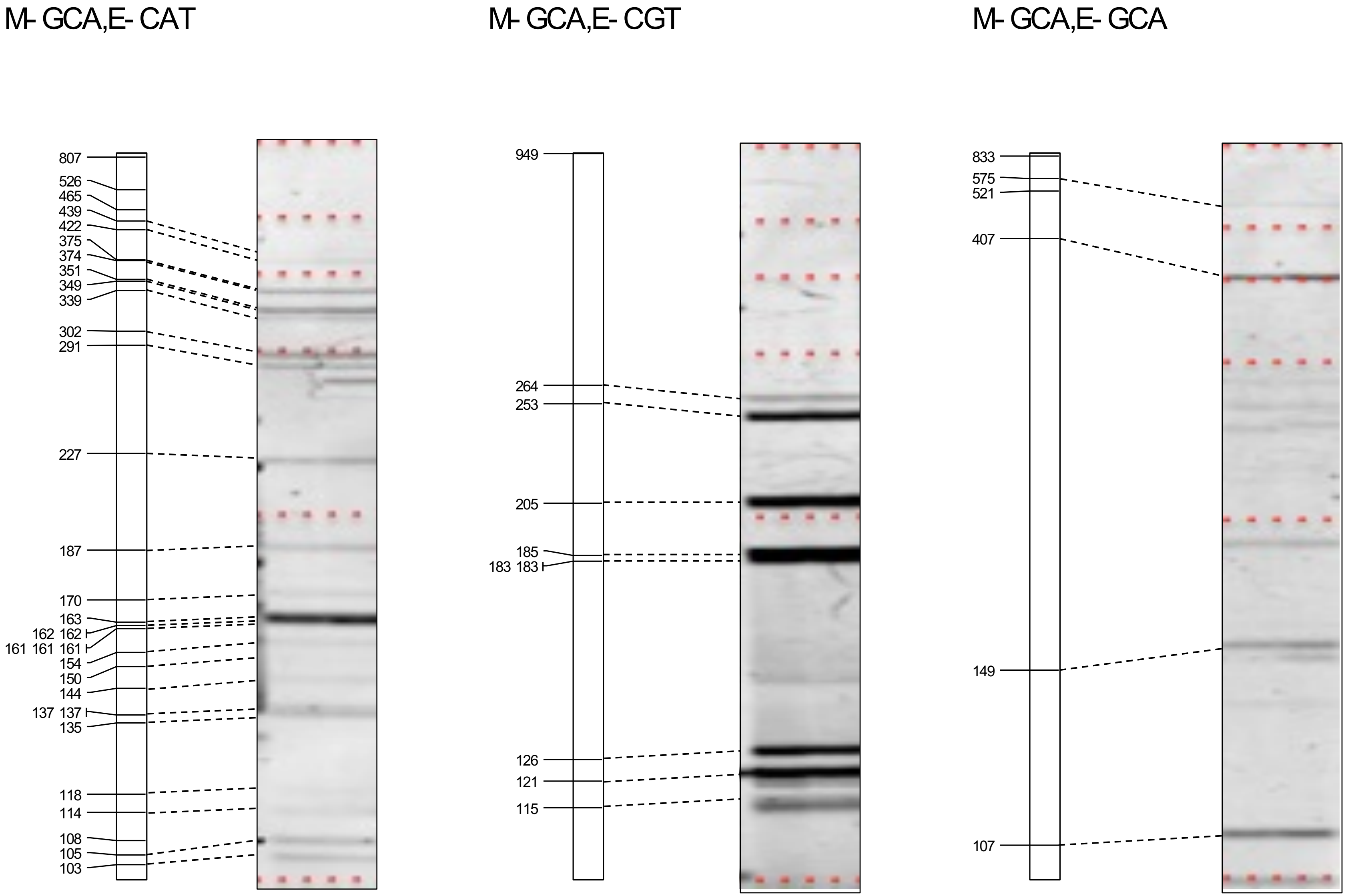


**Supplementary Figure S1 (continued)** *In silico* AFLP and experimental data.

M- GCA,E- GTA

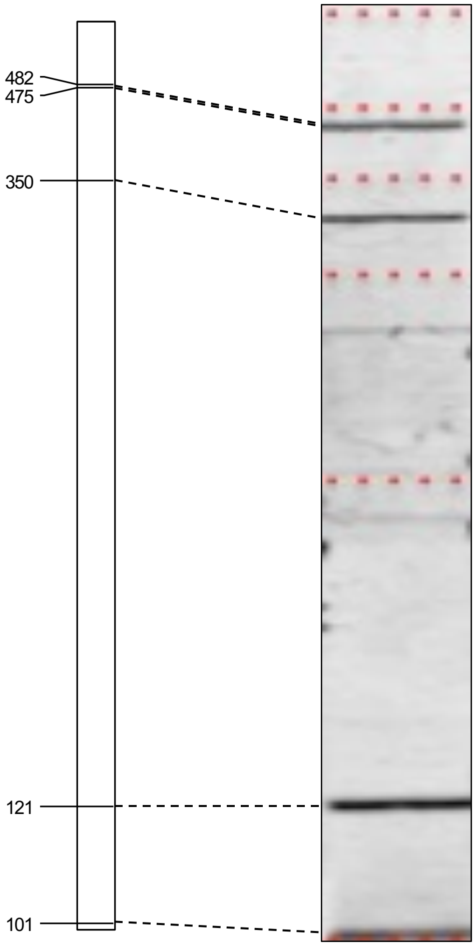


M- GCA,E- TAC

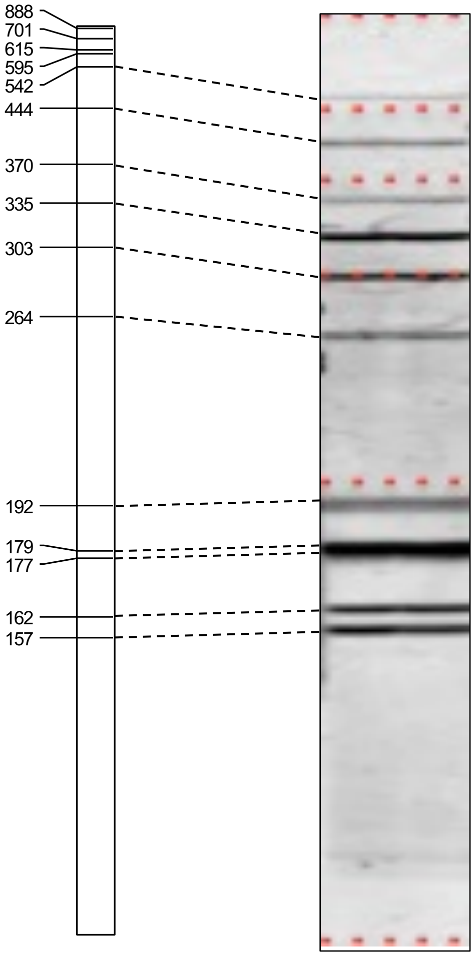


M- GCA,E- TGC

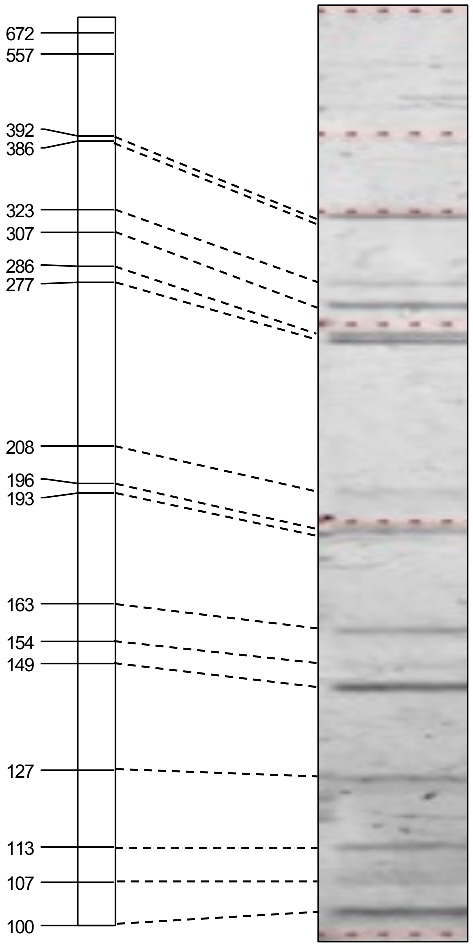


**Supplementary Figure S1 (continued)** *In silico* AFLP and experimental data.

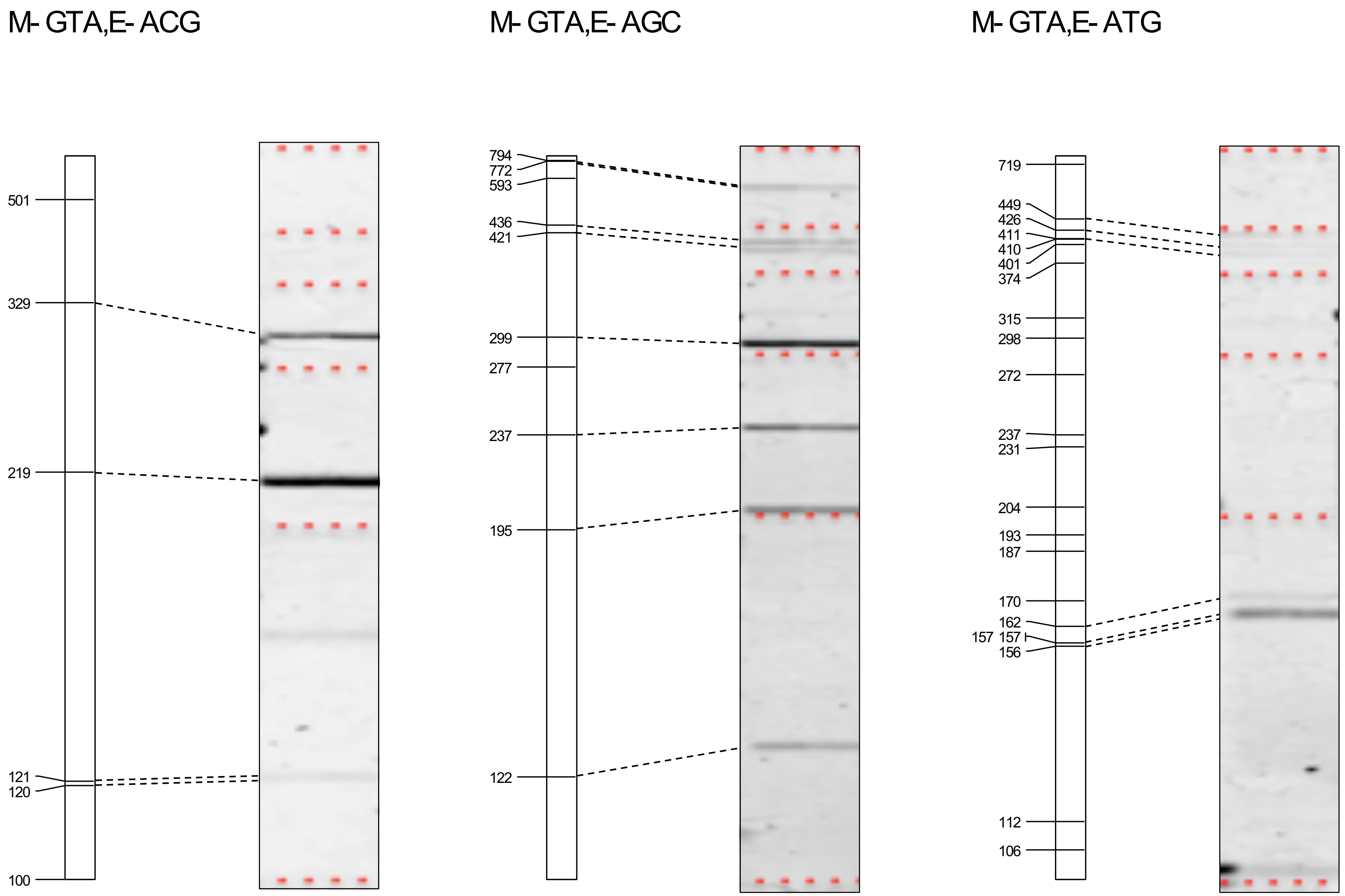


**Supplementary Figure S1 (continued)** *In silico* AFLP and experimental data.

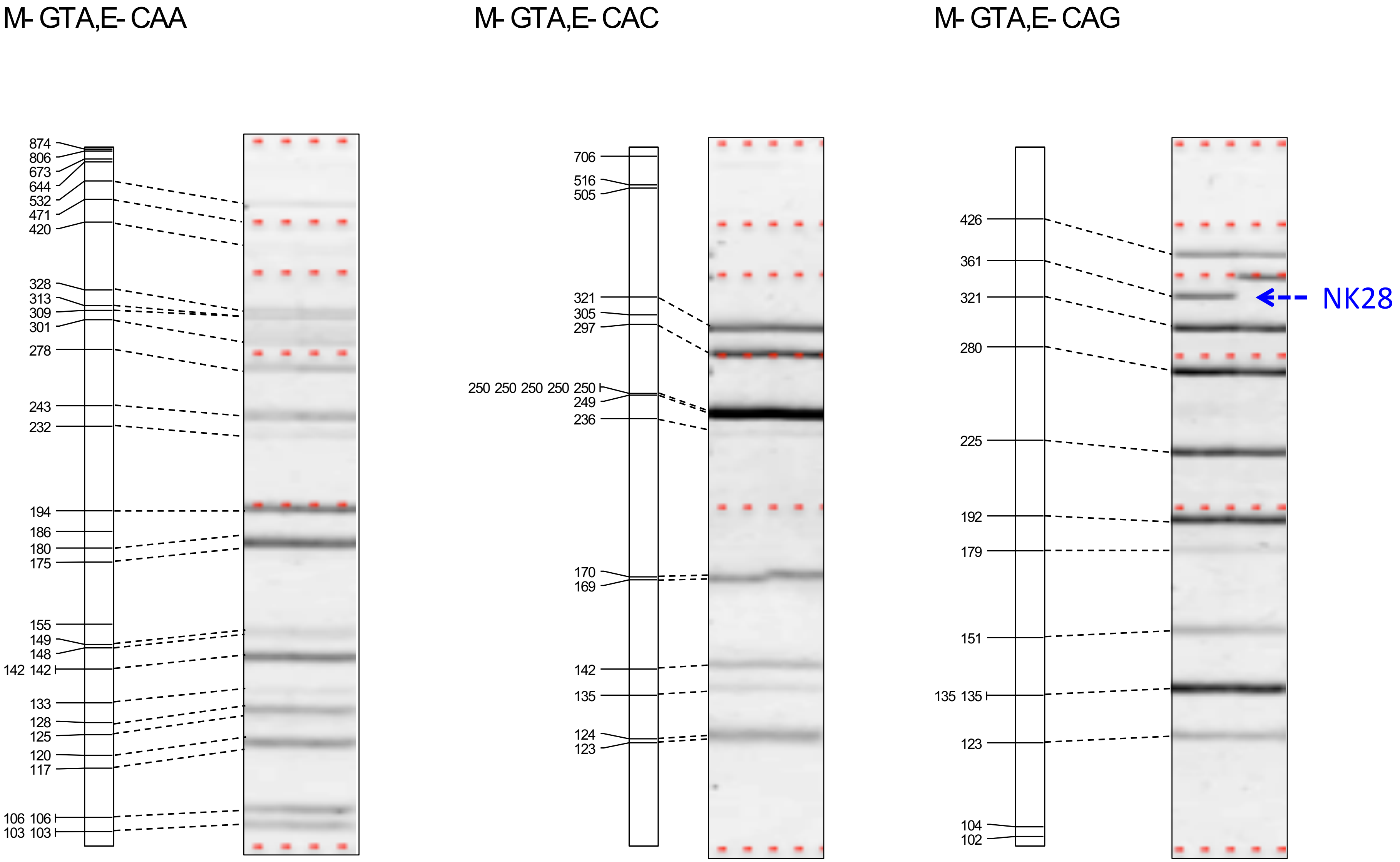


**Supplementary Figure S1 (continued)** *In silico* AFLP and experimental data.

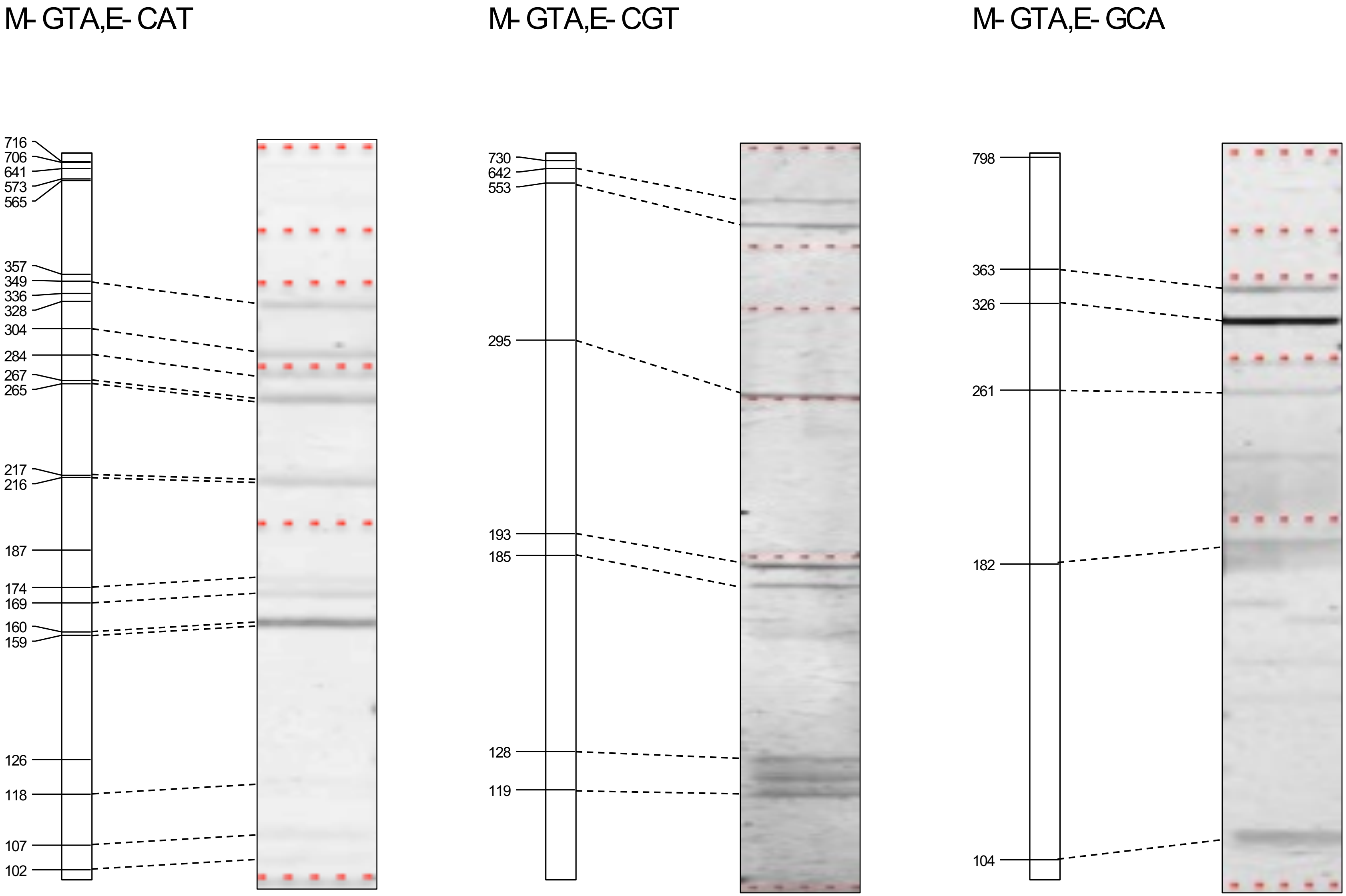


**Supplementary Figure S1 (continued)** *In silico* AFLP and experimental data.

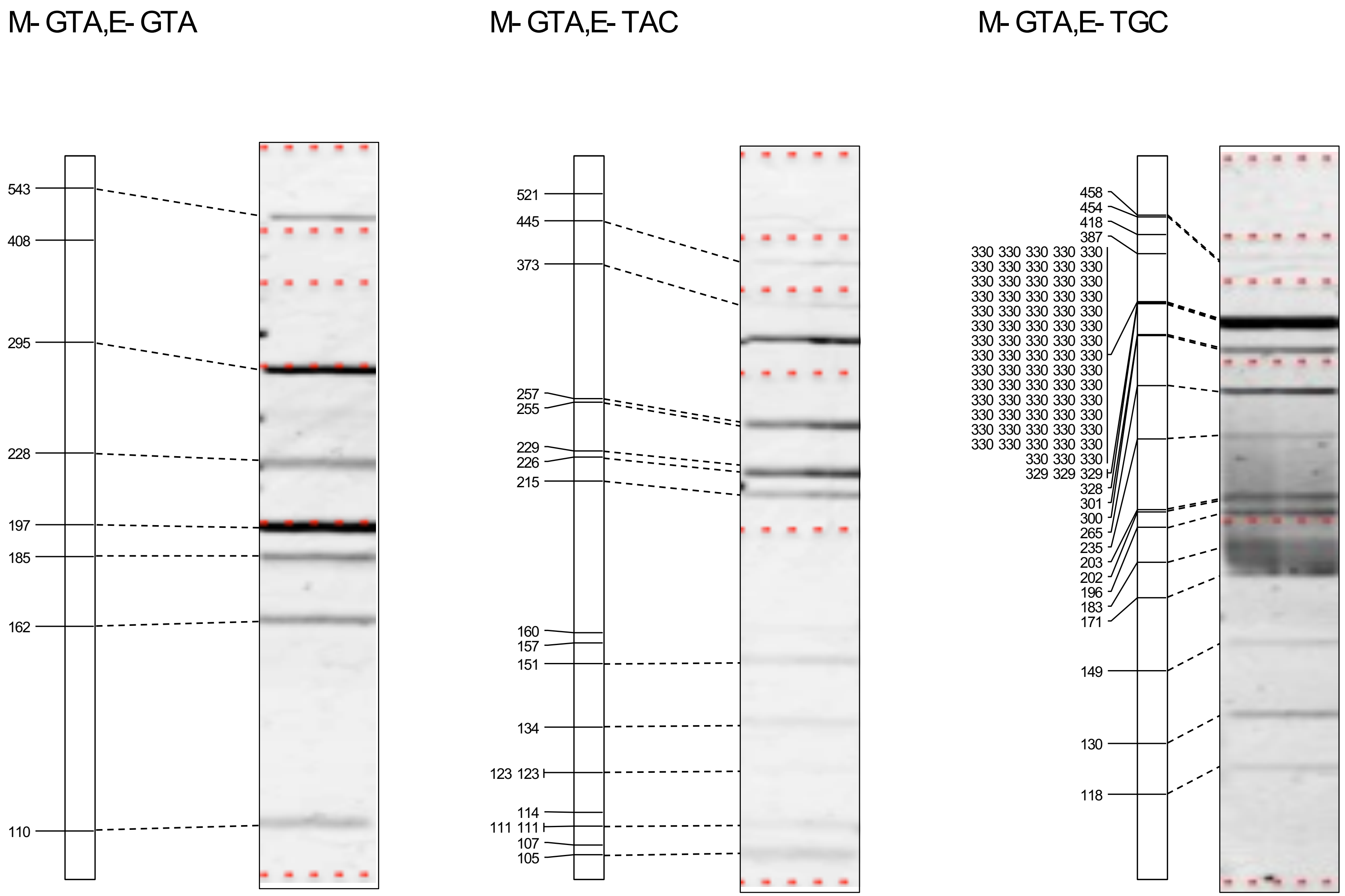


**Supplementary Figure S1 (continued)** *In silico* AFLP and experimental data.

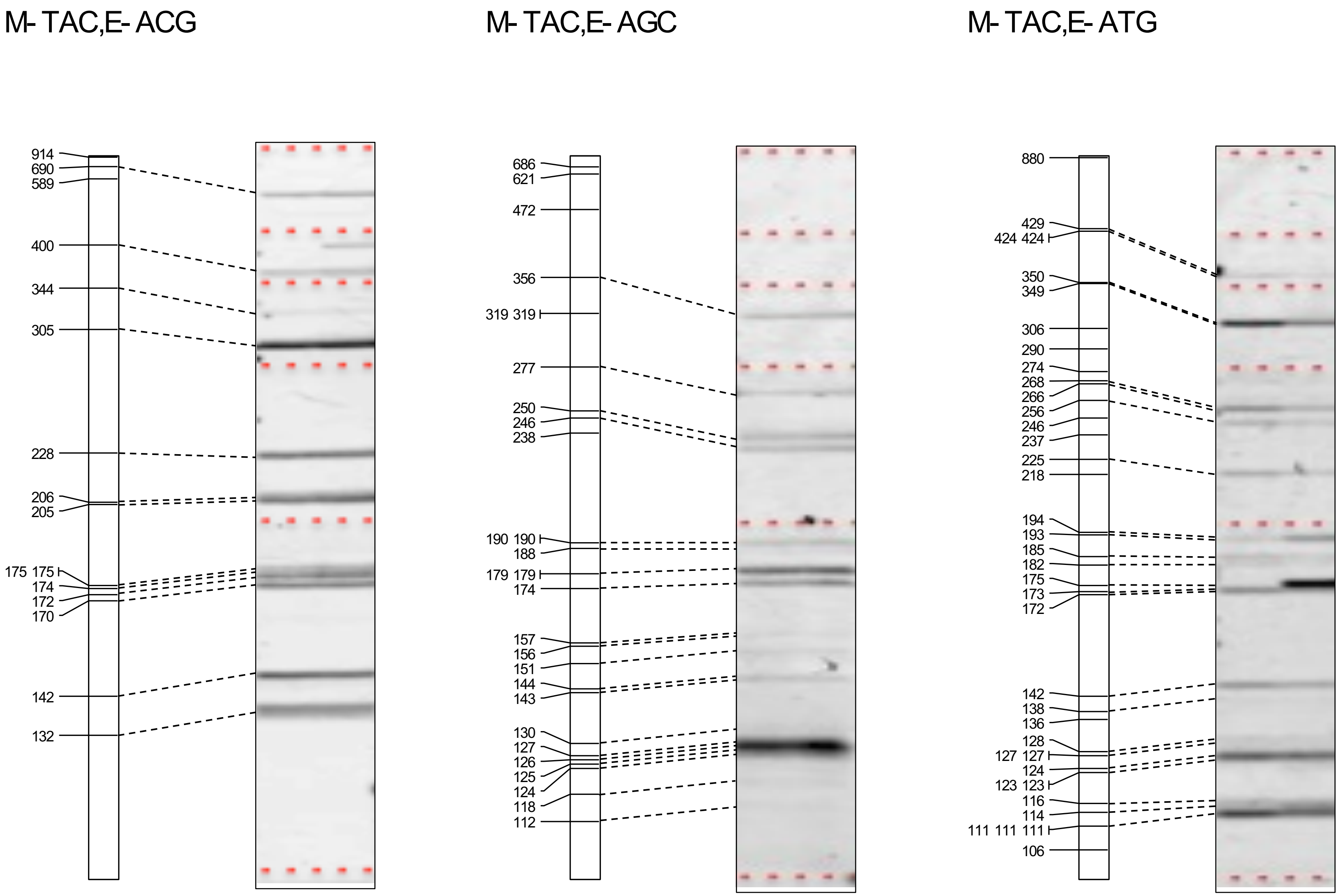


**Supplementary Figure S1 (continued)** *In silico* AFLP and experimental data.

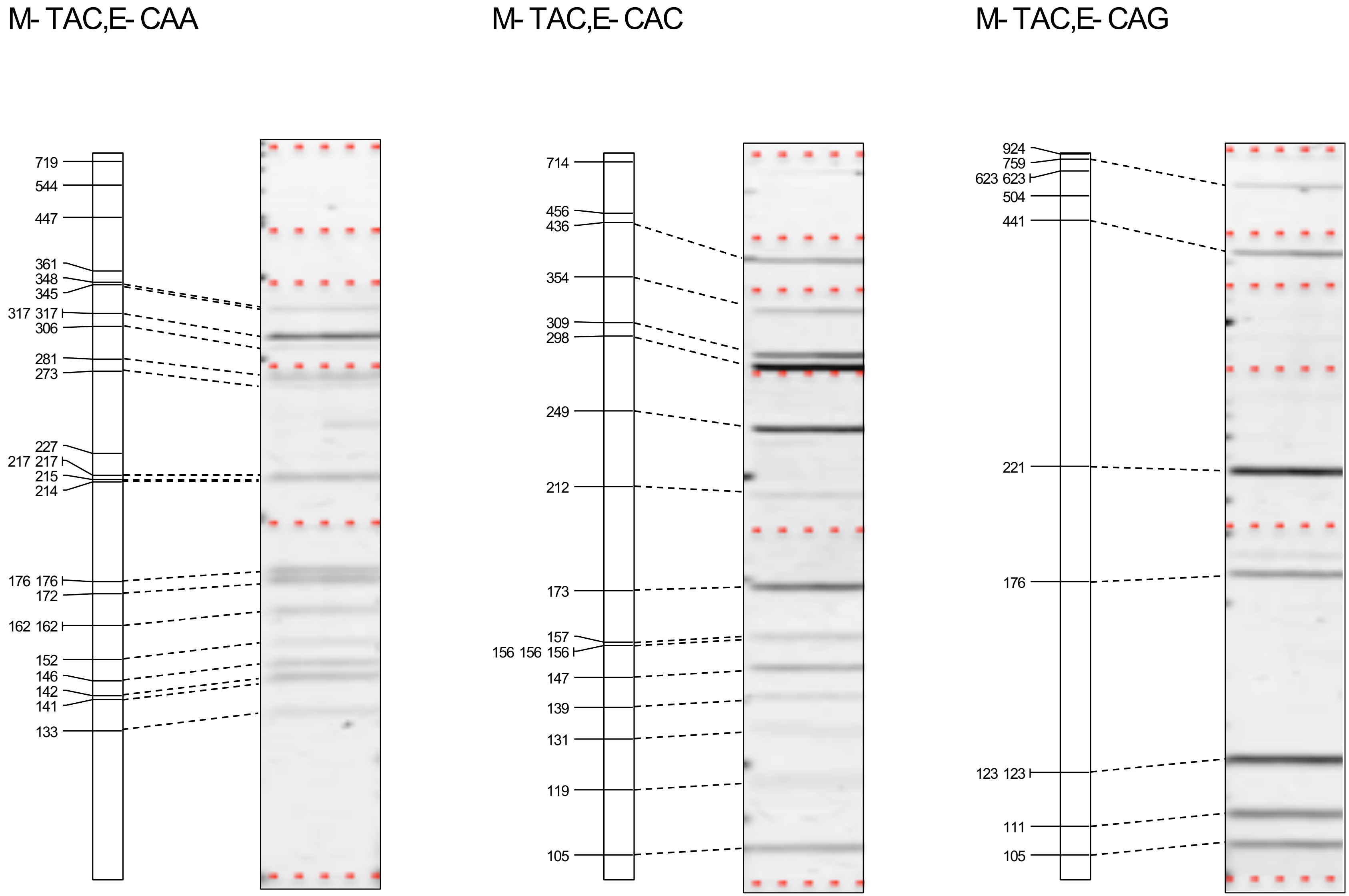


**Supplementary Figure S1 (continued)** *In silico* AFLP and experimental data.

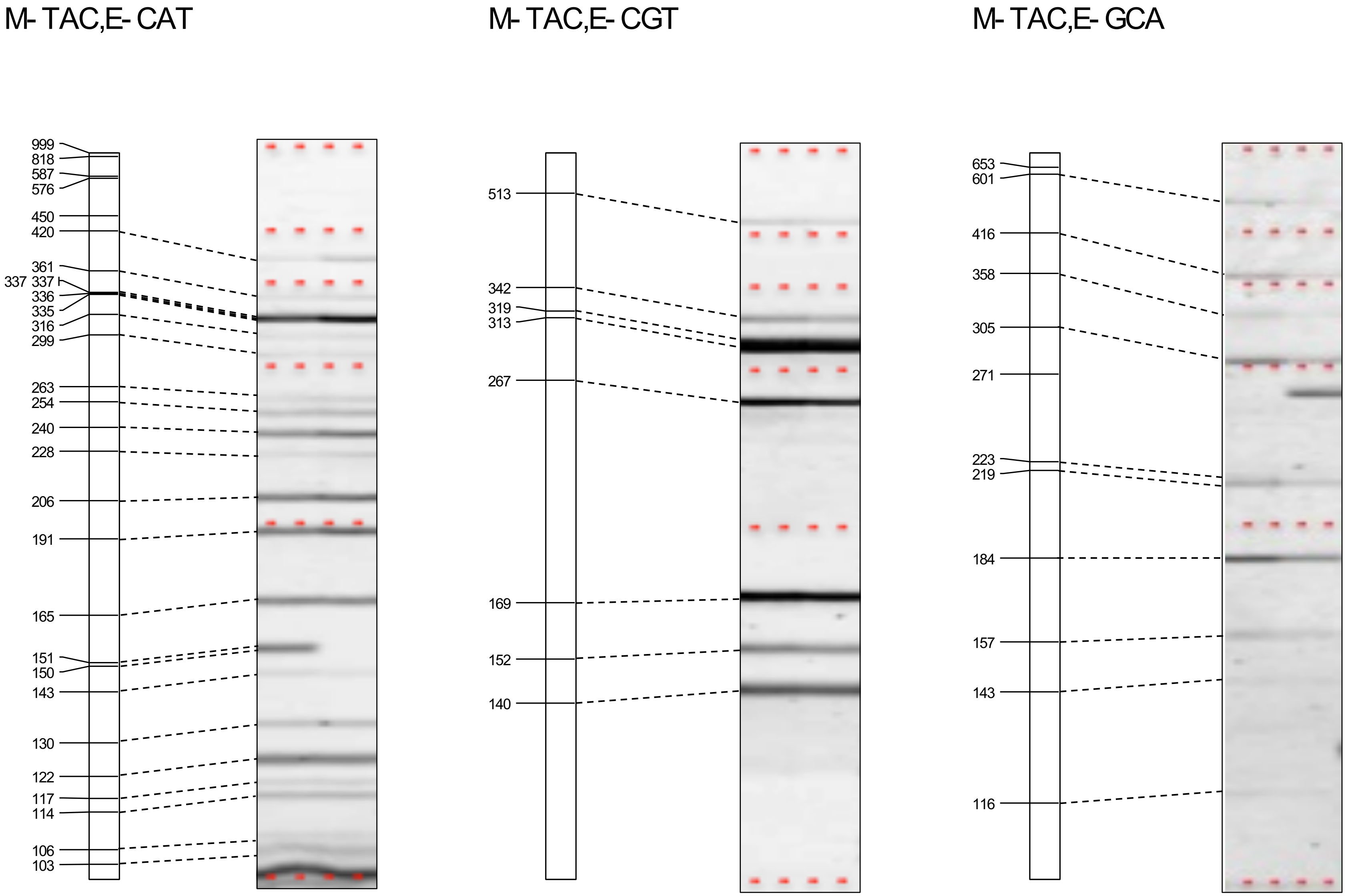


**Supplementary Figure S1 (continued)** *In silico* AFLP and experimental data.

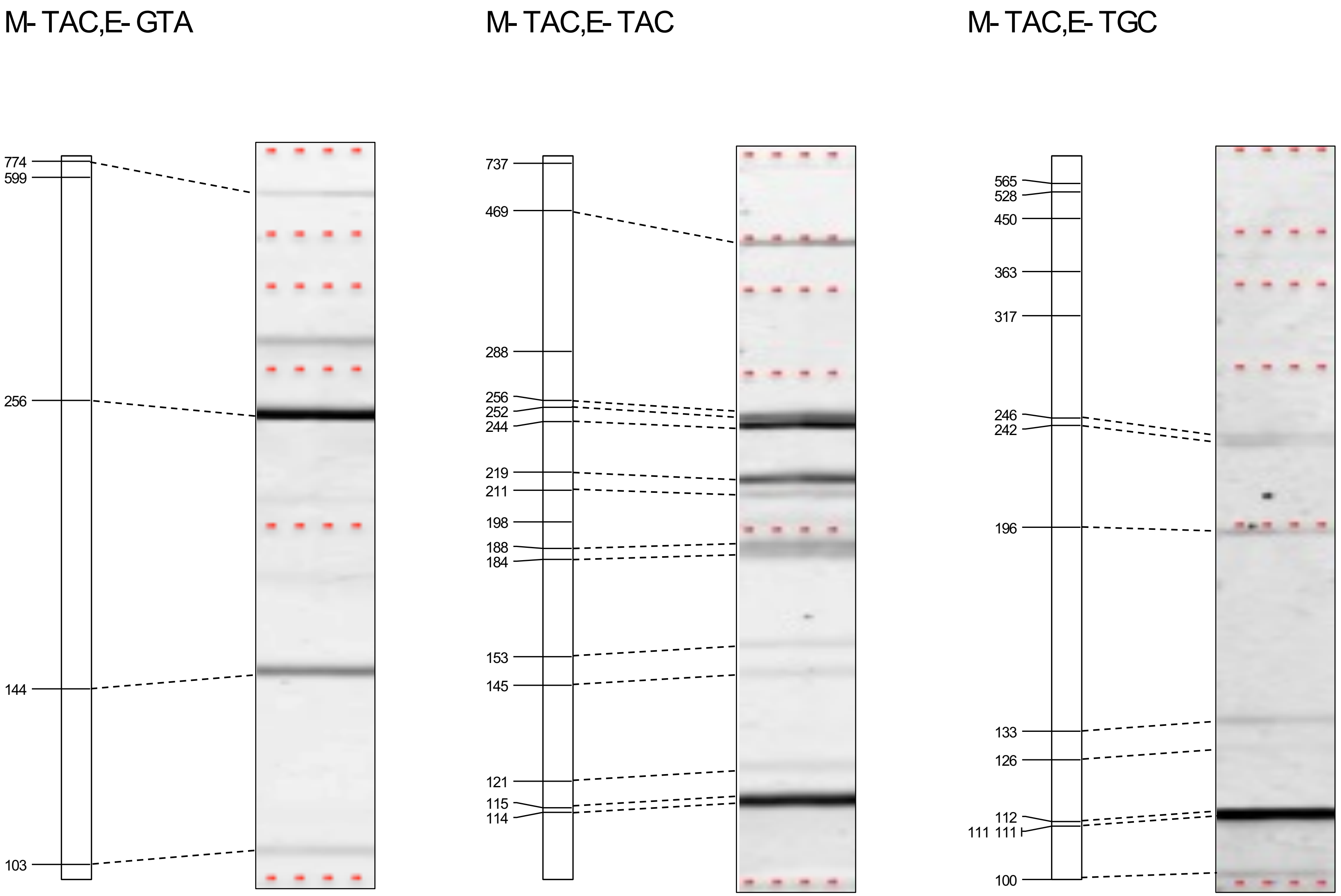


**Supplementary Figure S1 (continued)** *In silico* AFLP and experimental data.

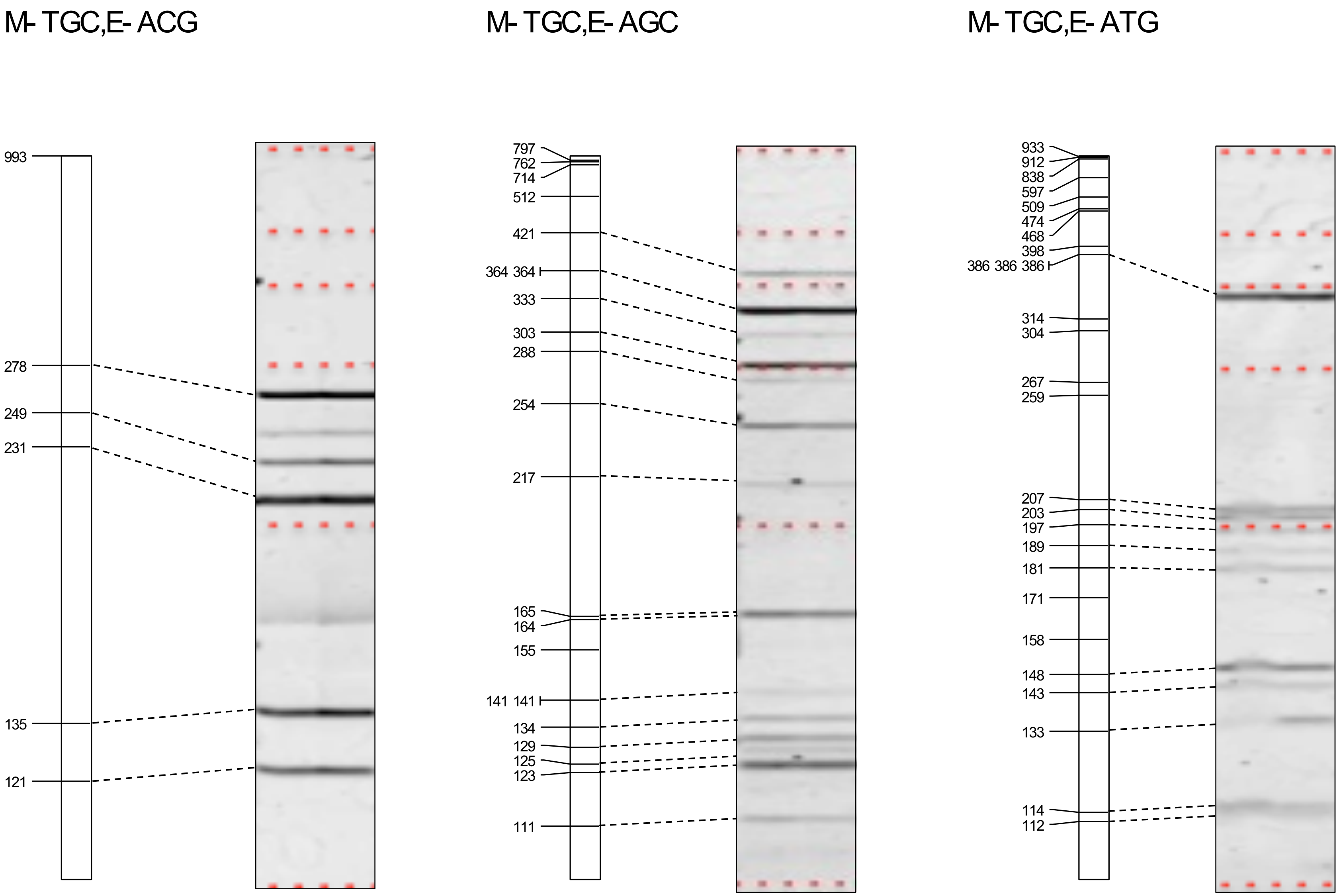


**Supplementary Figure S1 (continued)** *In silico* AFLP and experimental data.

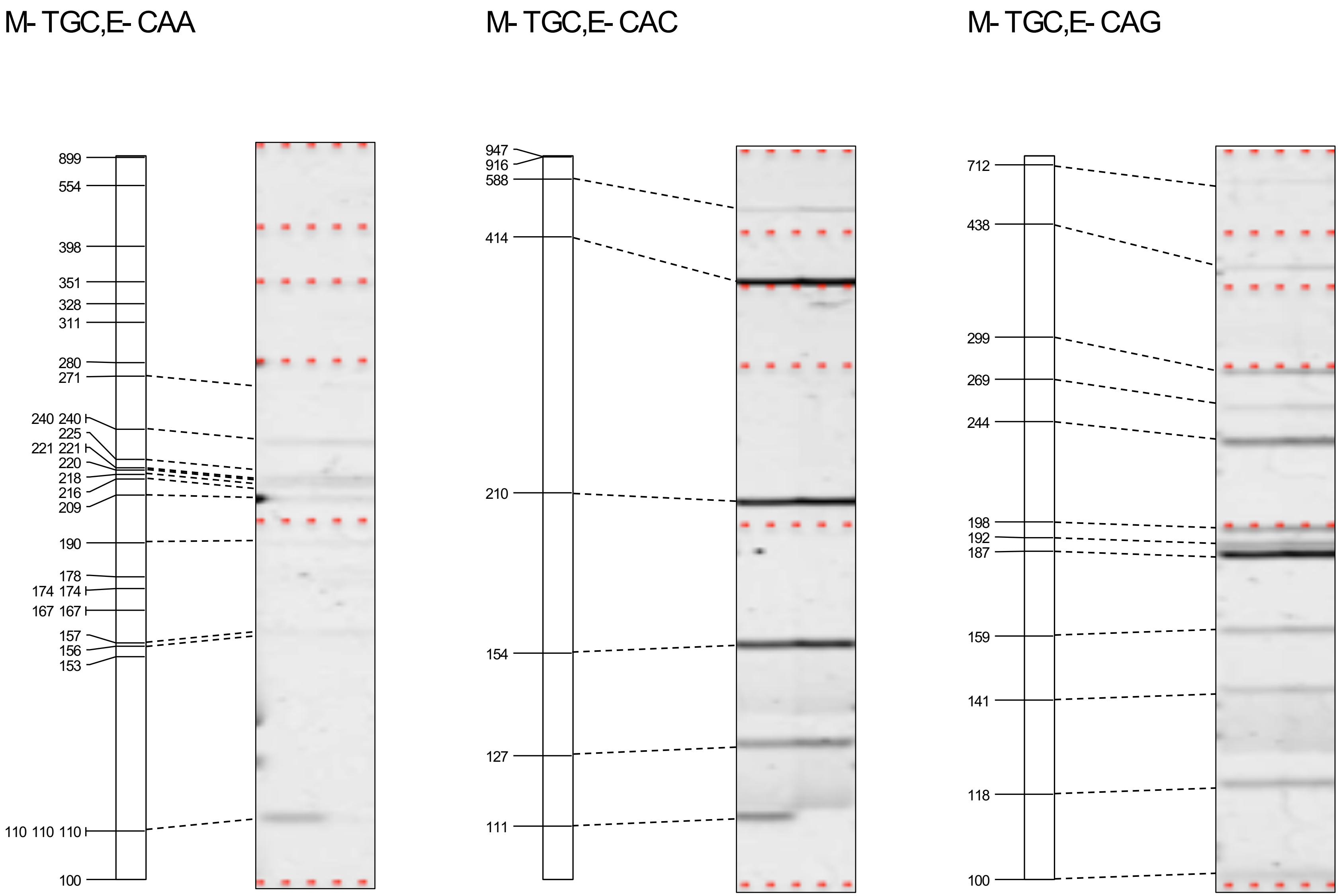


**Supplementary Figure S1 (continued)** *In silico* AFLP and experimental data.

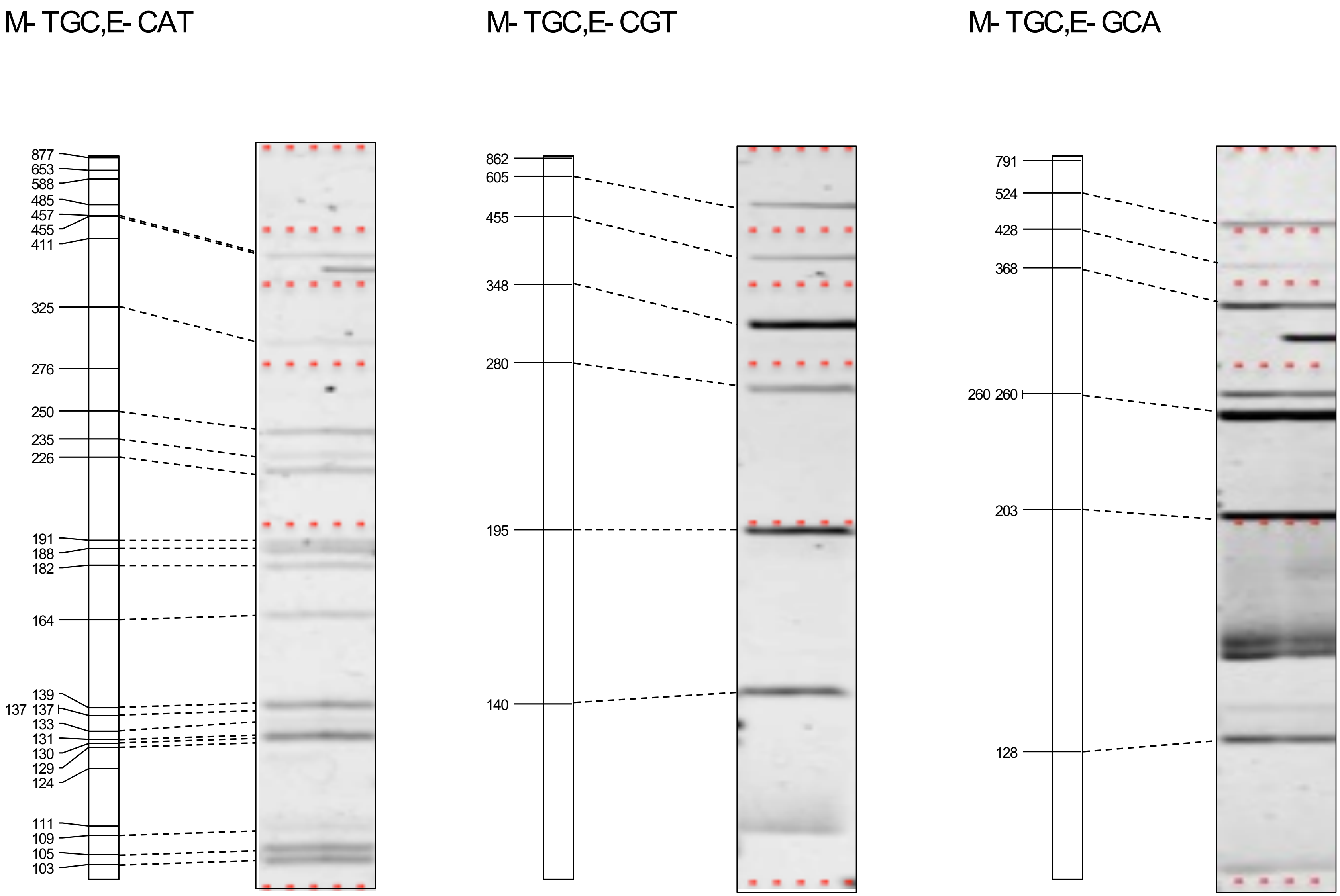


**Supplementary Figure S1 (continued)** *In silico* AFLP and experimental data.

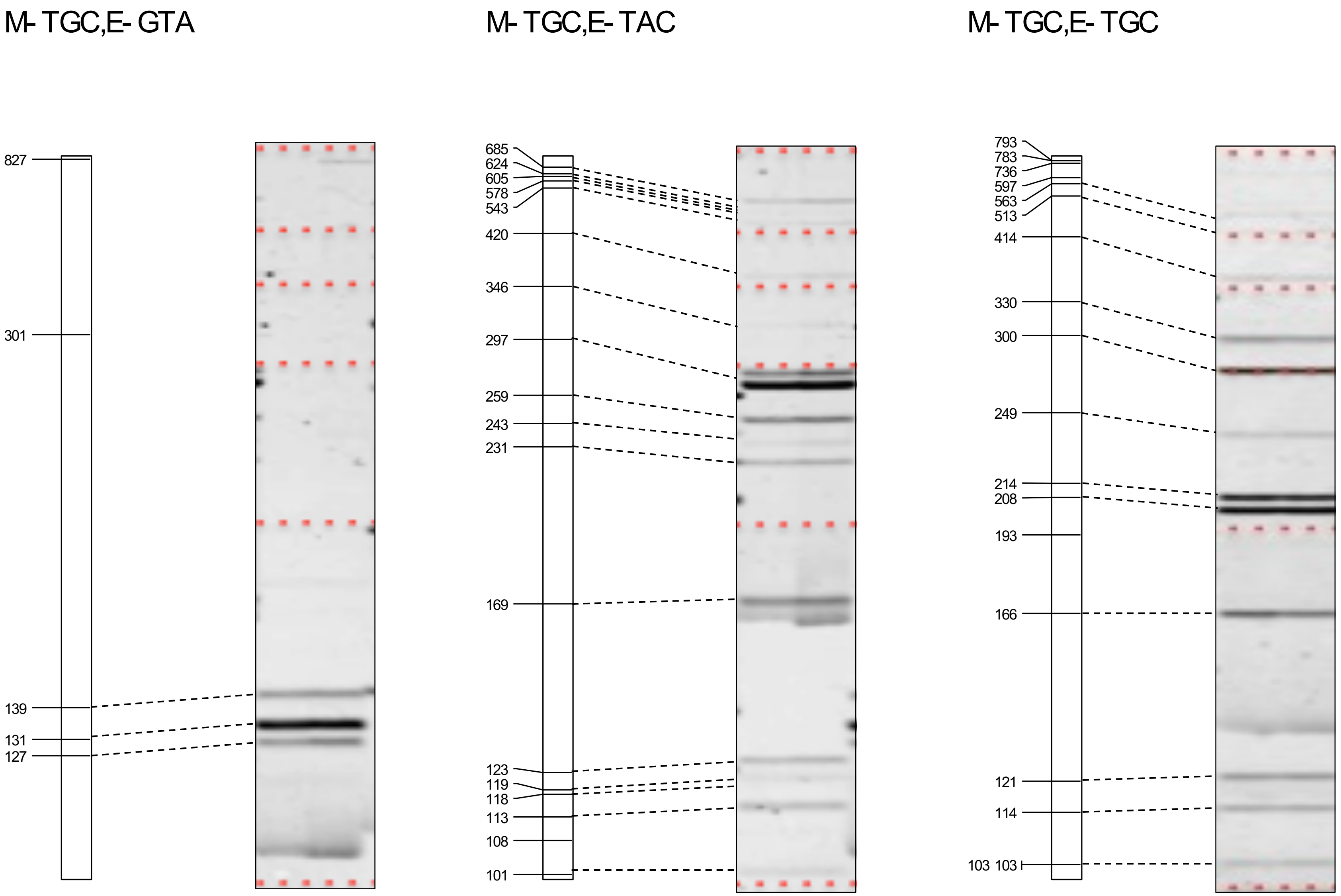


**Supplementary Figure S1 (continued)** *In silico* AFLP and experimental data.
