## Supplementary Figures S2 - S17 for "Optimizing selection of restriction enzymes for complexity-reduced genome sequencing in plants"

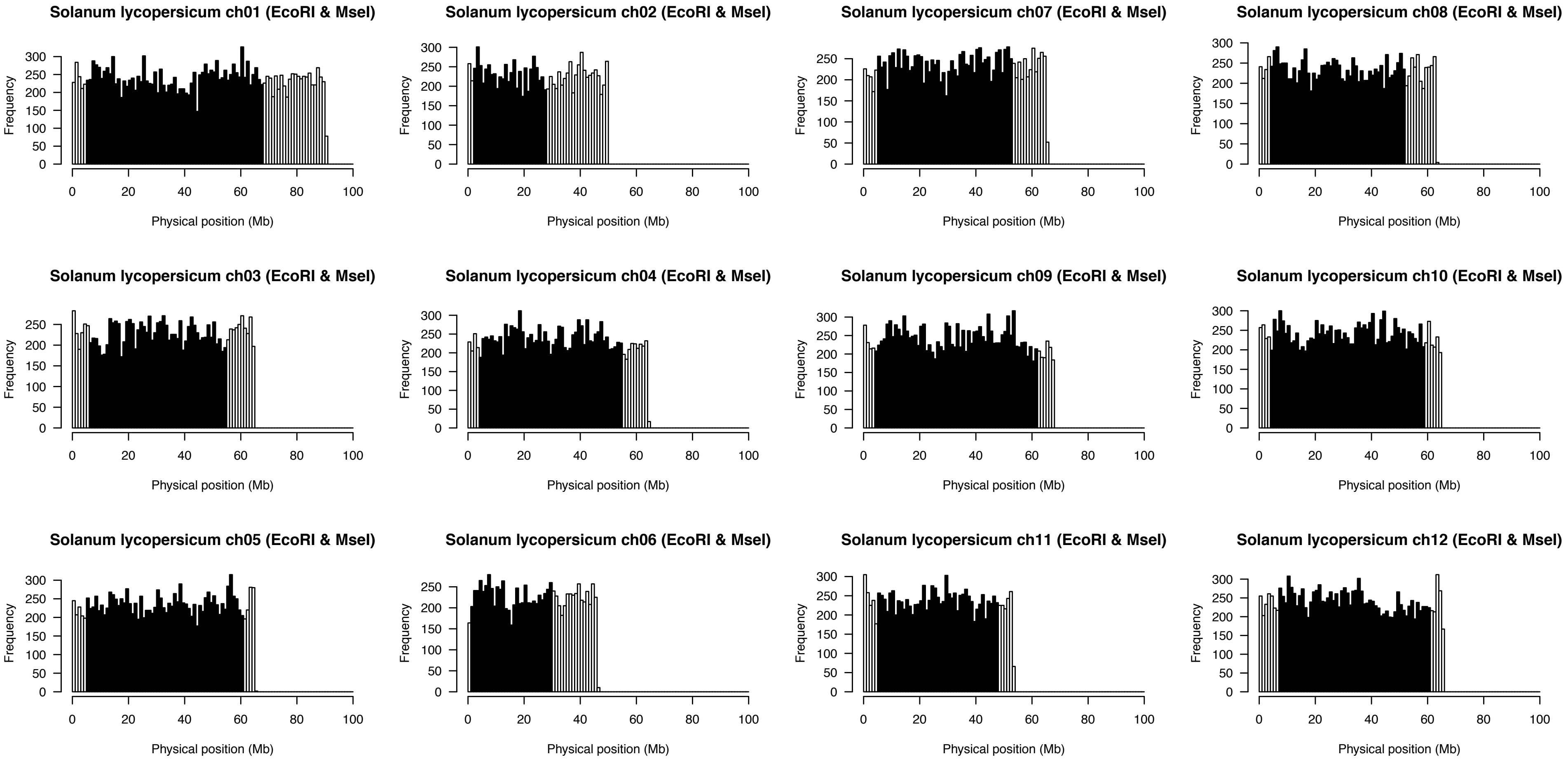


**Supplementary Figure S2** Distributions of the numbers of restriction fragments digested with EcoRI and MseI over the tomato genome. The non-overlapping window size is 1 Mb. Black and white bars indicate hetero- and euchromatic regions.

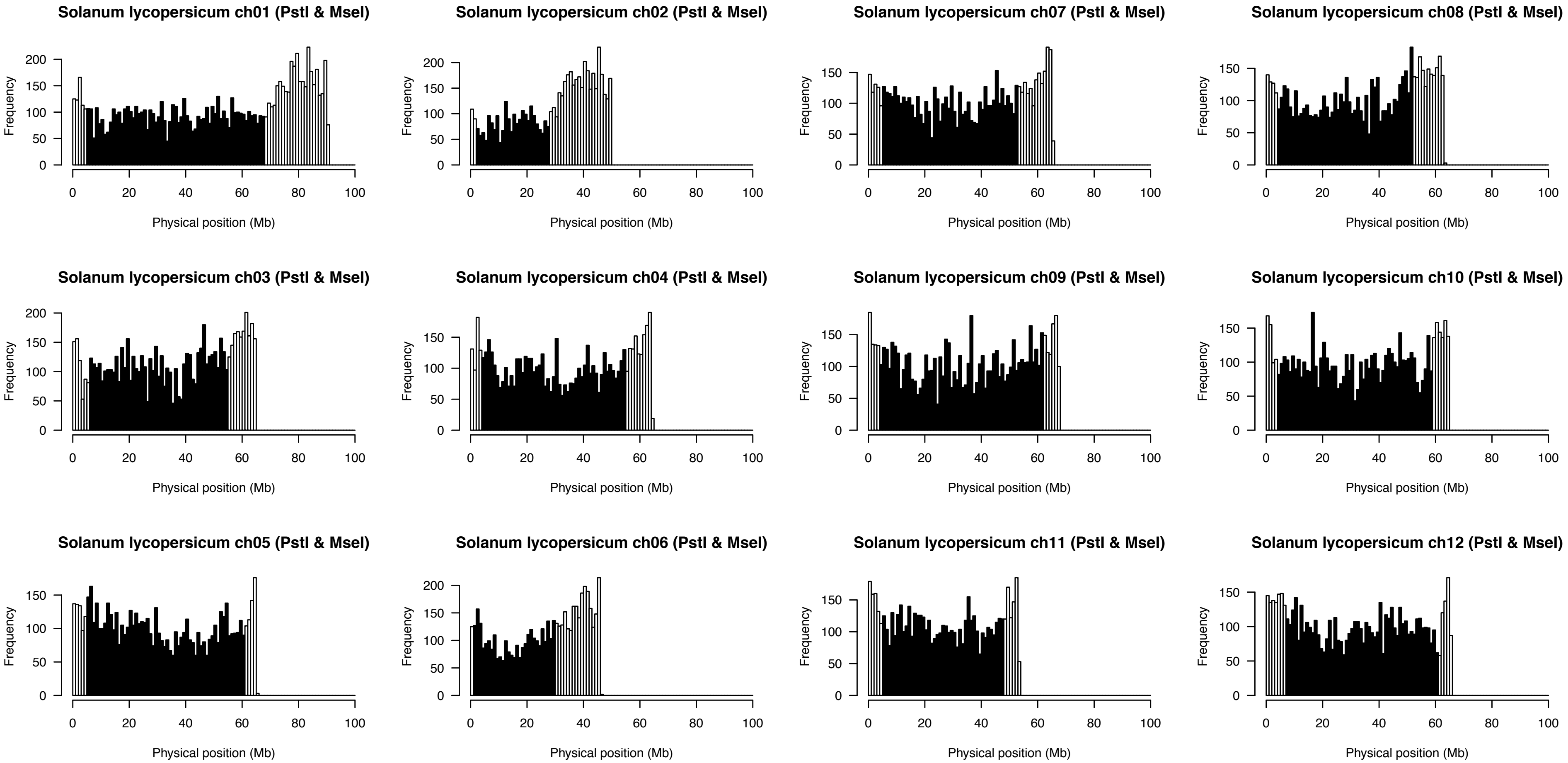


**Supplementary Figure S3** Distributions of the numbers of restriction fragments digested with PstI and MseI over the tomato genome. The non-overlapping window size is 1 Mb. Black and white bars indicate hetero- and euchromatic regions.

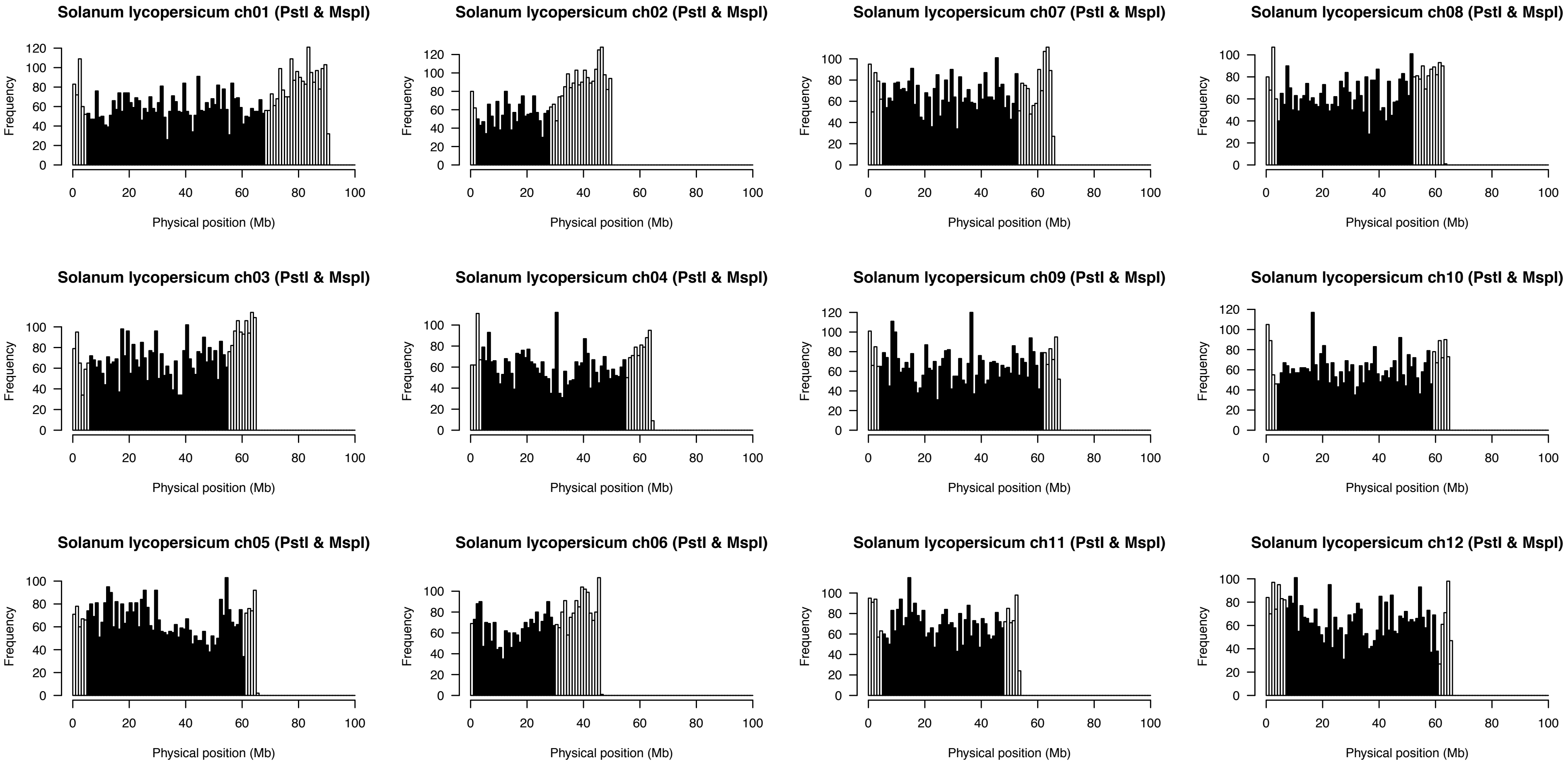


**Supplementary Figure S4** Distributions of the numbers of restriction fragments digested with PstI and MspI over the tomato genome. The non-overlapping window size is 1 Mb. Black and white bars indicate hetero- and euchromatic regions.

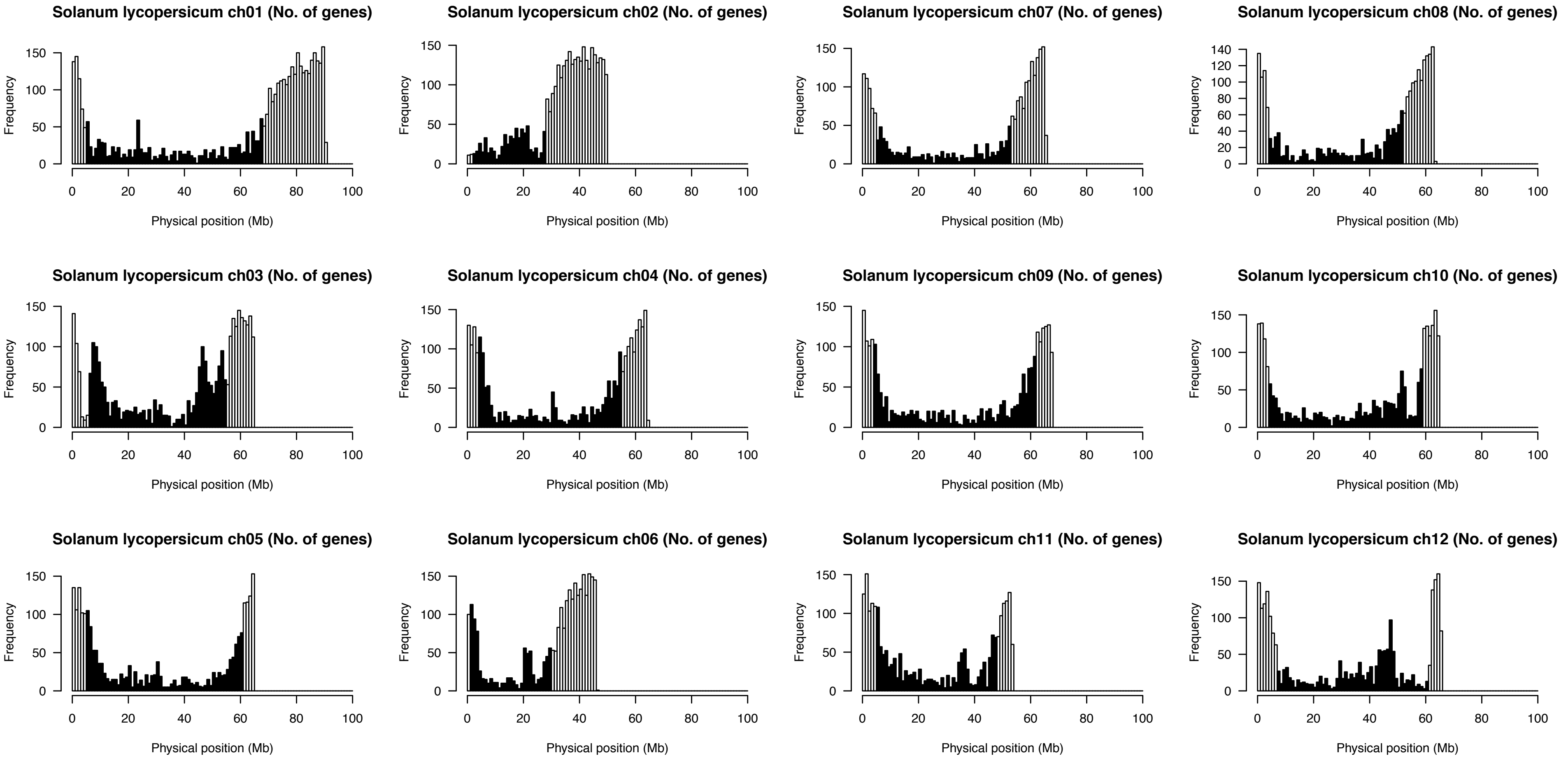


**Supplementary Figure S5** Distributions of the numbers of genes predicted from the tomato genome.
The non-overlapping window size is 1 Mb. Black and white bars indicate hetero- and euchromatic regions.

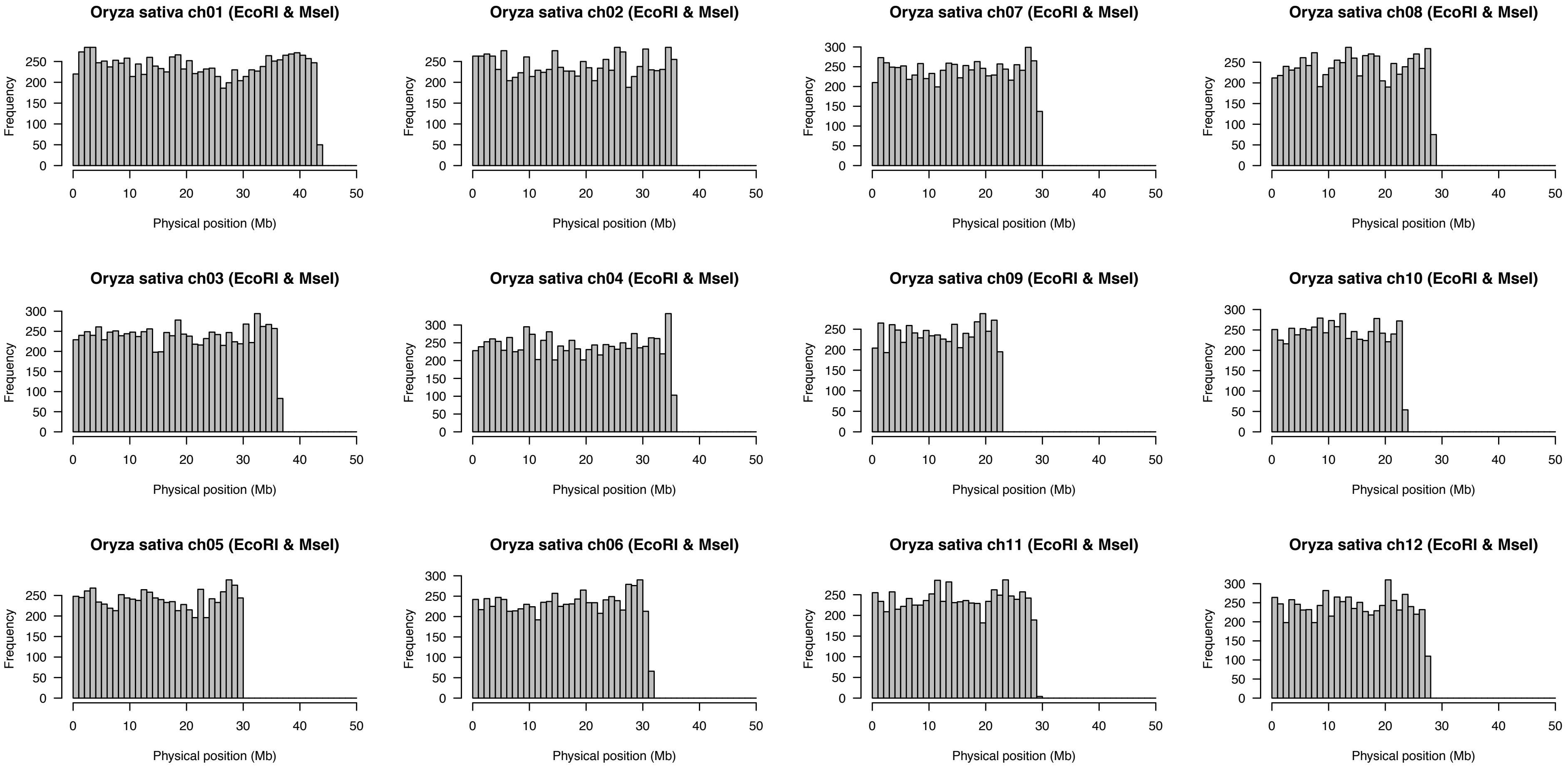


**Supplementary Figure S6** Distributions of the numbers of restriction fragments digested with EcoRI and MseI over the rice genome. The non-overlapping window size is 1 Mb.

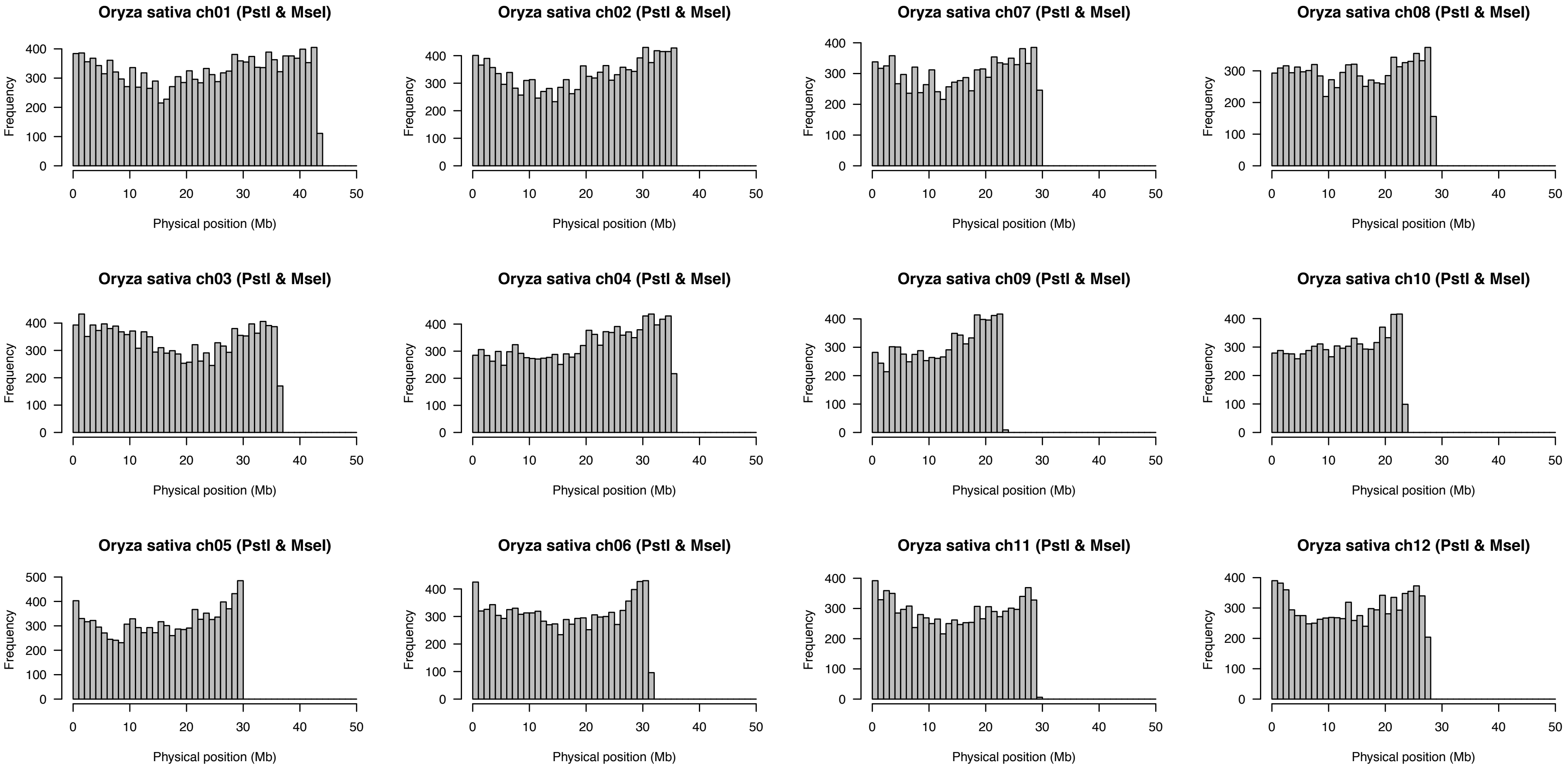


**Supplementary Figure S7** Distributions of the numbers of restriction fragments digested with PstI and MseI over the rice genome. The non-overlapping window size is 1 Mb.

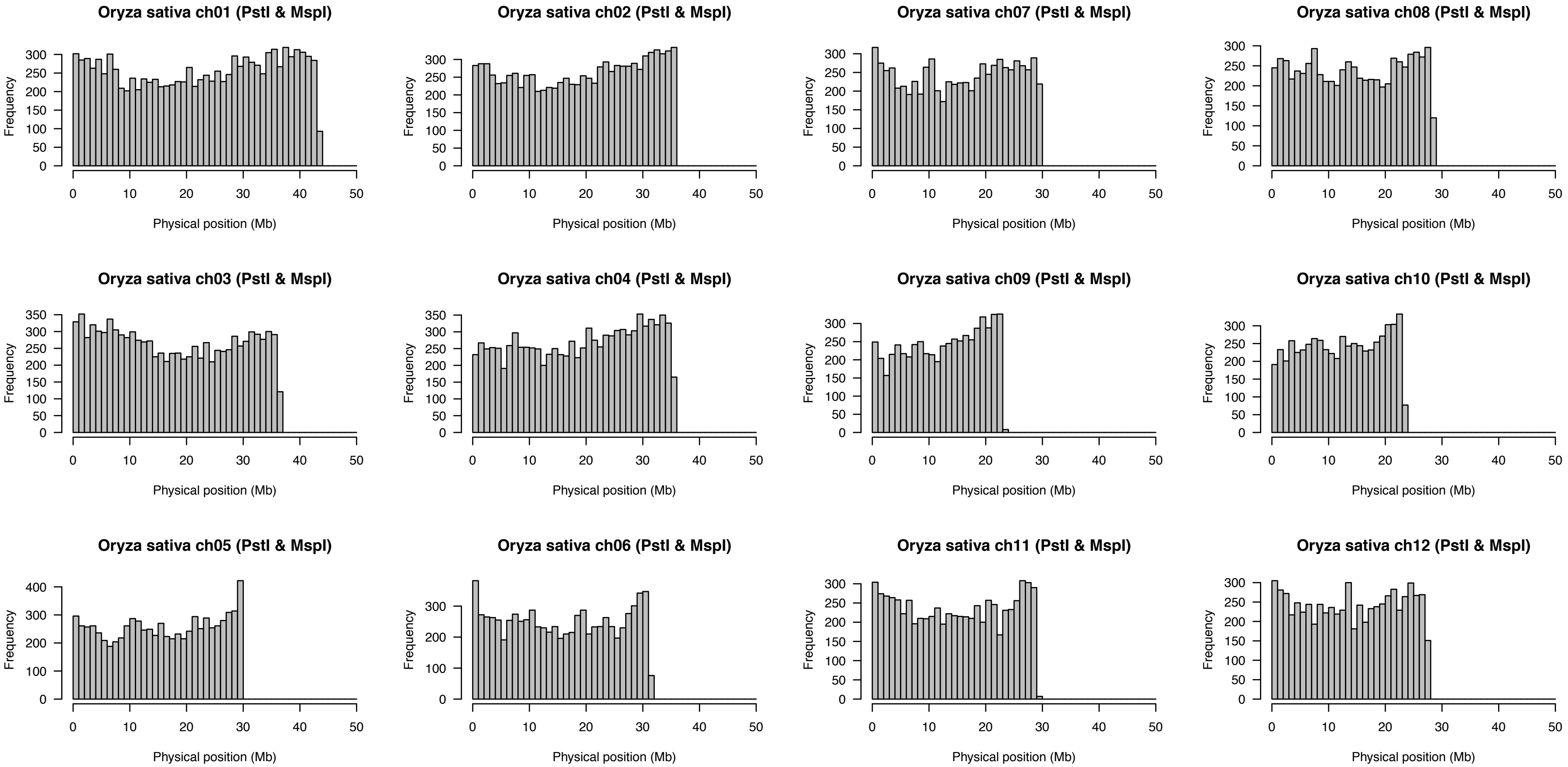


**Supplementary Figure S8** Distributions of the numbers of restriction fragments digested with PstI and MspI over the rice genome. The non-overlapping window size is 1 Mb.

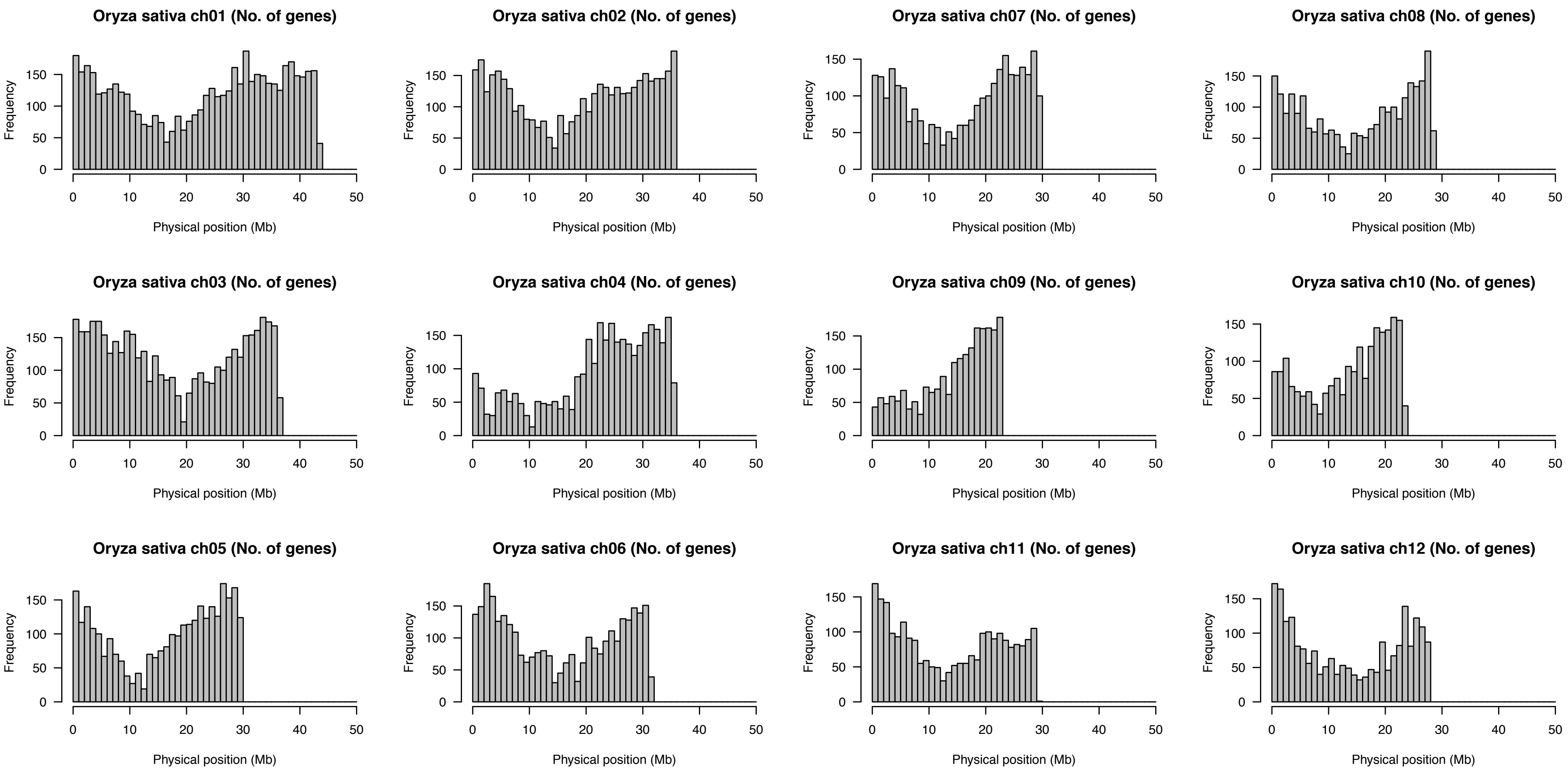


**Supplementary Figure S9** Distributions of the numbers of genes predicted from the rice genome.
The non-overlapping window size is 1 Mb.

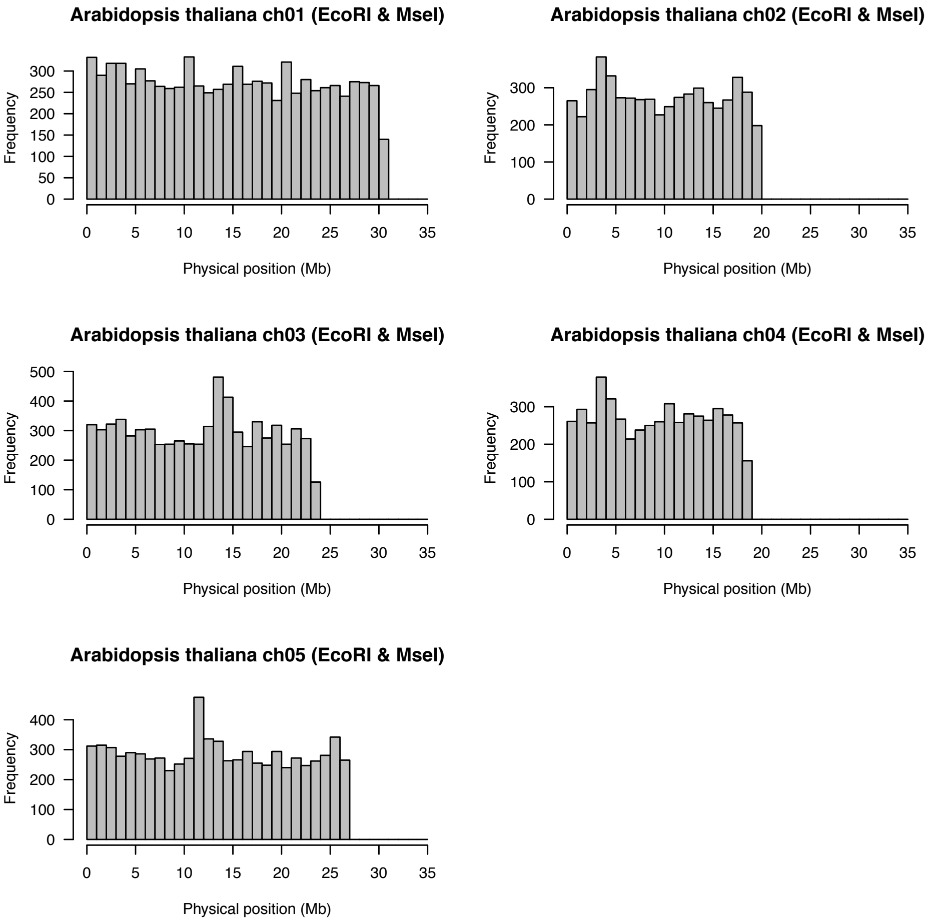


**Supplementary Figure S10** Distributions of the numbers of restriction fragments digested with EcoRI and MseI over the *Arabidopsis* genome. The non-overlapping window size is 1 Mb.

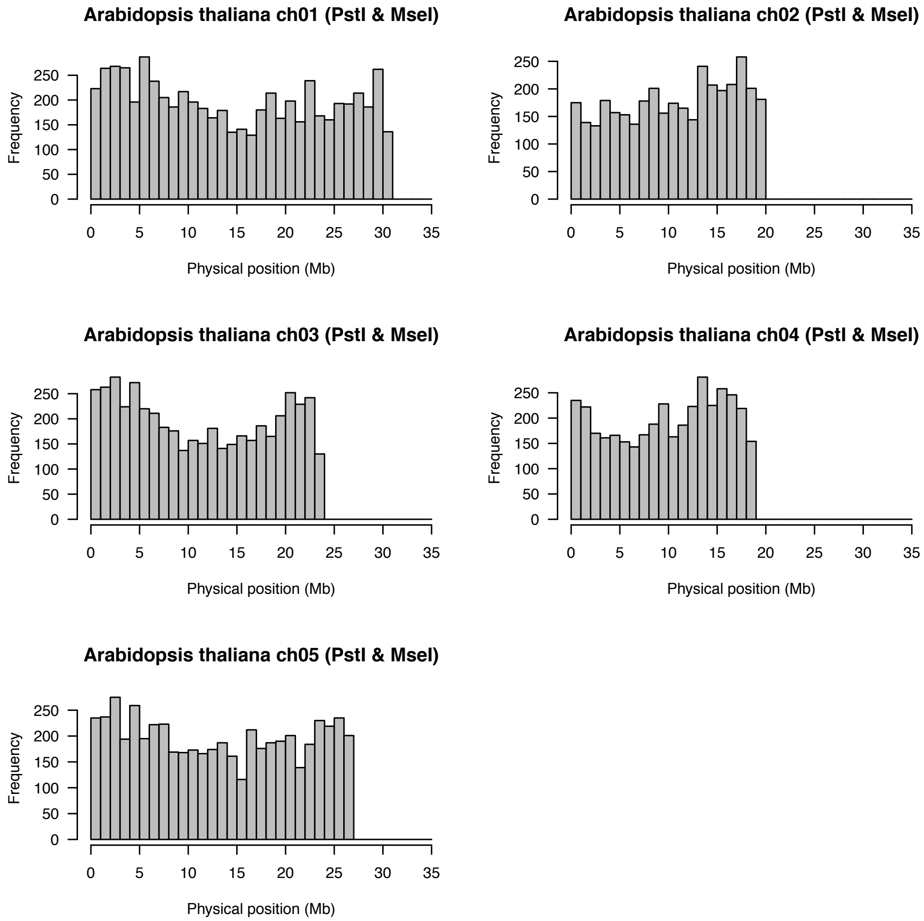


**Supplementary Figure S11** Distributions of the numbers of restriction fragments digested with PstI and MseI over the *Arabidopsis* genome. The non-overlapping window size is 1 Mb.

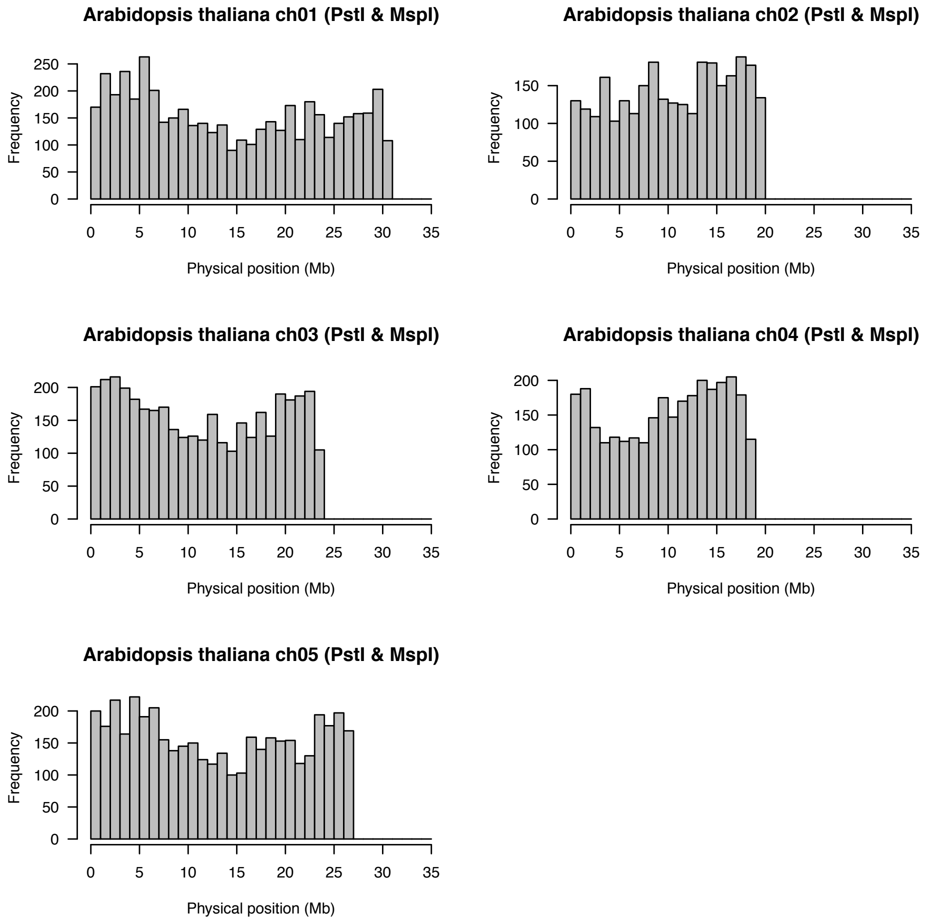


**Supplementary Figure S12** Distributions of the numbers of restriction fragments digested with PstI and MspI over the *Arabidopsis* genome. The non-overlapping window size is 1 Mb.

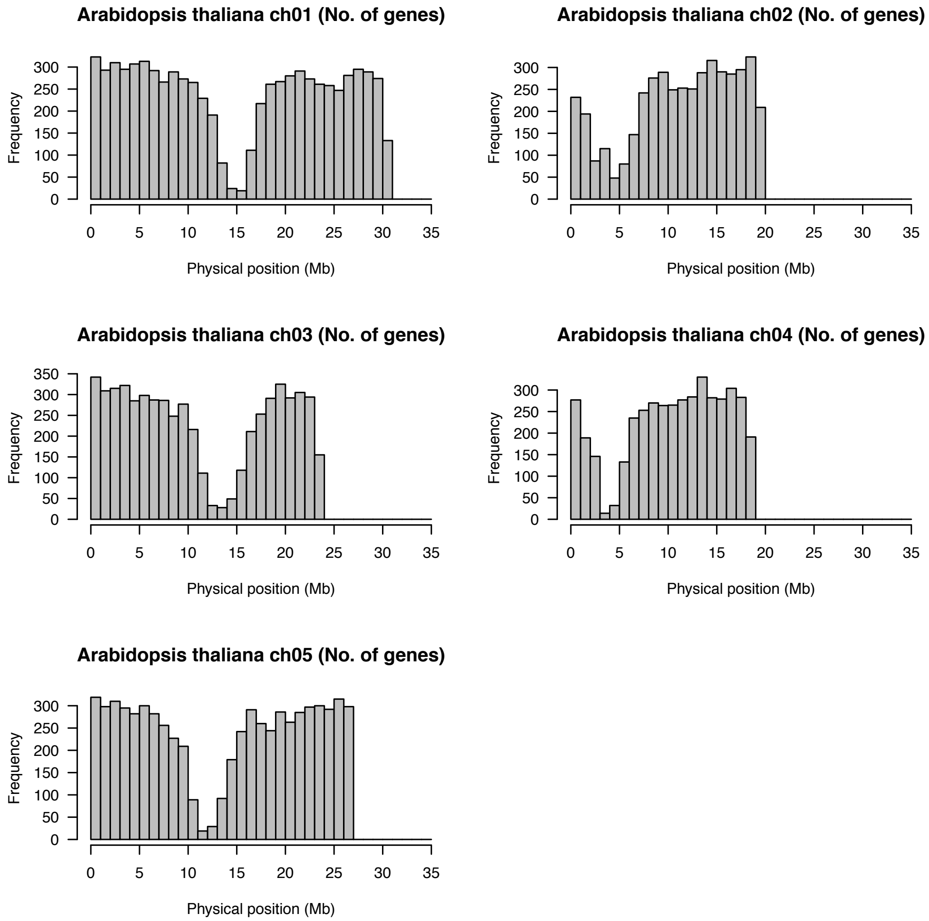


**Supplementary Figure S13** Distributions of the numbers of genes predicted from the *Arabidopsis* genome.
The non-overlapping window size is 1 Mb.

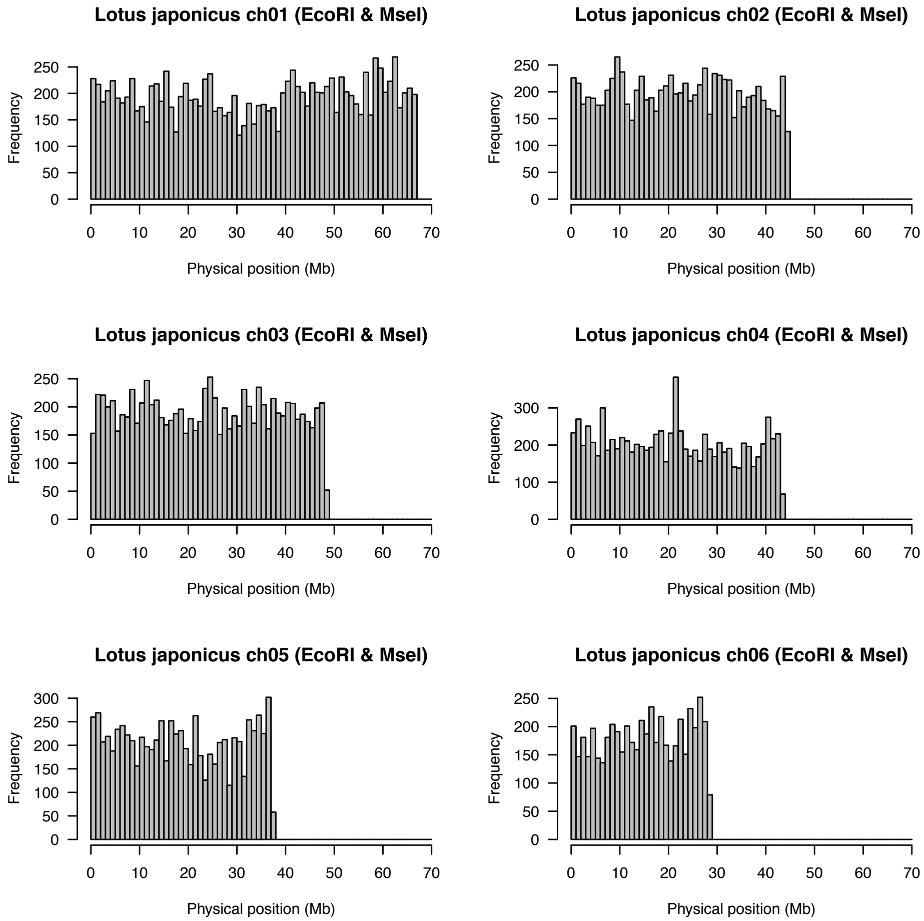


**Supplementary Figure S14** Distributions of the numbers of restriction fragments digested with EcoRI and MseI over the *Lotus* genome. The non-overlapping window size is 1 Mb.

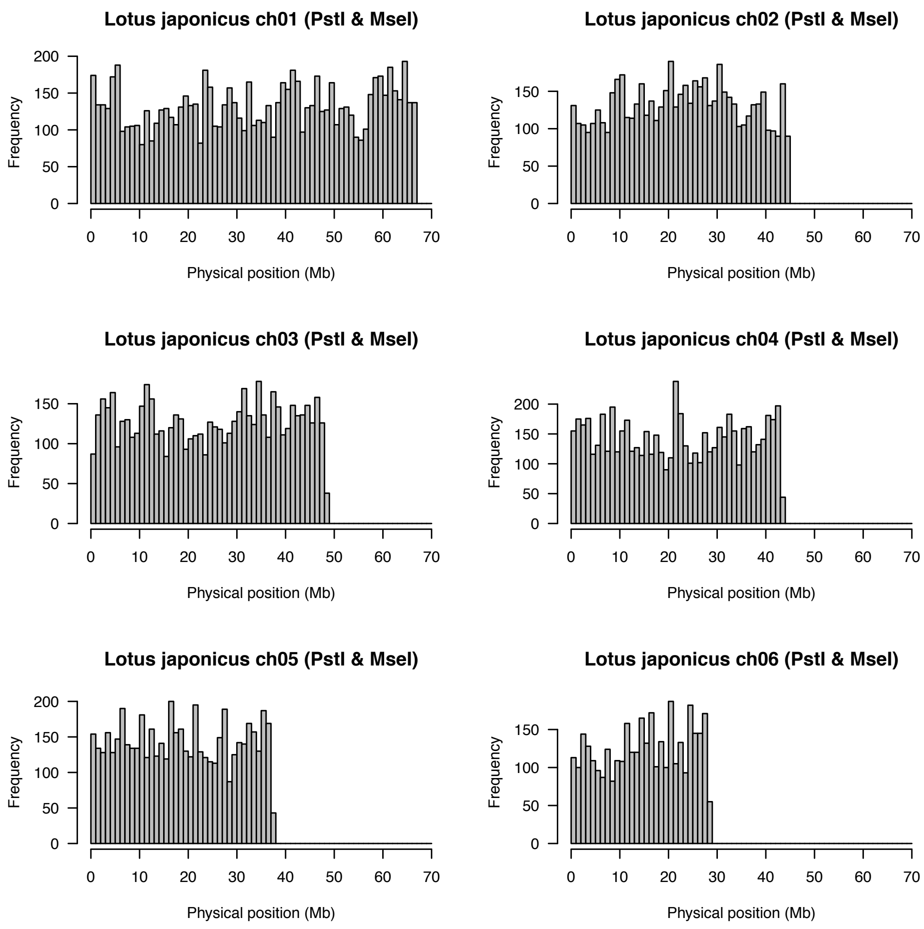


**Supplementary Figure S15** Distributions of the numbers of restriction fragments digested with PstI and MseI over the *Lotus* genome. The non-overlapping window size is 1 Mb.

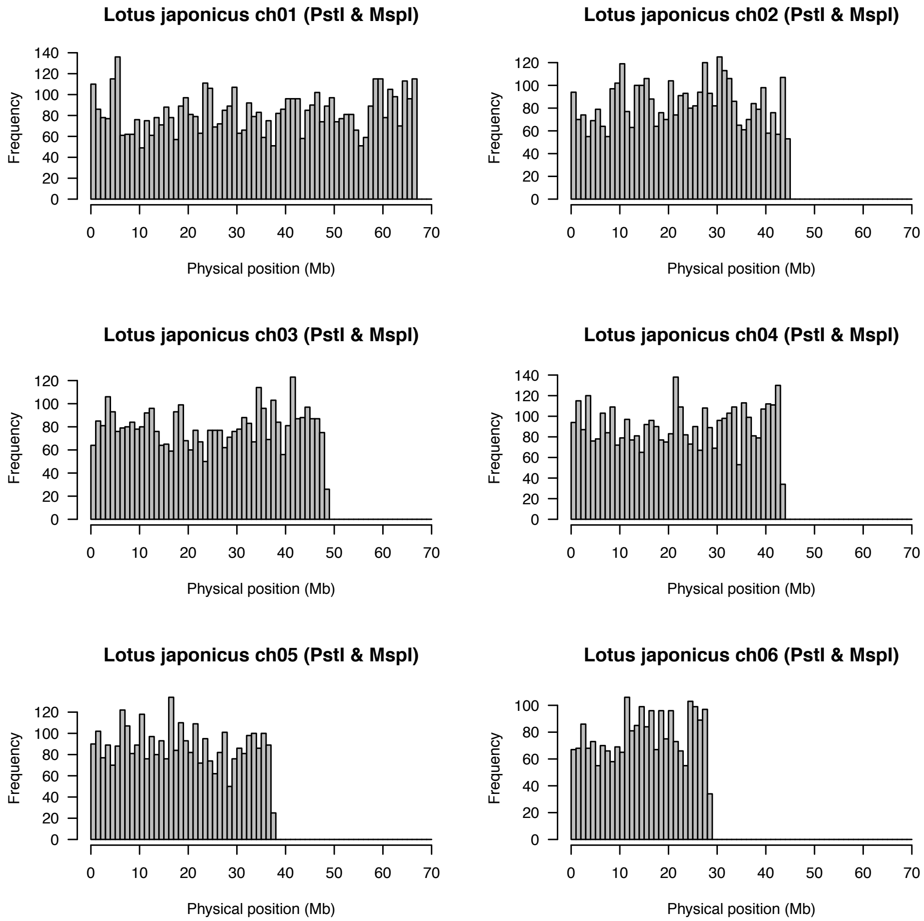


**Supplementary Figure S16** Distributions of the numbers of restriction fragments digested with PstI and MspI over the *Lotus* genome. The non-overlapping window size is 1 Mb.

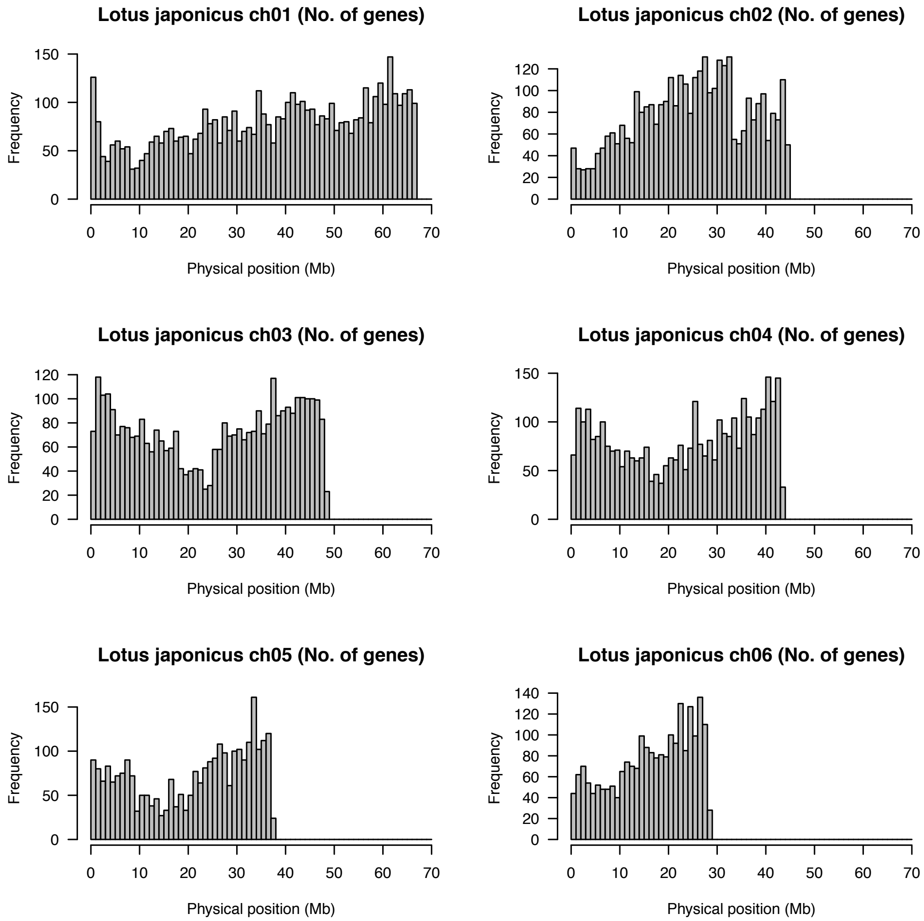


**Supplementary Figure S17** Distributions of the numbers of genes predicted from the *Lotus* genome.
The non-overlapping window size is 1 Mb.
